# Combining climatic and genomic data improves range-wide tree height growth prediction in a forest tree

**DOI:** 10.1101/2020.11.13.382515

**Authors:** Juliette Archambeau, Marta Benito Garzón, Frédéric Barraquand, Marina de Miguel Vega, Christophe Plomion, Santiago C. González-Martínez

## Abstract

Population response functions based on climatic and phenotypic data from common gardens have long been the gold standard for predicting quantitative trait variation in new environments. However, prediction accuracy might be enhanced by incorporating genomic information that captures the neutral and adaptive processes behind intra-population genetic variation. We used five clonal common gardens containing 34 provenances (523 genotypes) of maritime pine (*Pinus pinaster* Aiton) to determine whether models combining climatic and genomic data capture the underlying drivers of height-growth variation, and thus improve predictions at large geographical scales. The plastic component explained most of the height-growth variation, probably resulting from population responses to multiple environmental factors. The genetic component stemmed mainly from climate adaptation, and the distinct demographic and selective histories of the different maritime pine gene pools. Models combining climate-of-origin and gene pool of the provenances, and positive-effect height-associated alleles (PEAs) captured most of the genetic component of height-growth and better predicted new provenances compared to the climate-based population response functions. Regionally-selected PEAs were better predictors than globally-selected PEAs, showing high predictive ability in some environments, even when included alone in the models. These results are therefore promising for the future use of genome-based prediction of quantitative traits.

## 1 Introduction

Global change is expected to have a profound impact on forests (Franklin et al. 2016, Seidl et al. 2017), and whether tree populations will be able to migrate or persist across their current range is uncertain (Aitken et al. 2008). Assessing the potential of populations to accommodate future environmental conditions requires a thorough understanding of the origin of variation in quantitative traits subject to natural selection (Shaw and Etterson 2012, Alberto et al. 2013). To this aim, a necessary first step is to quantify the plastic and genetic components of adaptive traits and their interaction in multiple environments (Des Marais et al. 2013, Merilä and Hendry 2014), which has been done extensively in forest trees (Franks et al. 2014). A second step consists in identifying the underlying drivers of these components (Merilä and Hendry 2014). The plastic component corresponds to the ability of one genotype to produce varying phenotypes depending on the environment (Bradshaw 1965). Phenotypic plasticity can help individuals to overcome new conditions up to a certain threshold (Nicotra et al. 2010), and can be to some extent genetically assimilated and therefore involved in the evolutionary process of adaptation (Pigliucci et al. 2006). The genetic component can stem from both neutral (e.g. population demographic history and genetic drift) and adaptive processes (e.g. adaptation to local biotic and abiotic environments), both processes implying changes in allele frequencies. Populations are locally adapted when they have higher fitness in their own environment than populations from other environments (Kawecki and Ebert 2004). In forest trees, a large amount of work highlighted the importance of climate in driving the plastic and genetic responses of quantitative traits to new environmental conditions (Savolainen et al. 2007, Valladares et al. 2014b). However, it is still unclear how multiple and interacting drivers underlying quantitative trait variation could be combined to improve predictions of population responses to global change. The increasing availability of genomic data opens new opportunities to boost prediction accuracy, which is critical for breeding (i.e. genomic selection; Grattapaglia and Resende 2011), to anticipate future distribution of natural populations (e.g. Razgour et al. 2019), or to support the ongoing development of assisted gene flow strategies aiming to help populations adapt to future environments (Browne et al. 2019, Mahony et al. 2020, MacLachlan et al. 2021).

In forest trees, a long history of common gardens (Langlet 1971) has provided a unique framework to associate population-specific quantitative trait variation with large environmental or geographical gradients, and thus identify populations at risk under climate change (Rehfeldt et al. 1999, 2003, Savolainen et al. 2007, Pedlar and McKenney 2017, Rehfeldt et al. 2018, Fréjaville et al. 2020). The development of population response functions was a step forward to evaluate the relative contribution of plasticity -associated to current climatic conditions (i.e. the climate in the common gardens)- and genetic adaptation -associated to the past climatic conditions under which the populations have evolved (i.e. the climate-of-origin of the provenances tested)-in explaining quantitative trait variation (O’Neill et al. 2008, Wang et al. 2010). These models have now been applied to a large variety of traits (Leites et al. 2012a,b, Benito Garzón et al. 2019, Vizcaíno-Palomar et al. 2020) and one of their main conclusions is that trait variation across species ranges is mostly associated with the climate in the common garden (i.e. related to the plastic component) and, only to a much lesser extent, with the climate-of-origin of the provenances (i.e. related to the genetic component) (Leites et al. 2012b, Benito Garzón et al. 2019). Importantly, these models do not allow to determine to what extent associations between trait variation and provenance climate-of-origin, or the higher trait values of local compared to foreign populations, are caused by adaptive or neutral processes (Leimu and Fischer 2008, Hereford 2009, Franks et al. 2014). This limits our understanding of the genetic processes that led to the current patterns of quantitative trait variation, and therefore our ability to predict trait variation of new (untested in common gardens) populations under new environments.

The advent and generalization of genomic tools have enhanced our understanding of adaptive and neutral genetic processes resulting in trait variation, and their relationship with climatic gradients (Savolainen et al. 2013, Sork 2018, Leroy et al. 2020). Integrating genomic information into quantitative trait prediction would be highly valuable to consider intraspecific variability at a finer scale than in current models (Mahony et al. 2020), thereby probably improving model accuracy, especially for populations not previously planted in commons gardens. More specifically, rapidly growing knowledge on trait-associated alleles identified by Genome-Wide Association Studies (GWAS) is promising for anticipating the genetic response of populations to new environments (Exposito-Alonso et al. 2018, Browne et al. 2019). For example, Mahony et al. (2020) used counts of alleles positively associated with the traits of interest (PEAs) to describe patterns and identify drivers of local adaptation in lodgepole pine. Recent studies have shown that most quantitative traits are highly polygenic (see reviews in Pritchard et al. 2010, Barghi et al. 2020; and de Miguel et al. 2020 for maritime pine) and that the effect of trait-associated alleles may vary across environments (Anderson et al. 2013, Tiffin and Ross-Ibarra 2014), which complicates the use of genomic information in trait prediction. In addition, patterns in allele frequencies induced by population demographic history are often correlated with environmental gradients (Latta 2009, Alberto et al. 2013, Nadeau et al. 2016), which makes difficult to separate the signature of population structure from that of adaptive processes (Sella and N. H. Barton 2019, Sohail et al. 2019). At the species range scale, population structure hinders the use of genomic relationship matrices, which provide more accurate estimates of genetic parameters (e.g. breeding values, additive and non-additive variance) within breeding populations than previously used pedigree-based approaches (Bouvet et al. 2016, El-Dien et al. 2018). Indeed, admixed populations or distinct genetic groups may present different means and variances of their genetic values, which requires new statistical methods to estimate them (e.g. Muff et al. 2019). Thus, integrating genomic information into quantitative trait prediction in natural populations, while highly valuable, remains challenging.

Forest trees are remarkable models to study the genetic and plastic components of quantitative trait variation. Forest tree populations often have large effective population size and are distributed along a large range of environmental conditions, which makes them especially suitable to study current and future responses to climate (Savolainen et al. 2007, Alberto et al. 2013). Moreover, forest trees remain largely undomesticated (including those species with breeding programs) and, therefore, genetic variation in natural populations has been little influenced by human-induced selection (Neale and Savolainen 2004). However, forest trees have also large and complex genomes (especially conifers; Mackay et al. 2012), that show a rapid decay of linkage disequilibrium (Olson et al. 2010), and extensive genotyping would be needed to identify all (most) relevant polymorphisms underlying (highly polygenic) quantitative traits (Neale and Savolainen 2004, Jaramillo-Correa et al. 2015). In addition, although early results have been convincing in predicting trait variation within tree breeding populations (i.e. using populations with relatively low effective population size; Resende Jr et al. 2012, Resende et al. 2012, Jarquín et al. 2014), predicting the genetic component of trait variation across populations or geographical regions of forest trees remains poorly explored.

In the present study, we aim to identify the potential drivers of the plastic and genetic components of height growth in distinct maritime pine gene pools (i.e. genetic clusters) and investigate how common garden data can be combined with genomics to efficiently predict height-growth variation across the species range. We compared Bayesian hierarchical mixed models that inferred height-growth variation in maritime pine as a function of climatic and genomic-related variables, using a clonal common garden network (CLONAPIN) consisting of five sites and 34 provenances (523 genotypes and 12,841 trees). First, we evaluated the relative importance of potential drivers underlying height-growth variation. We expected that: (i) the plastic component explains most trait variation and is associated with climate in the common gardens, (ii) the genetic component is driven by both adaptive processes, such as adaptation to climate, and neutral processes, such as population demographic history. Second, we compared the out-of-sample predictive ability (on unknown observations or provenances) of models based exclusively on the common garden design and models including (either separately or jointly) potential predictors of the genetic component of trait variation, notably those related to climate and positive-effect height-associated alleles (PEAs). We expected that the distinct demographic history of maritime pine gene pools, the provenance climate-of-origin and the counts of PEAs, either combined or alone, may improve height-growth predictions of unknown provenances. We also expected that height-associated alleles selected regionally, i.e. in particular environments, would have a better predictive ability than globally-selected alleles. Our study is a step towards integrating the recent knowledge brought by large genomic datasets to the modeling of quantitative trait variation in forest trees. Combining common gardens with genomic tools hold great promise for speeding up and improving trait predictions at large scales and for a wide range of species and populations. However, a robust framework is needed to make reliable predictions and to determine when and to what extent genomics can help in making decisions in conservation strategies or in anticipating population responses to climate change.

## 2 Materials & Methods

### 2.1 Plant material and phenotypic measurements

Maritime pine (*Pinus pinaster* Ait., Pinaceae) is an economically important forest tree, largely exploited for its wood (Viñas et al. 2016). It has also an important ecological function stabilizing coastal and fossil dunes and as keystone species supporting forest biodiversity. Native to the western part of the Mediterranean Basin, the Atlas mountains in Morocco, and the south-west Atlantic coast of Europe, its natural distribution spans from the High Atlas mountains in the south (Morocco) to French Brittany in the north, and from the coast of Portugal in the west to western Italy in the east. Maritime pine is a wind-pollinated, outcrossing and long-lived tree species that can grow on a wide range of substrates, from sandy and acidic soils to more calcareous ones. It can also withstand many different climates: from the dry climate of the Mediterranean Basin to the highly humid climate of the Atlantic Europe region, and the continental climate of central Spain. Maritime pine populations are highly fragmented and can be grouped into six gene pools (Alberto et al. 2013, Jaramillo-Correa et al. 2015; see fig. 1), that is genetic clusters that cannot be differentiated on the basis of neutral genetic markers and that probably derive from a common glacial refuge (Bucci et al. 2007, Santos-del-Blanco et al. 2012).

**Figure 1.**
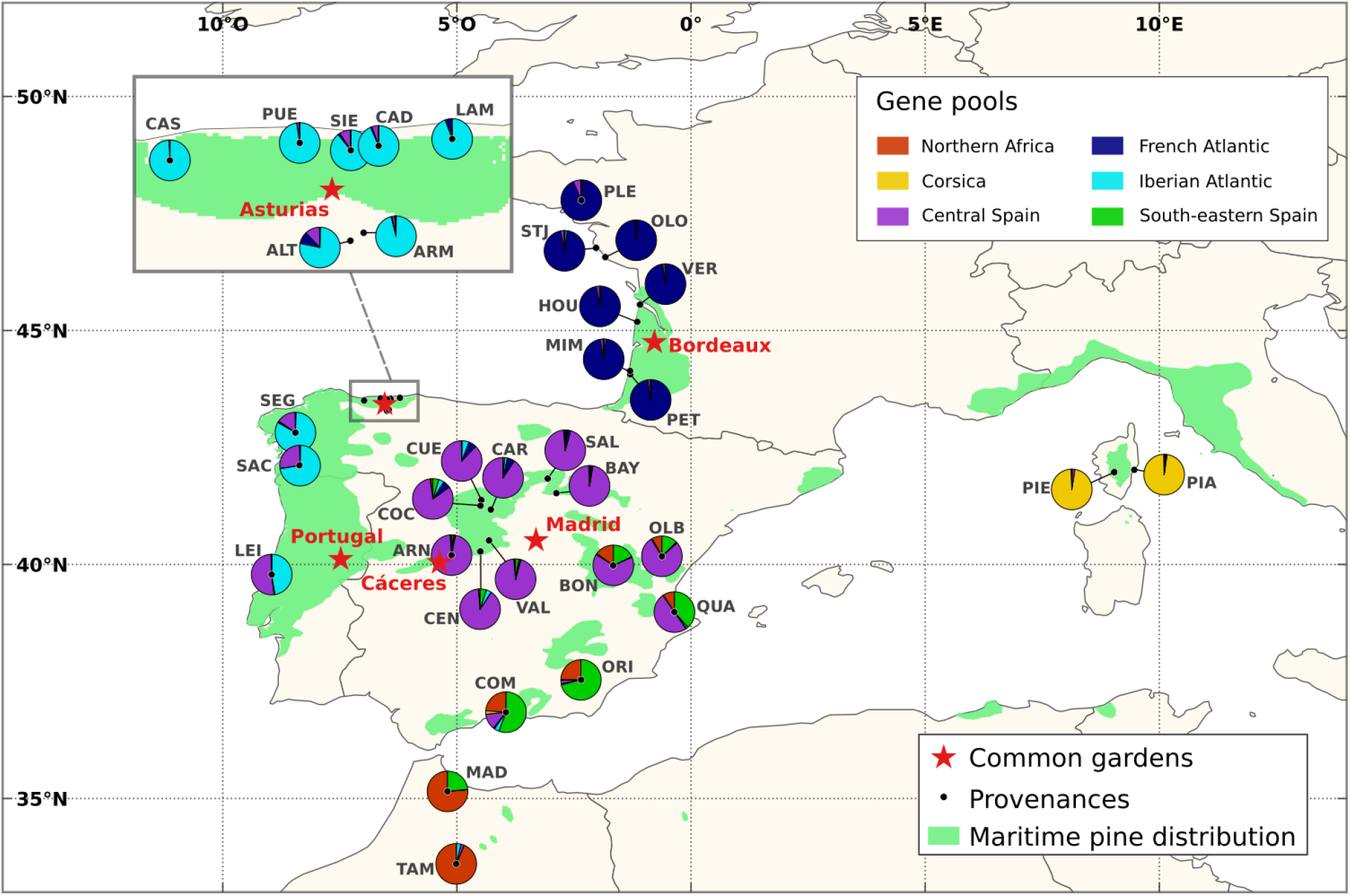
The five common gardens and 34 provenances of maritime pine (CLONAPIN common garden network) used in this study. The distribution of maritime pine is also shown (based on EUFORGEN map, http://www.euforgen.org/). Pie charts represent the proportions belonging to each gene pool for each provenance (see legend) as estimated in Jaramillo-Correa et al. (2015). Provenance names can be found in Table S2.

Height growth is a key adaptive trait in forest trees, including maritime pine. Height can be seen as the end-product of multiple ecophysiological processes that are both genetically regulated and affected by multiple environmental effects (Grattapaglia et al. 2009). As such, taller trees compete more efficiently for light, water and nutrients, and are also more likely to have high fecundity (Rehfeldt et al. 1999, Wu and Ying 2004, Aitken and Bemmels 2015). We obtained height data from the clonal common garden network CLONAPIN, consisting of five common gardens located in different environments (also referred as test sites; fig. 1). Three sites are located in the Atlantic Europe region, with mild winters, high annual rainfall and relatively wet summers: Bordeaux in the French part, and Asturias and Portugal in the Iberian part, the Portugal site experiencing slightly colder winters and half the summer precipitation than the site in Asturias. The two other sites, Cáceres and Madrid, are located in the Mediterranean region with high temperatures and intense summer drought, as well as large precipitation differences between summer and winter. In 2010 or 2011 depending on the test site, clonal replicates from 34 provenances were planted in a randomized complete block design with eight blocks. For each provenance, trees represent between 2 and 28 genotypes (clones), on average about 15 (see Rodríguez-Quilón et al. 2016 for details). Genotypes were originally sampled from natural populations, with enough distance among trees (over 50 m) to avoid sampling related individuals. Depending on the site, height was measured from one to four times, when the trees were between 13 and 41 month old (Table S1). Only survivors were measured for height, which resulted in a strongly unbalanced design as 92% and 75% of the trees died in Cáceres and Madrid, respectively (partly due to the clay soils and a strong summer drought). After removing genotypes for which we had no genomic information, we analyzed 33,121 height observations from 12,841 trees and 523 genotypes (Table S2).

### 2.2 Gene pool assignment and positive-effect alleles (PEAs)

DNA was extracted from leaves collected in the Asturias common garden and genotyped with a 9k Illumina Infinium SNP assay (described in Plomion et al. (2016)), resulting in 5,165 high-quality polymorphic SNPs scored on 523 genotypes. There were on average only 3.3 missing values per genotype (ranging between 0 and 142). For each genotype, the proportion belonging to each gene pool was estimated in Jaramillo-Correa et al. (2015), using nine nuSSRs as well as a subset of the same SNPs as in our study (1,745 SNPs) and the Bayesian approach available in Structure v. 2.3.3 (Pritchard et al. 2000; Table S3). This gene pool assignment aimed at reflecting the neutral genetic structure in maritime pine, which results from population demographic history and genetic drift, but may also arise from different selective histories across gene pools.

Based on the 523 genotypes for which there were both genotypic and phenotypic data, we performed four GWAS following the Bayesian variable selection regression (BVSR) methodology implemented in the piMASS software (Guan and Stephens 2011), correcting for population structure and using the height BLUPs reported in de Miguel et al. (2020), that accounted for site and block effects. First, a global GWAS was performed to identify SNPs that have an association with height at range-wide geographical scales, thus using the combined phenotypic data from the five common gardens. Second, three regional GWAS were performed to identify SNPs that have a local association with height in a particular geographical region *r* (i.e. in a particular environment), thus using separately data from the Iberian Atlantic common gardens (Asturias and Portugal), the French Atlantic common garden (Bordeaux) and the Mediterranean common gardens (Madrid and Cáceres). For each of the four GWAS, we selected the 350 SNPs (∼7% top associations) with the highest absolute Rao-Blackwellized estimates of the posterior effect size, corresponding approximately to the estimated number of SNPs with non-zero effects on height in a previous multi-trait study using the same SNP marker set (de Miguel et al. 2020). These SNPs were used to compute the counts of global and regional positive-effect alleles (gPEAs and rPEAs) for each genotype (see section 2.1 of the Supplementary Information for more details).

### 2.3 Climatic data

In forest trees, large-scale patterns of allele frequencies or quantitative trait variation are known to be associated with climatic variables related to mean temperature and precipitation (e.g. Eckert et al. 2010, McLane et al. 2011, Leites et al. 2019, Fréjaville et al. 2020, Mahony et al. 2020), or episodic climatic conditions, such as summer aridity or maximum temperatures (Rehfeldt et al. 2003, Grivet et al. 2011, McLane et al. 2011, Jaramillo-Correa et al. 2015, Fréjaville et al. 2020). As climate change will cause major changes in temperature and precipitation in the near future, particularly in the Mediterranean basin, there is a need to understand the complex influence of climatic variables on quantitative trait variation. We extracted monthly and yearly climatic data from the EuMedClim database with 1 km resolution (Fréjaville and Benito Garzón 2018). The climatic similarity among test sites was described by a covariance matrix Ω including six variables related to both extreme and average temperature and precipitation in the test sites during the year preceding the measurements, and with at most a correlation coefficient of 0.85 among each other (see section 3.1 in the Supplementary Information for more details). The climatic similarity among provenances was described by a covariance matrix Φ including four variables related to the mean temperature and precipitation in the provenance locations over the period from 1901 to 2009 (i.e. representing the climate under which provenances have evolved), and with at most a correlation coefficient of 0.77 among each other (see section 3.2 in the Supplementary Information for more details).

### 2.4 Hierarchical height-growth models

Twelve height-growth models were compared. We first built two baseline models relying exclusively on the common garden design and aimed at quantifying the relative contribution of the genetic and plastic components of height-growth variation (*models M1* and *M2*; Table 1). Second, we used climatic and genomic data to detect association of height-growth variation with potential underlying drivers related to plasticity, adaptation to climate or gene pool assignment (i.e. a proxy of the population demographic history and genetic drift experienced by the populations), and estimated gene pool-specific total genetic variances (*models M3* to *M6*; Table 1). Third, we built models either including separately or combining potential drivers of the genetic component of height-growth variation to predict unknown observations and provenances without relying on the common garden design (*models M7* to *M12*; Table 1). In every model, the logarithm of height (log(*h*)) was used as a response variable to stabilize the variance. Tree age at the time of measurement *i* was included as a covariate to account for the average height-growth trajectory. This implies that all models shared the form log(*h*_*i*_) = *f* (age_*i*_) + *m*(covariates), where *m*(covariates) is the rest of the model. Therefore, all models can also be written *h*_*i*_ = exp(*f* (age_*i*_)) exp(*m*(covariates)), which explains why covariates in our models affect height growth (i.e. modulate the height-growth trajectory) rather than simply height. We used a second-degree polynomial to account for tree age 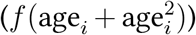 because the logarithm of height first increases linearly with age and then reaches a threshold (fig. S11). Each tree was measured between one and four times (14% of the trees were measured only once), but we did not include a varying intercept for each tree as it resulted in model miss-specification warnings and strong overfitting. A description of each model specification follows.

**Table 1:**
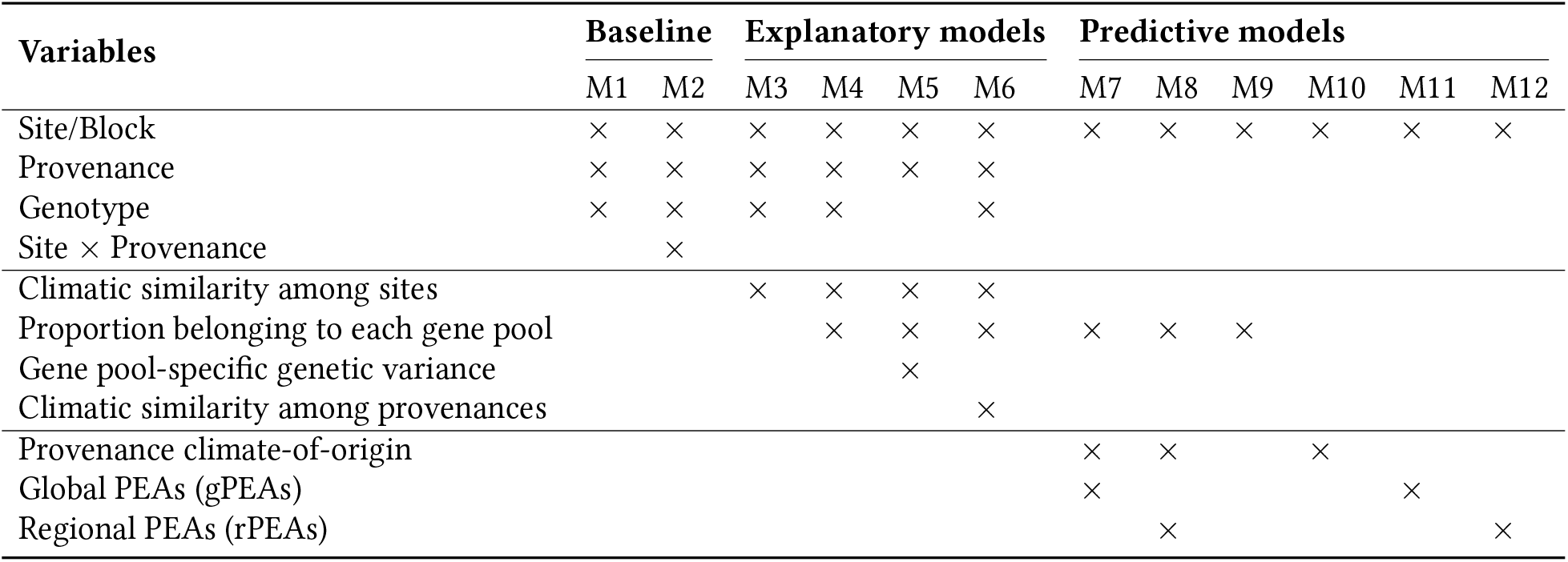
Variables included in the height-growth models. Baseline *models M1* and *M2* separate the genetic and plastic components of height-growth variation via varying intercepts relying exclusively on the common garden design. Explanatory models (*models M3* to *M6*) test different hypotheses regarding the potential drivers underlying height-growth variation. Predictive models (*models M7* to *M12*) are used to compare the predictions on new observations and provenances when combining or including separately genomic and climatic drivers of height-growth variation. The provenance climate-of-origin is evaluated using the precipitation of the driest month, *min*.*pre*, and the maximum temperature of the warmest month, *max*.*temp*. gPEAs and rPEAs correspond to the counts of height-associated positive-effect alleles, selected either globally (across all common gardens) or regionally (in specific common gardens). The provenance climate-of-origin and the PEAs were included in the predictive models with site-specific slopes. All models also contained the age effect, not shown in the table.

#### 2.4.1 Baseline *models M1* and *M2*: separating the genetic and plastic components of height-growth variation

In the baseline *model M1*, height *h* was modeled as a function of tree age, varying intercepts for the sites *S*_*s*_ and blocks nested within sites *B*_*b*(*s*)_ (i.e. the plastic component), and varying intercepts for the provenances *P*_*p*_ and genotypes within provenances *G*_*g*(*p*)_ (i.e. the genetic component):

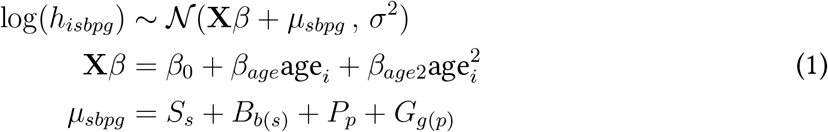

where **X** is the 3-column design matrix and *β* is a vector including the intercept *β*_0_ and the coefficients *β*_*age*_ and *β*_*age*2_ of the fixed effect variables (*age* and *age*^2^, respectively). *µ*_*sbpg*_ is the vector of varying intercepts. *Model M2* was based on *model M1* but including an interaction term between provenance and site (*S*_*s*_*P*_*p*_). We also performed a model without the genetic component (called *M0*) whose outputs are reported in the Supplementary Information.

#### 2.4.2 Explanatory *models M3* to *M6*: potential drivers underlying height-growth variation

In *model M3*, we hypothesized that the plastic component of height growth was influenced by the climatic similarity among test sites during the year preceding the measurements. This model can be expressed with the same likelihood as *M1* but with the vector of varying intercepts equal to:

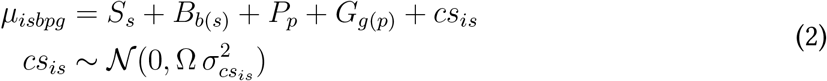

where Ω is the covariance matrix describing the climatic similarity between test sites *s* during the year *i* preceding the measurements (fig. S6) and *cs*_*is*_ are varying intercepts associated with the climatic conditions in each test site *s* during the year *i*. In *M3*, the plastic component was partitioned between the regression on the climatic covariates (*cs*_*is*_) and the deviations related to block and site effects due to the local environmental conditions that are not accounted for by the selected climatic covariates.

In *models M4, M5* and *M6*, we investigated the drivers of the genetic component of height growth. In *M4*, we hypothesized that the genetic component was influenced by the proportion belonging to each gene pool *j. M5* extends *M4* by estimating different total genetic variances in each gene pool while accounting for admixture among gene pools, following Muff et al. (2019). Equations for *M4* and *M5* can be found in section 4 of the Supplementary Information. In *M6*, we hypothesized that populations are genetically adapted to the climatic conditions in which they evolved. Thus, we quantified the association between height growth and the climatic similarity among provenances, while still accounting for the gene pool assignment, such as:

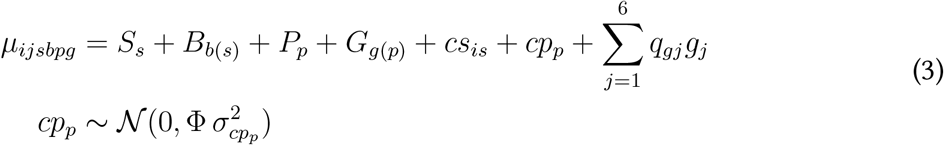

where *q*_*gj*_ corresponds to the proportion belonging of each genotype *g* to the gene pool *j, g*_*j*_ is the mean relative contribution of gene pool *j* to height growth, Φ is the covariance matrix describing the climatic similarity between provenances *p* (fig. S9) and *cp*_*p*_ are varying intercepts associated with the climate in each provenance *p*. Therefore, in *M6*, the genetic component was partitioned among the regression on the climatic covariates (*cp*_*p*_), the gene pool covariates (*g*_*j*_), and the deviations related to the genotype (*G*_*g*(*p)*_) and provenance (*P*_*p*_) effects (resulting, for example, from adaptation to environmental variables not measured in our study).

#### 2.4.3 Predictive *models M7* to *M12*: combining climatic and genomic information to improve predictions

In this last set of models, we replaced the provenance and genotype intercepts by different potential drivers of height-growth variation that do not rely directly on the common garden design, namely the gene pool assignment (as in *M4*), two variables describing the climate in the provenance locations (*min*.*pre* the precipitation of the driest month and *max*.*temp* the maximum temperature of the warmest month) and either global or regional PEAs. This allowed us to determine whether these potential drivers were able to predict the height-growth genetic component as accurately as the provenance and genotype intercepts (i.e. the variables relying directly on the common garden design). In *models M7* and *M8*, the potential predictors were all included together in the models to quantify their predictive performance conditionally to the other predictors, and were expressed as follows (here for *M7*):

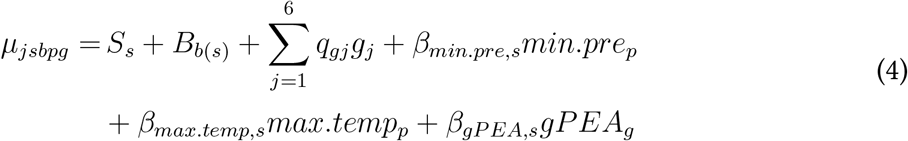

where *min*.*pre*_*p*_ and *max*.*temp*_*p*_ are the climatic variables in the provenance locations, *β*_*min*.*pre,s*_ and *β*_*max*.*temp,s*_ their site-specific slopes, *gPEA*_*g*_ the counts of global PEAs and *β*_*gPEA,s*_ its site-specific slopes. *M8* is identical to *M7*, except that the counts of gPEAs were replaced by counts of rPEAs (i.e. regionally-selected alleles, with positive effects in specific geographical regions/environments). We also performed models in which the potential predictors were included individually to determine their specific predictive performance: the gene pool assignment in *M9*, the provenance climate-of-origin in *M10* and the counts of gPEAs and rPEAs, in *M11* and *M12*, respectively.

All models were inferred in a Bayesian framework as this approach better handles unbalanced and multilevel designs (Clark 2005) and also to better propagate sources of uncertainty from data and parameter values into the estimates (de Villemereuil 2019). Priors used in the models were weakly informative and are provided in section 4.2 of the Supplementary Information. To build the models, we used the *brms* package (Bürkner 2017), based on the no-U-turn sampler algorithm. Models were run with four chains and between 2,000 and 3,000 iterations per chain depending on the models (including 1,000 warm-up samples not used for the inference). All analyses were undertaken in R version 3.6.3 (R Core Team 2020) and scripts are available at https://github.com/JulietteArchambeau/HeightPinpinClonapin.

### 2.5 Comparing model goodness-of-fit and predictive ability

Three partitions of the data (P1, P2 and P3) were used to evaluate model goodness-of-fit (i.e. in-sample explanatory power, using training datasets) and predictive ability (out-of-sample predictive power, using test datasets). In P1, we aimed to predict new observations, an observation being a height-growth measurement in a given year on one individual. P1 corresponds to a random split of the data between 75% of observations used to fit the models (the training dataset of 24,840 observations) and 25% of observations used to evaluate model predictions (the test dataset of 8,281 observations). Notice that the test dataset of the P1 partition was not totally independent from the training dataset as it belongs to the same genotypes/provenances and blocks/sites. In P2 and P3, we aimed to predict new provenances. P2 corresponds to a random split between a training dataset of 28 provenances and a test dataset containing the remaining 6 provenances. P3 corresponds to a non-random split between a training dataset of 28 provenances and a test dataset containing 6 provenances with at least one provenance from each under-represented gene pool (i.e. northern Africa, south-eastern Spain and Corsican gene pools; see section 6.3 of the Supplementary Information for details). Therefore, the test datasets of the P2 and P3 partitions represent fully independent sets of provenances.

To evaluate the model goodness-of-fit, we calculated the in-sample (in the training dataset) proportion of the variance explained by each model *m* in each common garden *s*, conditional on the age effect, such as: 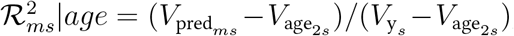, where 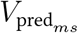 is the variance of the modeled predictive means from model *m* in site *s*, 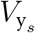 the phenotypic variance in the site *s* and 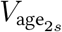 the variance explained by the age effect in the model *M2* in site *s*. We used 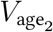 of model *M2* and not of model *m* because the variance predicted by the different fixed effects of some of the models (*M7* to *M12*) could not be properly separated. Moreover, as *M2* is the model with the highest predictive ability among the models relying only on the common garden design (Table S4), it constitutes an adequate baseline for model comparison. In addition, for baseline models *M1* and *M2*, we also calculated the in-sample proportion of the variance explained by the different model components (i.e. genetic, environment and genetic × environment) conditional on the age effect, e.g. for the genetic component in *M*1: 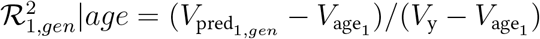 where 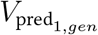 is the variance explained by the genetic component (including the provenance and clone effects) in *M1, V*_y_ the phenotypic variance and 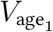 the variance explained by the age effect in *M1*.

Finally, to evaluate the model predictive ability, we calculated the out-of-sample (in the test dataset) proportion of the variance predicted by each model *m* in each common garden *s* conditional on the age effect, that we called *prediction* 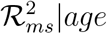. Details about calculating *prediction* 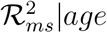 and some supplementary indexes used for model comparison are presented in section 5 of the Supplementary Information.

## 3 Results

### 3.1 Underlying drivers of height-growth variation

In this part, we disentangled the different components of height-growth variation and provided insights on their underlying drivers. Baseline and explanatory models (i.e. *models M1* to *M6*) explained ∼81.5% of height-growth variation, including 57% due to the age effect (Table S4). Based on *M1*, ∼47% (45-48% CIs) of the variation that was not explained by the age effect (i.e. deviating from the growth trajectory) came from the plastic component, ∼11% (11-12% CIs) from the genetic component and ∼43% (42-44% CIs) remained unexplained (fig. 2A & Table S5). In *M2* (same model as *M1* but adding the provenance-by-site interaction), the proportion of variance explained by the provenance-by-site interaction was not different from zero (Table S5). Therefore, we mostly interpret parameter estimates of *M1* (fig. 3), whose results are very similar to *M2*, but with smaller credible intervals (Tables S15 & S18). The plastic component was largely driven by the variance among sites 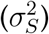, with very little contribution of the variance among blocks 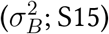. Trees grew the least in Madrid and the most in Asturias (fig. 3 & Table S16). The genetic component was equally attributed to the variance among provenances 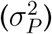 and genotypes (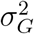; Table S15), with the average height of the provenances appearing to be influenced by their belonging to particular gene pools (fig. 3; and more details in section 6.1.1 of the Supplementary Information).

**Figure 2.**
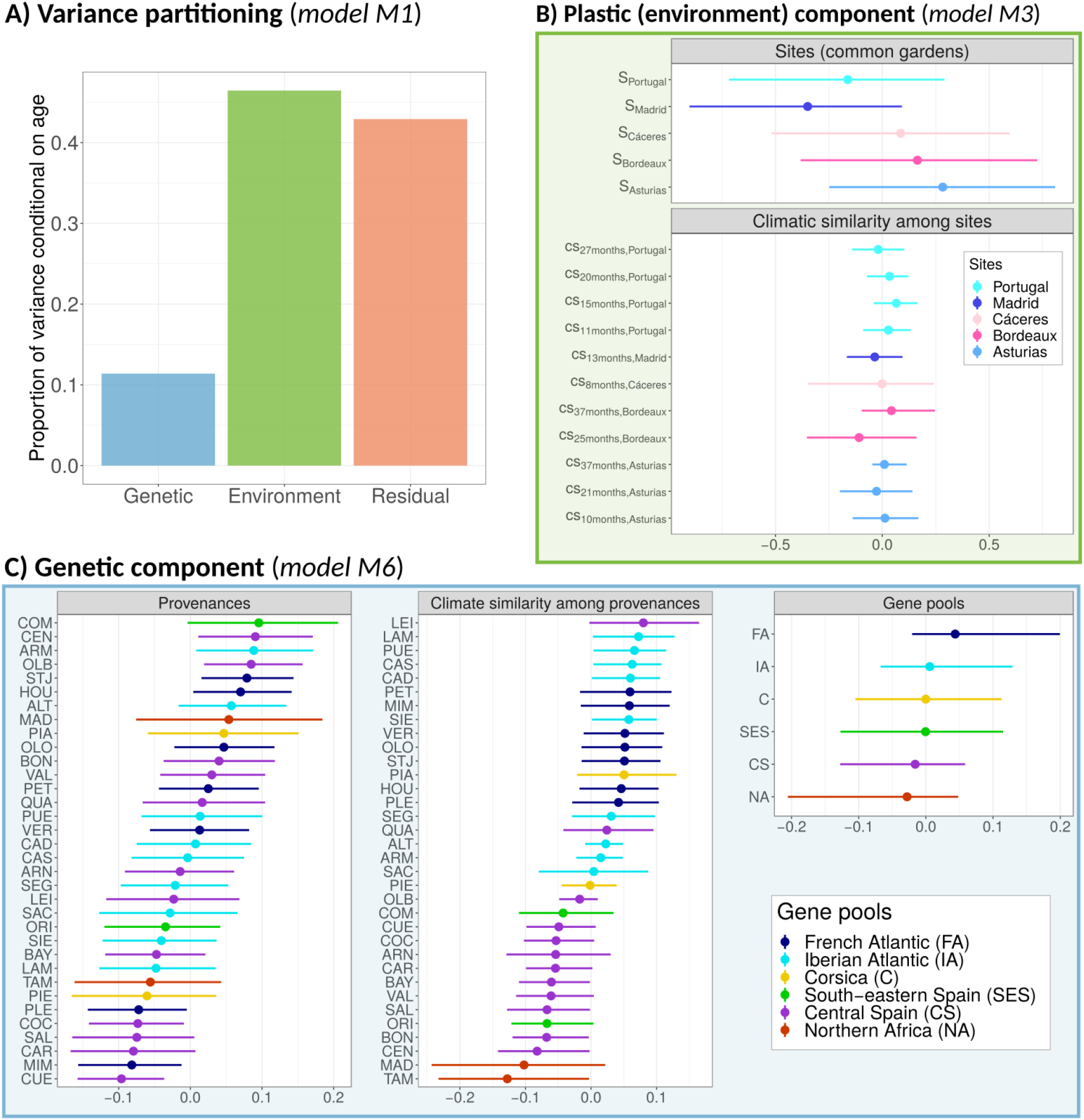
Understanding the genetic and plastic bases of height-growth variation and their potential underlying drivers. A) shows the variance partitioning conditional on age from *model M1* in the P1 partition. B) displays the partitioning of the plastic (i.e. environment) component in *model M3* among the intercepts of the sites (common gardens) (*S*_*s*_) and the intercepts associated with the climatic similarity among sites during the year preceding the measurements (*cs*_*is*_). C) displays the partitioning of the genetic component in model *M6* among the intercepts of the provenances (*P*_*p*_), the intercepts associated with the climatic similarity among provenances (*cp*_*p*_) and the intercepts of the the gene pools (*g*_*j*_). The median and 0.95 credible intervals shown in B) and C) were obtained by fitting the *models M3* and *M6* on the P1 partition. Provenance names can be found in Table S2.

**Figure 3.**
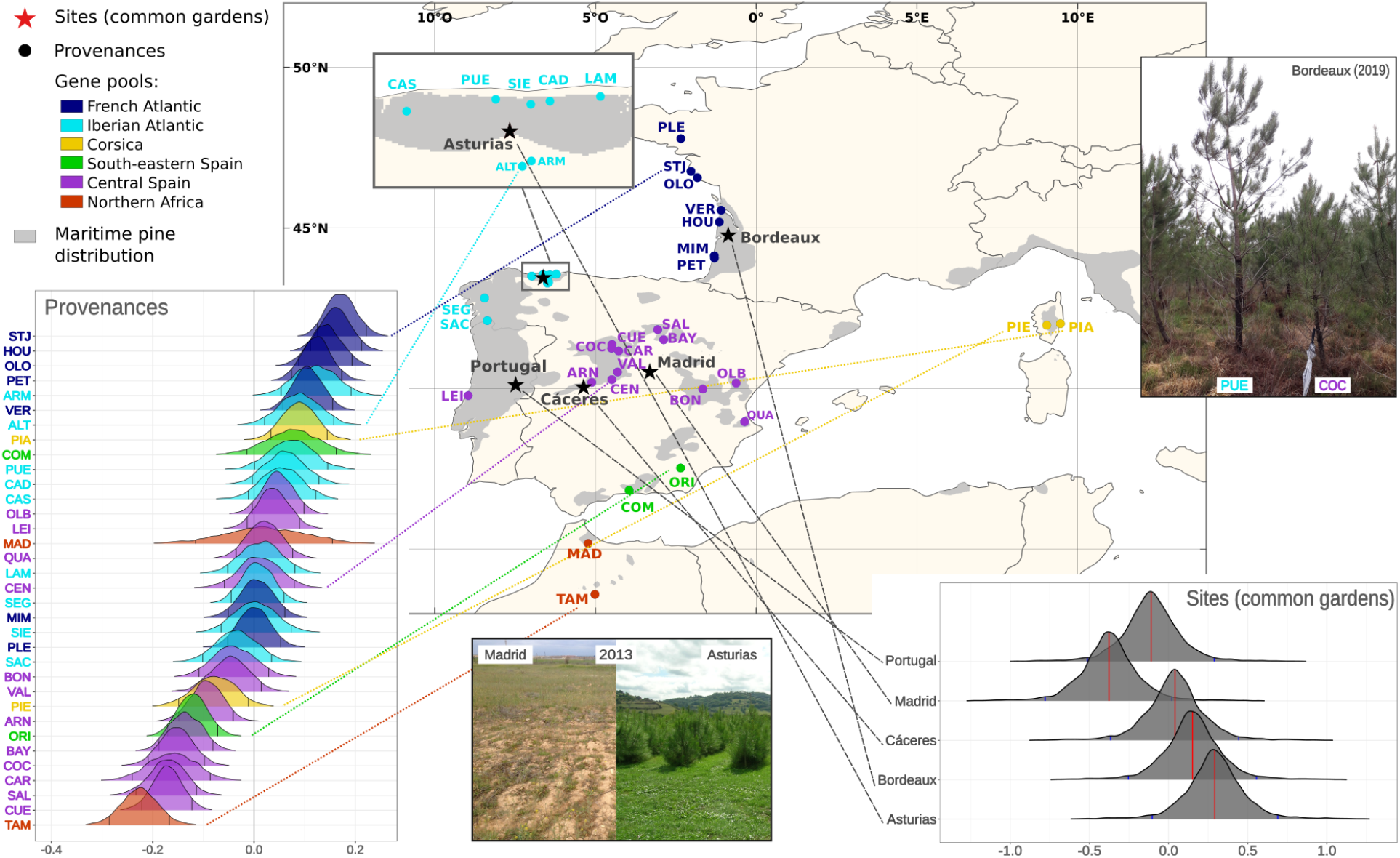
Posterior distributions of the site and provenance intercepts (*S*_*s*_ and *P*_*p*_) in *model M1* on a map representation. Provenances are colored according to the main gene pool they belong to. The exact values of the median, standard deviation and 0.95 credible interval of the posterior distributions of the site and provenance intercepts are shown in Tables S16 and S17, respectively. The top right picture shows the height difference in 2019 between one tree from Coca in central Spain (COC) and another from Puerto de Vega in the Iberian Atlantic region (PUE) growing next to each other in the Bordeaux common garden. The bottom picture shows the height difference between the trees growing in Madrid and Asturias, under highly contrasted environments, three years after plantation (2013). Provenance names can be found in Table S2.

Based on *M3*, the plastic component of height-growth came only marginally from the variance associated with climate similarity among sites, which was more than five times lower than the variance associated with site intercepts (fig. 2B & Table S19). However, *M3* may be unable to separate the effect of these two components (see section 6.1.2 in the Supplementary Information). Indeed, when estimating the effect of the climate similarity among sites in a model that did not include varying intercepts for the sites, we found that height growth was positively associated with the climatic conditions in Bordeaux and Asturias, and negatively with those in Madrid and Cáceres, the two Mediterranean sites, and to a lesser extent also in Portugal (Table S24).

Based on *M6*, the genetic component of height growth was mostly determined by the climatic similarity among provenances and to a lesser extent by the gene pool assignment (fig. 2C & Table S29). However, the effects of the gene pools and climatic similarity among provenances were partially confounded, so that the association between height growth and the gene pools was stronger when the climatic similarity among provenances was not included in the models (i.e. *model M4*; Table S25). Populations from climatic regions neighboring the Atlantic Ocean, and mainly belonging to the French and Iberian Atlantic gene pools, were generally the tallest (e.g. CAD, SIE, PUE, LAM and CAS in northwestern Spain; all provenances along the French Atlantic coast; Figs. 2 & 3). Interestingly, the Leiria (LEI) provenance, which has a strong Iberian Atlantic component (Table S3) and had the highest climate intercept estimate (similar to that of the French Atlantic provenances; fig. 2C), was not among the tallest provenances (fig. 3), probably due to its mixed ancestry with the central Spain gene pool (Table S3). Also, the Corsican provenances showed contrasted climate intercepts (fig. 2), with a positive influence on height growth for Pinia (PIA) but not for Pineta (PIE), located under more Mediterranean conditions, which could explain their large differences in height growth (fig. 3). Finally, the four provenances from south-eastern Spain and northern Africa gene pools, under harsh Mediterranean climates, showed all negative climate intercepts (fig. 2). Noticeably, the total genetic variance of the Iberian and French Atlantic gene pools were likely to be lower than that of the Corsican and south-eastern Spain gene pools, and to a lesser extent the central Spain gene pool, thus resulting in gene pool-specific heritabilities (*model M5*; Table S28 and fig. S13A).

### 3.2 Improved prediction of new observations and provenances by combining climatic and genomic data

In this part, we compared the baseline model *M2* (relying exclusively on the common garden design) to the predictive models that either combine genomic and climatic drivers of height-growth variation (i.e. *models M7* and *M8*) or include each driver separately (i.e. *models M9* to *M12*). Models combining genomic and climatic data generally explained in-sample variation almost as well as *M2*, and sometimes even better; e.g. *model M8* (which includes regional PEAs, rPEAs) in the Mediterranean sites (Madrid and Cáceres) (fig. S10). Models including each driver of height-growth variation separately had a lower goodness-of-fit (for all common gardens) than both *M2* and the models combining the genomic and climatic data, except for *M12* (the model including only rPEAs), which explained in-sample variation almost as well as *M2* and even better than *M7* in Madrid (fig. S10).

Model differences in their predictive ability on new observations (observations not used to fit the models; test dataset of the P1 partition) showed similar patterns than for the goodness-of-fit (Table 4), which was expected as the new observations were sampled among the same provenances and genotypes. However, importantly, models combining genomic and climatic data provided much better predictions of height-growth on new provenances (provenances not used to fit the models; test datasets of the P2 and P3 partitions) than did *M2*, with *M8* having a better predictive ability than *M7* in the Mediterranean sites in the P2 partition and in the Atlantic sites in the P3 partition (Table 4). Models including each driver of height-growth variation separately had also a higher predictive ability on new provenances than *M2*, albeit lower than models combining genomic and climatic data, except *model M12* that showed a higher predictive ability than *M7* in the Mediterranean sites in the P2 partition (Table 4). In *model M12*, one standard deviation increase in rPEAs was associated, on average, with 19.0% increase in height in Madrid, 12.7% in Cáceres, 13.0% in Portugal, 10.4% in Asturias and 9.6% in Bordeaux (section 6.4 of the Supplementary Information). More details on model comparisons are given in section 5 of the Supplementary Information.

## 4 Discussion

We combined genomic, climatic and phenotypic data from five common gardens and 34 provenances of maritime pine (over 30,000 observations) to predict range-wide variation in height growth, a key adaptive trait in forest trees. The plastic component explained the largest part of the deviation from the mean height-growth trajectory (∼47%), probably due to multiple (confounded) environmental factors, including climate. The genetic component explained ∼11% of the deviation from the mean height-growth trajectory and was mainly associated with the provenance climate-of-origin (a proxy of adaptation to climate), whose effect was partially confounded with the proportion belonging to distinct gene pools (a proxy for population demographic history and genetic drift, probably reflecting also the different selective histories of the gene pools). Importantly, we showed that models combining climatic drivers of adaptation, gene pool assignment and counts of height-associated positive-effect alleles (PEAs) captured well the genetic component underlying height-growth variation. They also better predicted height growth of new provenances than models relying exclusively on the common garden design or models including separately climatic and genomic information (e.g. the widely used climate-based population response functions). Interestingly, PEAs that show a regional association with height growth (rPEAs) had a higher predictive ability than PEAs identified globally across the species range (gPEAs). These results pave the way towards integrating genomics into large-scale predictive models of quantitative trait variation.

### 4.1 Predominant role of height-growth plasticity

Plants are known for their remarkable phenotypic plasticity to changing environments (Bradshaw 1965). In long-lived forest trees, the plastic component of quantitative trait variation estimated based on the common garden design is generally higher than the genetic component (Franks et al. 2014, Benito Garzón et al. 2019), e.g. in maritime pine (Chambel et al. 2007, Corcuera et al. 2010, de la Mata et al. 2012, Vizcaíno-Palomar et al. 2020). This plastic component is also generally associated with the climatic conditions experienced by the trees (Franks et al. 2014, Benito Garzón et al. 2019), allowing them to overcome changing climate up to a certain threshold (Matesanz et al. 2010, Nicotra et al. 2010, Valladares et al. 2014a). In our study, the plastic component of height growth was largely higher than the genetic component (fig. 2) and, although climate plays a role, was likely to be driven by multiple and interacting drivers including the biotic environment, soil quality, and other factors not considered in our study.

Plants also present an important genetic variation in plasticity (i.e. the genotype-by-environment interaction, G × E; Des Marais et al. 2013, Sork 2018), often approximated by the family or provenance-by-site interaction in forest tree common gardens, as is the case in our study. G × E is particularly prevalent for growth traits in trees (Li et al. 2017), as already shown in maritime pine (Alía et al. 1997, Corcuera et al. 2010, Correia et al. 2010, de la Mata et al. 2012; but see Chambel et al. (2007) where no provenance-specific responses were observed under two different watering regimes). In our study, provenance-by-site interaction was only weakly associated with height growth and the proportion of variance it explained was not different from zero (*model M2*; Table S5). Previous work in the context of tree breeding argued that G × E may hinder model transferability across sites and populations (Resende Jr et al. 2012, Resende et al. 2012). In maritime pine, our results suggest that large-scale predictions of height-growth variation will be only marginally impacted by not accounting for provenance-by-environment interaction. However, further work is necessary to assess the importance of the genetic variation of plasticity at the genotype level.

### 4.2 Potential drivers underlying height-growth genetic component

Our study shows that the height-growth genetic component in maritime pine is mostly associated with adaptation to climate, whose effect is partially confounded with the effect of gene pool assignment, reflecting both adaptive (different selective histories) and neutral processes (population demographic history and genetic drift) (fig. 2; see also Jaramillo-Correa et al. 2015). For example, the higher growth of most provenances from the French Atlantic gene pool (known for their high growth under a wide range of conditions, including Mediterranean sites in our study; see also Alía et al. 1997, Corcuera et al. 2010, de la Mata et al. 2012) was both associated with the provenance climate-of-origin and the gene pool assignment. As another example, in the northern Africa gene pool, the Madisouka (MAD) provenance was taller than the Tamrabta (TAM) provenance, which could be both explained by its noticeable ancestry proportion (23.3%) from the south-eastern Spanish gene pool (Jaramillo-Correa et al. 2015) or its adaptation to lower elevation (300 m lower than TAM). As a last example, the Leiria (LEI) provenance grew well in Asturias and Bordeaux as was the case for French Atlantic provenances (that share similar climates) but unlike them, it did not maintain growth in drier and warmer sites, probably due to a different genetic background (this provenance has a strong central Spain gene pool component; Table S3). Nevertheless, in contrast to the three examples above, for some provenances, the effects of the gene pool assignment and adaptation to climate on height growth could be clearly separated. This was the case, for example, for the Corsican provenances: the higher growth of Pinia (PIA) than Pineta (PIE) can only be explained by adaptation to different environmental conditions (and in particular climate), as both belong to the same gene pool. Indeed PIA is at the sea level under a climate similar to that of provenances from Central and south-eastern Spain whereas PIE is located at an altitude of 750 m a.s.l. in the mountains under a climate similar to that of the Atlantic provenances (fig. S9). These different adaptations within a same gene pool calls for a more targeted investigation of the Corsican gene pool. More generally, a *Q*_*ST*_ − *F*_*ST*_ analysis supported adaptive differentiation of height growth in maritime pine (see details in section 7 of the Supplementary Information).

The entanglement of the effect of climate adaptation and gene pool assignment to explain the genetic component of height-growth variation may partly stem from the distinct selective histories experienced in different parts of maritime pine range, despite gene pools being identified using genetic markers considered neutral (Jaramillo-Correa et al. 2015). This is supported by the estimation of gene pool-specific heritabilities in our study (*model M5*): the Corsican gene pool, and to a lesser extent the south-eastern Spain gene pool, have higher heritabilities than the French and Iberian Atlantic gene pools (Fig. S13; and see section 6.1.3 for a potential explanation of this pattern).

Overall, maritime pine proved to be a particularly suitable model species to study the joint influence of genetic neutral (population demographic history, genetic drift) and adaptive (climate adaptation) processes on quantitative traits. Further work on provenances that have different demographic histories but are exposed to similar climates (e.g. the LEI provenance and provenances from the Atlantic gene pools) would be relevant for understanding how a given genetic background guides population adaptation. Conversely, targeting provenances that have a similar demographic history but are found in highly contrasted environments (e.g. the Corsican provenances) would be valuable to identify signatures of adaptation while avoiding common issues due to confounding population structure (Berg et al. 2019, Sella and N. H. Barton 2019, Sohail et al. 2019). Likewise, investigating trait genetic architecture will also help better understand how adaptive and neutral processes have shaped the genotype-phenotype map and how this will influence future responses to selection (e.g. Kardos and Luikart 2021; see de Miguel et al. 2020 for maritime pine). Finally, it would also be critical to consider drivers of adaptation other than climate, such as resistance to pathogens or other biotic-related traits.

### 4.3 Towards integrating genomics into population response functions

Anticipating how provenances will grow in new environments is key to guide forest conservation strategies and population translocations to compensate for rapid climate change (Aitken and Whitlock 2013). To date, population response functions based on the climate in the provenance location have been the most widely used method for anticipating trait values when transplanting provenances in new environments (Rehfeldt et al. 1999, 2003, O’Neill et al. 2008, Wang et al. 2010, Pedlar and McKenney 2017, Rehfeldt et al. 2018, Fréjaville et al. 2020). Genome-informed predictive modeling of key adaptive traits is highly promising as it may provide a mean to further integrate adaptive or neutral genetic variation in the predictions, and to consider intraspecific variability at a finer scale than current models, thus gaining in prediction accuracy (Holliday et al. 2017). In valley oak, Browne et al. (2019) used genomic estimated breeding values (GEBVs; sum of the marker predicted effects, also known as polygenic scores) to identify genotypes that will grow faster under future climates. In lodgepole pine, Mahony et al. (2020) showed that phenotype-associated positive-effect alleles (PEAs, as used in our study) can predict phenotypic traits (e.g. cold injury) as well as climatic or geographical variables. In our study, we investigated whether including genomic information related to past demographic and selective processes resulting in distinct gene pools and counts of trait-associated alleles could improve range-wide height-growth predictions in maritime pine. Models combining climatic conditions in the provenance location, gene pool assignment, and PEAs captured most of the genetic component of height-growth variation (see fig. S10) and better predicted height growth of new provenances, compared to models relying exclusively on the common garden design or models including separately climatic or genomic information (see fig. 4). This suggests that range-wide trait prediction would benefit from jointly considering different sources of information (i.e. climatic and genomic), even though they may have overlapping effects (e.g. confounded effects of provenance climate-of-origin and gene pool assignment), as it may help to embrace the complexity and multidimensionality of the genetic component underlying quantitative traits. Noticeably, regional PEAs were generally better predictors of height growth in new provenances than gene pool assignment or provenance climate-of-origin as, when they were included alone in the models, they made better predictions in the driest common gardens (Madrid, Cáceres and Portugal) and similar ones to models combining multiple drivers of height growth variation in all common gardens except Bordeaux (P2 partition in fig. 4). Although this highlights the major role that trait-associated alleles identified using GWAS may play in predictive modeling, predicting traits of new provenances depends also on the number of provenances used to fit the models and the strength of the genetic relationship among them (Resende et al. 2012, Jarquín et al. 2014, Moghaddar et al. 2014, Hidalgo et al. 2016). This was reflected in our study by better predictive ability on new provenances in the P2 partition (random) compared to the P3 partition (containing provenances from underrepresented gene pools) for models including climatic and genomic information separately but not for models considering both jointly (fig. 4). Thus combining multiple sources of information may also be particularly relevant for predicting traits in marginal or difficult-to-access populations, as they normally belong to underrepresented geographical areas/gene pools in ecological and genetic studies.

**Figure 4.**
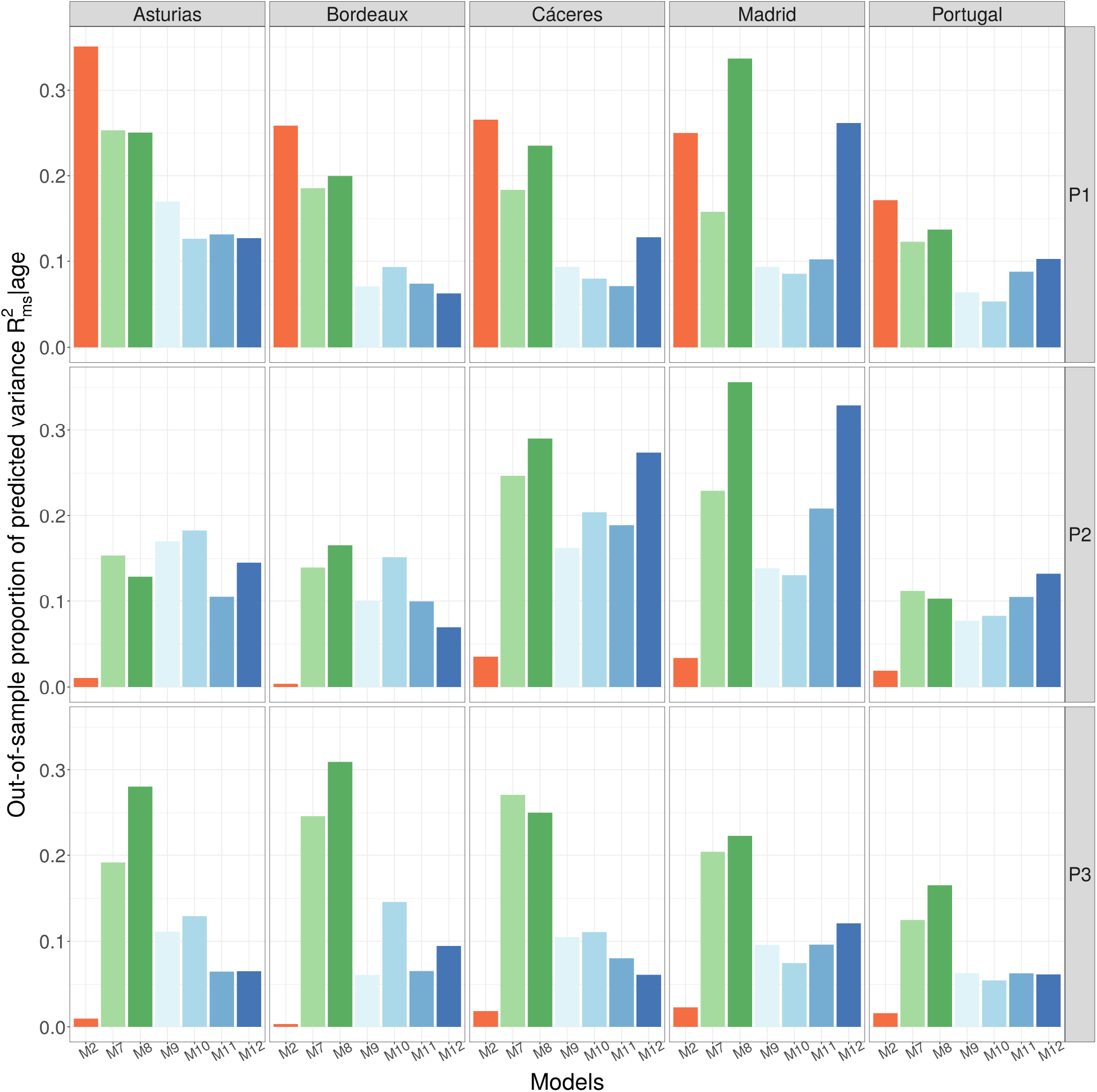
Model predictive ability on new observations (P1 partition) or new provenances (P2 and P3 partitions) based on the out-of-sample proportion of predicted variance conditional on the age effect (*prediction* 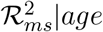) in the test datasets (data not used to fit the models). In the P1 partition, the training dataset was obtained by randomly sampling 75% of the observations and the test dataset contains the remaining 25% observations. In the P2 partition, the training dataset was obtained by randomly sampling 28 provenances and the test dataset contains the remaining 6 provenances. The P3 partition corresponds to a non-random split between a training dataset of 28 provenances and a test dataset containing 6 provenances with at least one provenance from each under-represented gene pool. The exact values of the *prediction* 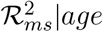 estimates and their associated credible intervals can be found in Tables S4 (P1 partition), S9 (P2 partition) and S12 (P3 partition).

The high predictive ability of PEAs, both alone and combined with climatic and gene pool information, was somehow unexpected given the sparse genomic sampling in our study: 5,165 SNPs to cover the 28 Gbp maritime pine genome (Zonneveld 2012). Indeed, conifers have particularly huge genomes, generally ranging from 18 to 35 Gbp (Mackay et al. 2012) and thus rendering the current cost of whole-genome resequencing prohibitive (Holliday et al. 2017). Targeted genotyping approaches, such as the one used in the present study, select candidate genes based on previous population and functional studies, thus allowing to include potential targets of selection and climate adaptation, but probably inducing an ascertainment bias (Jaramillo-Correa et al. 2015). However, as height is a particularly polygenic trait (degree of polygenicity estimated at ∼7% in de Miguel et al. 2020), we were able to identify a considerable number of PEAs despite the weak genome coverage of our study. Further genomic sampling would be highly valuable to capture the polygenic architecture of height more broadly, turning PEAs into much better predictors than the provenance climate-of-origin or the gene pool assignment, and ultimately making climatic data redundant, at least for main range populations (see above for marginal populations). This would also allow to characterize the genetic variation within provenances more precisely, thereby increasing the estimation accuracy and reducing the residual variance. Similar to Mahony et al. (2020) and MacLachlan et al. (2021) who selected the positive-effect alleles as the 1% of SNPs that showed the strongest association with phenotypes (estimated via a GWAS performed on 18,525 SNPs), we used PEA counts instead of the more commonly used polygenic scores (Pritchard et al. 2010, Browne et al. 2019, Fuller et al. 2020). Unlike polygenic scores, PEAs do not account for allele effect sizes, thus minimizing the circularity of the analysis (i.e. effect sizes that are estimated based on the same dataset as the one used for the models, only serve for PEAs identification) and potentially enhancing the prediction accuracy across genetic groups compared to polygenic scores. Indeed, low observed transferability of polygenic scores across genetic groups (Martin et al. 2017, N. Barton et al. 2019, Martin et al. 2019) may stem from varying effect sizes of “peripheral” alleles (i.e. alleles indirectly affecting the phenotype), as suggested in Mathieson 2021).

Although combining climatic and genomic information allowed us to capture most of the genetic component of height-growth variation (fig. S10), the residual variance remained high in our study. As already mentioned, this may be partly related to the models’ difficulty in accounting for genetic variation within provenances, which might be improved by denser genomic sampling. However, this unexplained variance may also originate from developmental stochasticity, which can play an important role in explaining differences between individuals with the same genotype (Vogt 2015, Ballouz et al. 2019). Height growth may also be influenced by the correlative effects of other traits. For example, Stern et al. (2020) recently showed that variation in some human traits (hair color and educational attainment), previously thought to be under selection, can instead be explained by indirect selection via a correlated response to other traits. Therefore, multi-trait models may be the next necessary step to improve our understanding and predictive ability of quantitative trait variation at large geographical scales (e.g. Csilléry et al. 2020).

A last noticeable results was that rPEAs (positive-effect alleles identified in specific geographical regions, i.e. particular environments) had generally a higher predictive ability than gPEAs (positive-effect alleles identified range-wide) (Figure 4). Interestingly, only a small proportion of rPEAs were shared among geographical regions in our study (20% shared between the Iberian and French Atlantic regions, 12% between the French Atlantic and Mediterranean regions, and 24% between the Iberian Atlantic and Mediterranean regions; Figure S2), although we cannot exclude that the proportion of shared rPEAs among regions is a function of the sample size (see details in the section 2.2 of the Supplementary Information). Moreover, those that were shared among different regions showed consistently similar effects across regions (e.g. positive effects in two or more regions rather than antagonist effects). This supports the predominance of conditional neutrality, i.e. alleles that are advantageous in some environments and neutral in others, over antagonistic pleiotropy, i.e. alleles that are advantageous in some environments and disadvantageous in others (Tiffin and Ross-Ibarra 2014). Such pattern has already been reported in plants (Prunier et al. 2012, Anderson et al. 2013). Our results show that, despite a high stability in the level of polygenicity for height between the Atlantic and Mediterranean regions (de Miguel et al. 2020), height-growth variation in Mediterranean sites is unlikely to be affected by the same loci as in the other regions, probably as a result of genetic divergence in separated southern refugia during the last glaciation. Overall, identifying positive-effect alleles for different geographical regions separately has the potential to greatly improve the predictive ability of the models, but at the cost of reducing GWAS power (due to lower sample size than in global, wide-range analyses).

Finally, caution has to be taken when generalizing our results to older trees as the drivers of height growth in young trees may differ from that of adult trees. For example, G × E on tree height can be age-dependant (Gwaze et al. 2001, Zas et al. 2003, Rehfeldt et al. 2018) and the plastic component may be higher in younger trees, especially in maritime pine (Vizcaíno-Palomar et al. 2020). Nevertheless, a recent measurement in the Bordeaux common garden (2018) showed a high correlation between young saplings and 10-year old trees for height (Pearson’s correlation coefficient of 0.893 based on height BLUPs; see de Miguel et al. 2020 for details on BLUP estimation). Moreover, our study remains indicative of how trees respond to varying environmental conditions during establishment and early-growing stages, a critical phase where most mortality (i.e. selection) is expected to take place (Postma and Ågren 2016). In addition to ontogenic effects, high mortality in the Mediterranean common gardens (Cáceres and Madrid), after a marked summer drought, may have biased estimates of some parameters of interest. Indeed, if this environmental filtering was not independent of tree height, it could have resulted in an underestimation of the genetic variance. Nonetheless, height distributions in Cáceres and Madrid were only slightly right-skewed, suggesting uniform selection across height classes (fig. S21), and thus no bias due to high mortality in these common gardens.

## 5 Conclusion

The present study connects climate-based population response functions that have been extensively used in predictive models for forest trees (Rehfeldt et al. 1999, 2003, Wang et al. 2010, Leites et al. 2012a) with recent genomic approaches to investigate the potential drivers behind the genetic and plastic components of height-growth variation and predict how provenances will grow when transplanted into new climates. The integration of genomic data into range-wide predictive models is in its infancy and still lacks a well-established framework, especially for non-model species such as forest trees. We showed that combining climatic and genomic information (i.e. provenance climate-of-origin, gene pool assignment and trait-associated positive-effect allele counts) can improve model predictions for a highly polygenic adaptive trait such as height growth, despite sparse genomic sampling. Further genomic sampling may help to improve the accuracy of the estimates, notably through improved characterization of within-provenance genetic variation. Moreover, comparative studies between maritime pine and more continuously distributed species (e.g. Scots pine; Alberto et al. 2013) and/or living under stronger climatic limitations, would be highly valuable to determine whether our findings can be generalized to species with contrasted population demographic and selective history. Finally, our study focuses specifically on the height-growth genetic component of standing populations, but considering evolutionary processes (e.g. genetic drift in small populations, extreme selection events, etc.) into the predictions would be necessary to anticipate the response of future forest tree generations to changing climatic conditions and thus provide a much-needed longer-term vision (Waldvogel et al. 2020)

## 6 Acknowledgements

We thank A. Saldaña, F. del Caño, E. Ballesteros and D. Barba (INIA) and the ‘Unité Expérimentale Forêt Pierroton’ (UEFP, INRAE; doi:10.15454/1.5483264699193726E12) for field assistance (plantation and measurements). Data used in this research are part of the Spanish Network of Genetic Trials (GENFORED, http://www.genfored.es). We thank all persons and institutions linked to the establishment and maintenance of field trials used in this study. We are also very grateful to Ricardo Alía who contributed to the design and establishment of the CLONAPIN network and provided comments on the manuscript, and to Benjamin Brachi, Thibault Poiret, Andrew J. Eckert and one anonymous reviewer who provided constructive and valuable comments on a previous version of the manuscript. Thanks are extended to Juan Majada for initiating and supervising the establishment of the CLONAPIN network. JA was funded by the University of Bordeaux (ministerial grant). This study was funded by the Spanish Ministry of Economy and Competitiveness through projects RTA2010-00120-C02-02 (CLONAPIN), CGL2011-30182-C02-01 (AdapCon) and AGL2012-40151-C03-02 (FENOPIN). The study was also supported by the ‘Initiative d’Excellence (IdEx) de l’Université de Bordeaux - Chaires d’installation 2015’ (EcoGenPin) and the European Union’s Horizon 2020 research and innovation programme under grant agreement No 773383 (B4EST).

## 7 Author contributions

SCG-M and CP designed the experiment and supervised the curation of field data. MdM cleaned and formatted the phenotypic and genomic data, and produced the BLUPs used in GWAS. SCG-M and MdM ran the GWAS to identify the positive-effect alleles. SCG-M, MBG, JA and FB conceived the paper methodology. JA conducted the data analyses. SCG-M, MBG, JA and FB interpreted the results. JA led the writing of the manuscript. All authors contributed to the manuscript and gave final approval for publication.

## 8 Data and script availability

Data are publicly available. SNP data were deposited in the Dryad repository at http://dx.doi.org/10.5061/dryad.8d6k1. Height data have been deposited in GENFORED, the Spanish Network of Genetic Trials (http://www.genfored.es). Scripts are available at https://github.com/JulietteArchambeau/HeightPinpinClonapin.

