## Supplementary Information for "Combining climatic and genomic data improves range-wide tree height growth prediction in a forest tree"

**1 Details about the experimental design** **3**

**2 Height-associated positive-effect alleles (PEAs)** **6**

**7 $Q_{ST} - F_{ST}$ analysis 54**

**8 Distribution of heights in Cáceres and Madrid. 55**

### 1 Details about the experimental design

| | All | Training P1 | Test P1 | Training P2 | Test P2 | Training P3 | Test P3 |
|---|---|---|---|---|---|---|---|
| **All sites** | 33,121 | 24,840 | 8,281 | 27,349 | 5,772 | 26,172 | 6,949 |
| **Asturias** | 11,920 | 8,934 | 2,986 | 9,813 | 2,107 | 9,420 | 2,500 |
| **Bordeaux** | 6,473 | 4,836 | 1,637 | 5,285 | 1,188 | 5,063 | 1,410 |
| **Cáceres** | 340 | 249 | 91 | 297 | 43 | 272 | 68 |
| **Madrid** | 1,046 | 807 | 239 | 876 | 170 | 855 | 191 |
| **Portugal** | 13,342 | 10,014 | 3,328 | 11,078 | 2,264 | 10,562 | 2,780 |
| **Asturias - 10 months old** | 3,967 | 2,949 | 1,018 | 3,268 | 699 | 3133 | 834 |
| **Asturias - 21 months old** | 3,979 | 3,022 | 957 | 3,275 | 704 | 3,143 | 836 |
| **Asturias - 37 months old** | 3,974 | 2,963 | 1,011 | 3,270 | 704 | 3,144 | 830 |
| **Bordeaux - 25 months old** | 3,237 | 2,420 | 817 | 2,643 | 594 | 2,532 | 705 |
| **Bordeaux - 37 months old** | 3,236 | 2,416 | 820 | 2,642 | 594 | 2,531 | 705 |
| **Cáceres - 8 months old** | 340 | 249 | 91 | 297 | 43 | 272 | 68 |
| **Madrid - 13 months old** | 1,046 | 807 | 239 | 876 | 170 | 855 | 191 |
| **Portugal - 11 months old** | 4,152 | 3,102 | 1,050 | 3,442 | 710 | 3,266 | 886 |
| **Portugal - 15 months old** | 3,773 | 2,833 | 940 | 3,134 | 639 | 2,980 | 793 |
| **Portugal - 20 months old** | 2,752 | 2,078 | 674 | 2,288 | 464 | 2,192 | 560 |
| **Portugal - 27 months old** | 2,665 | 2,001 | 664 | 2,214 | 451 | 2,124 | 541 |

**Table S1:** Number of observations in the entire dataset (after filtering) and in each of the three partitions. In the P1 partition, the training dataset was obtained by randomly sampling 75% of the observations and the test dataset contains the remaining 25% observations. In the P2 partition, the training dataset was obtained by randomly sampling 28 provenances and the test dataset contains the remaining 6 provenances. The P3 partition corresponds to a non-random split between a training dataset of 28 provenances and a test dataset containing 6 provenances with at least one provenance from each under-represented gene pool (i.e. northern Africa, south-eastern Spain and Corsican gene pools).

| Code | Name | Number of genotypes | Number of trees | Number of observations |
|---|---|---|---|---|
| **ALT** | Alto de la Llama | 9 | 216 | 580 |
| **ARM** | Armayán | 8 | 213 | 547 |
| **ARN** | Arenas de San Pedro | 17 | 412 | 1083 |
| **BAY** | Bayubas de Abajo | 18 | 462 | 1181 |
| **BON** | Boniches | 9 | 221 | 594 |
| **CAD** | Cadavedo | 10 | 245 | 658 |
| **CAR** | Carbonero el Mayor | 6 | 156 | 398 |
| **CAS** | Castropol | 10 | 246 | 642 |
| **CEN** | Cenicientos | 9 | 207 | 561 |
| **COC** | Coca | 18 | 424 | 1114 |
| **COM** | Cómpeta | 4 | 109 | 272 |
| **CUE** | Cuellar | 28 | 680 | 1750 |
| **HOU** | Hourtin | 26 | 645 | 1669 |
| **LAM** | Lamuño | 9 | 216 | 563 |
| **LEI** | Leiria | 23 | 549 | 1439 |
| **MAD** | Madisouka | 1 | 19 | 54 |
| **MIM** | Mimizan | 18 | 445 | 1111 |
| **OLB** | Olonne sur Mer | 22 | 552 | 1476 |
| **OLO** | Olba | 24 | 563 | 1441 |
| **ORI** | Oria | 26 | 651 | 1720 |
| **PET** | Petrocq | 24 | 594 | 1496 |
| **PIA** | Pinia | 16 | 413 | 1046 |
| **PIE** | Pineta | 9 | 220 | 582 |
| **PLE** | Pleucadec | 20 | 480 | 1234 |
| **PUE** | Puerto de Vega | 8 | 198 | 497 |
| **QUA** | Quatretonda | 17 | 448 | 1156 |
| **SAC** | San Cipriano de Ribaterme | 9 | 208 | 499 |
| **SAL** | San Leonardo | 14 | 323 | 804 |
| **SEG** | Sergude (Huerto Semillero) | 21 | 536 | 1340 |
| **SIE** | Sierra de Barcia | 8 | 203 | 506 |
| **STJ** | St-Jean des Monts | 28 | 718 | 1824 |
| **TAM** | Tamrabta | 15 | 320 | 839 |
| **VAL** | Valdemaqueda | 12 | 286 | 750 |
| **VER** | Le Verdon | 27 | 663 | 1695 |

**Table S2:** Provenance information: provenance codes used in the study, provenance names, number of genotypes, trees and observations (an observation being a height-growth measurement in a given year on one individual) per provenance.

| Provenance | NA | C | CS | FA | IA | SES |
|---|---|---|---|---|---|---|
| **ALT** | 0.003 | 0.000 | 0.119 | 0.096 | **0.780** | 0.003 |
| **ARM** | 0.005 | 0.007 | 0.021 | 0.006 | **0.959** | 0.001 |
| **ARN** | 0.010 | 0.002 | **0.958** | 0.007 | 0.010 | 0.013 |
| **BAY** | 0.003 | 0.004 | **0.966** | 0.013 | 0.010 | 0.004 |
| **BON** | 0.152 | 0.010 | **0.654** | 0.003 | 0.002 | 0.179 |
| **CAD** | 0.002 | 0.001 | 0.053 | 0.010 | **0.933** | 0.002 |
| **CAR** | 0.001 | 0.001 | **0.904** | 0.060 | 0.022 | 0.011 |
| **CAS** | 0.001 | 0.000 | 0.005 | 0.001 | **0.991** | 0.001 |
| **CEN** | 0.013 | 0.002 | **0.892** | 0.003 | 0.041 | 0.050 |
| **COC** | 0.017 | 0.005 | **0.826** | 0.061 | 0.043 | 0.047 |
| **COM** | 0.239 | 0.028 | 0.127 | 0.011 | 0.039 | **0.556** |
| **CUE** | 0.003 | 0.001 | **0.874** | 0.063 | 0.056 | 0.002 |
| **HOU** | 0.004 | 0.001 | 0.026 | **0.960** | 0.007 | 0.002 |
| **LAM** | 0.002 | 0.001 | 0.003 | 0.050 | **0.943** | 0.000 |
| **LEI** | 0.004 | 0.003 | **0.512** | 0.007 | 0.472 | 0.002 |
| **MAD** | **0.764** | 0.001 | 0.000 | 0.002 | 0.000 | 0.233 |
| **MIM** | 0.004 | 0.002 | 0.024 | **0.952** | 0.013 | 0.005 |
| **OLB** | 0.080 | 0.010 | **0.776** | 0.006 | 0.003 | 0.126 |
| **OLO** | 0.004 | 0.001 | 0.007 | **0.980** | 0.006 | 0.003 |
| **ORI** | 0.249 | 0.005 | 0.027 | 0.001 | 0.009 | **0.709** |
| **PET** | 0.003 | 0.001 | 0.021 | **0.966** | 0.004 | 0.004 |
| **PIA** | 0.004 | **0.974** | 0.010 | 0.001 | 0.008 | 0.003 |
| **PIE** | 0.007 | **0.970** | 0.022 | 0.000 | 0.000 | 0.001 |
| **PLE** | 0.005 | 0.001 | 0.060 | **0.924** | 0.005 | 0.004 |
| **PUE** | 0.002 | 0.000 | 0.021 | 0.001 | **0.974** | 0.002 |
| **QUA** | 0.092 | 0.006 | **0.499** | 0.005 | 0.015 | 0.383 |
| **SAC** | 0.004 | 0.001 | 0.268 | 0.002 | **0.723** | 0.002 |
| **SAL** | 0.010 | 0.004 | **0.944** | 0.027 | 0.008 | 0.008 |
| **SEG** | 0.003 | 0.001 | 0.151 | 0.013 | **0.829** | 0.003 |
| **SIE** | 0.003 | 0.001 | 0.089 | 0.015 | **0.891** | 0.001 |
| **STJ** | 0.003 | 0.002 | 0.027 | **0.947** | 0.017 | 0.004 |
| **TAM** | **0.937** | 0.000 | 0.025 | 0.000 | 0.037 | 0.001 |
| **VAL** | 0.012 | 0.003 | **0.943** | 0.006 | 0.011 | 0.025 |
| **VER** | 0.003 | 0.002 | 0.019 | **0.972** | 0.003 | 0.001 |

**Table S3:** Mean proportion belonging to each gene pool for each provenance. For each provenance, the highest proportion belonging to a given gene pool is in bold. The gene pools come from: northern Africa (NA), Corsica (C), central Spain (CS), French Atlantic region (FA), Iberian Atlantic region (IA) and south-eastern Spain (SES).

#### 2 Height-associated positive-effect alleles (PEAs)

##### 2.1 Calculation of the counts of height-associated positive-effect alleles

This section complements the section 2.2 in the manuscript and we explain here in more details how we calculated the counts of global and regional height-associated positive-effect alleles. As already explained in the manuscript, for each of the four GWAS (a global GWAS and three regional GWAS), we selected the 350 SNPs with the highest absolute estimates of the posterior effect size (i.e. Rao-Blackwellized estimates), corresponding approximately to the estimated number of SNPs with non-zero effects on height in a previous study (i.e. the level of polygenicity; de Miguel et al. 2020). Then, for each selected allele, if the posterior effect size was negative, it was converted to positive values and the reference allele flipped to select only alleles that have a positive effect on height (positive-effect alleles; PEAs). Thus, we ended up with four groups of PEAs. One group had a global positive effect (i.e. range-wide effect) on height and was used to calculate the count of global PEAs that each sapling has ($gPEA_g$ variable). Trees with the same genotype had the same $gPEA_g$, such as $gPEA_g = \sum_{l=1}^{350} G_{lg}$, where $G_{lg} = \{0, 1, 2\}$ is the number of global PEAs that the genotype $g$ has at the locus $l$. The three other groups of PEAs had a regional positive-effect on height (i.e. specific effect in a given geographical region/in a particular environment) and were used to calculate the number of regional PEAs that each tree has ($rPEA_{gr}$ variable). $rPEA_{gr}$ was calculated based on both the tree genotype $g$ and the region $r$ of its planting site, such as $rPEA_{gr} = \sum_{l=1}^{350} G_{lgr}$, where $G_{lgr} = \{0, 1, 2\}$ is the number of PEAs specific to the geographical region $r$ that the genotype $g$ has at the locus $l$ (see fig. S1 below for a summary diagram of how the counts of global and regional PEAs were obtained).

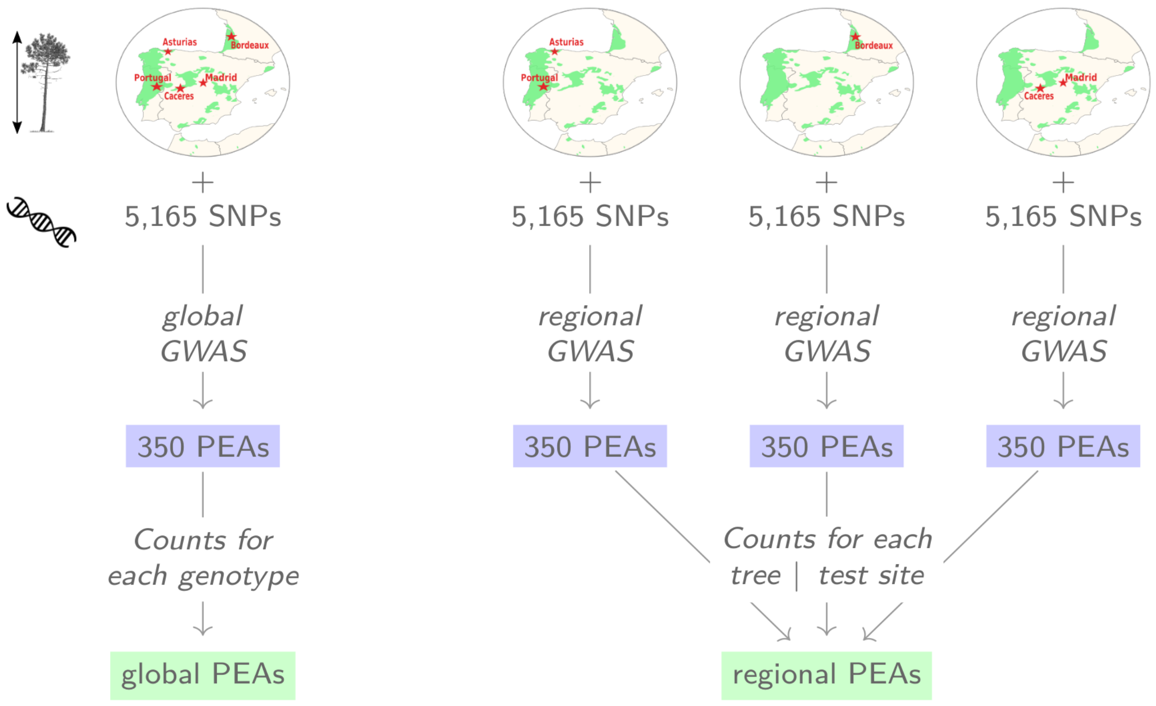

**Figure S1.** Schematic representation of the calculation of the global PEA counts ($gPEAg$ variable) and regional PEA counts ($rPEAgr$ variable).

#### 2.2 Shared proportion of globally and regionally selected height-associated SNPs

A small proportion of regionally-selected height-associated SNPs was shared among the different regions: 20% (69 SNPs) shared between the French Atlantic (Bordeaux) and the Iberian Atlantic region (Asturias and Portugal), 12% (41 SNPs) shared between the Mediterranean (Cáceres and Madrid) and the French Atlantic region and 24% (83 SNPs) shared between the Iberian Atlantic and the Mediterranean region (fig. S2). Interestingly, height-associated SNPs that were shared among different regions show consistently similar effects across regions (e.g. positive effects in two or more regions rather than antagonist effects): we did not detect a single SNP with an antagonistic effect (fig. S2).

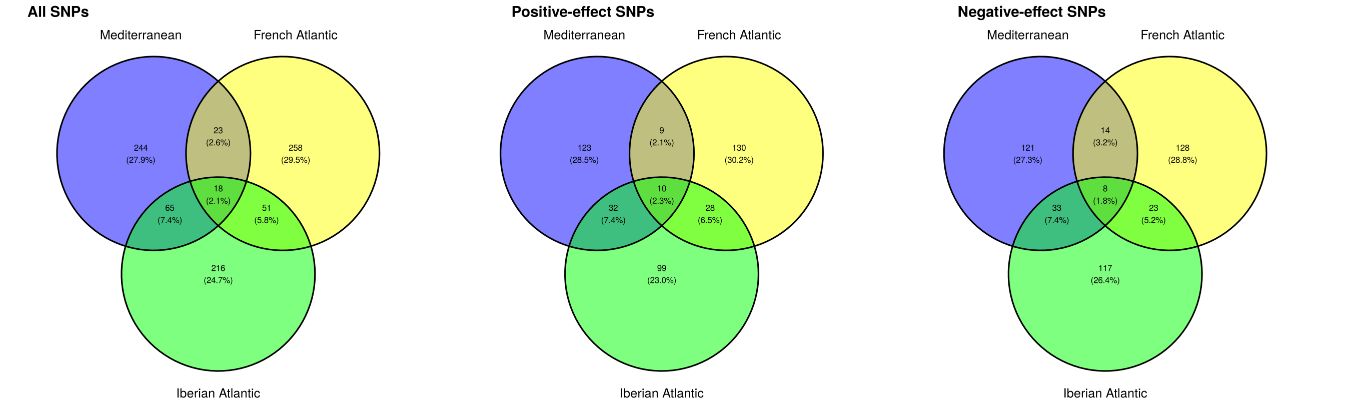

**Figure S2.** Number and proportion of height-associated alleles shared among regions: the French Atlantic region (Bordeaux), the Iberian Atlantic region (Asturias and Portugal) and the Mediterranean region (Cáceres and Madrid). This figure was obtained before transforming the height-associated negative-effect alleles in positive-effect alleles by flipping their reference allele.

82.9% of the globally-selected SNPs were at least selected once in a regional GWAS too (fig. S3). Height-associated SNPs that were selected both globally and regionally (at least in one region) show consistently similar effects (i.e. either positive or negative, but not antagonistic effects): we did not find a single SNP with an antagonist effect when selected globally or regionally.

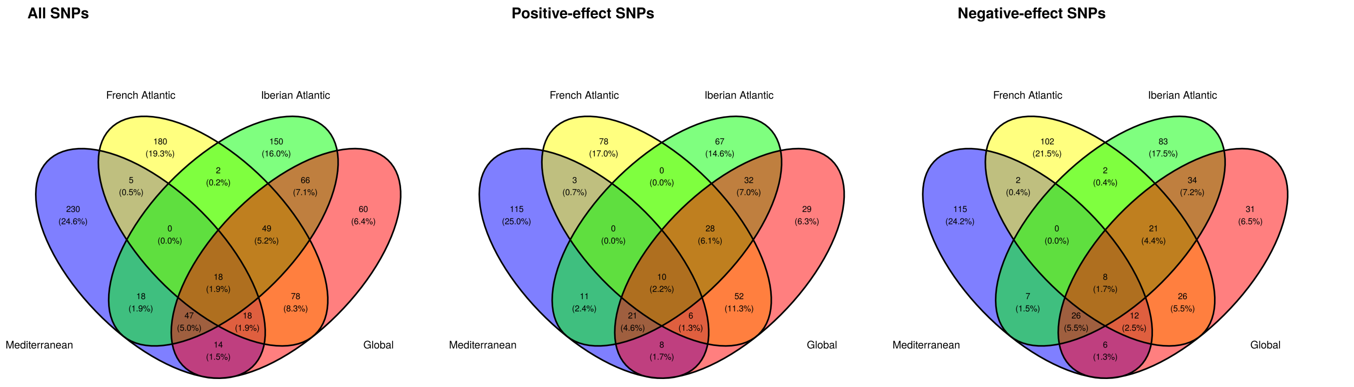

**Figure S3.** Number and proportion of alleles shared among the sets of 350 height-associated alleles selected globally and those selected regionally.

Importantly, we cannot exclude that the proportion of SNPs shared among regions or shared among globally-selected and regionally-selected SNPs is a function of the sample size. Indeed, the two regions with the lowest sample size (French Atlantic and Mediterranean regions) are also the ones that share the lowest number of height-associated SNPs (fig. S2). Similarly, the globally-selected height-associated SNPs share the highest proportion with regionally-selected height-associated SNPs from the Iberian Atlantic region (the region with the highest sample size) and the lowest proportion with regionally-selected height-associated SNPs from the Mediterranean region (the region with the lowest sample size) (fig. S3).

### 3 Climatic data

#### 3.1 In the test sites

We extracted monthly climatic data from the EuMedClim database at 1-km resolution (Fréjaville and Benito Garzón 2018). We calculated six variables that describe both extreme and average temperature and precipitation conditions in the test sites during the year preceding the measurements: the mean of monthly precipitation (*mean.pre*, mm), minimum of monthly minimum temperatures (*min.tmn*, °C), minimum of monthly precipitation during summer -June to September- (*min.presummer*, °C), the mean of monthly maximum temperatures (*mean.tmax*, °C), maximum of monthly precipitation (*max.pre*, mm), maximum of monthly maximum temperatures (*max.tmx*, °C). These variables had at most a correlation coefficient of 0.85 among each other (fig. S4). Due to the unbalanced number of measurements among test sites (trees were measured only once in the hottest and driest sites, Cáceres and Madrid, as survival was very low), some of these variables were slightly correlated with tree age (at most with a correlation coefficient of 0.56 for the mean of the monthly precipitation; fig. S4). We decided not to include soil variables (from the European Soil Database: https://esdac.jrc.ec.europa.eu/) in the analyses as they were highly correlated to some of the climatic variables. Likewise, we did not include variables related to water balance or evapotranspiration potential as they were highly correlated with temperature and precipitation variables.

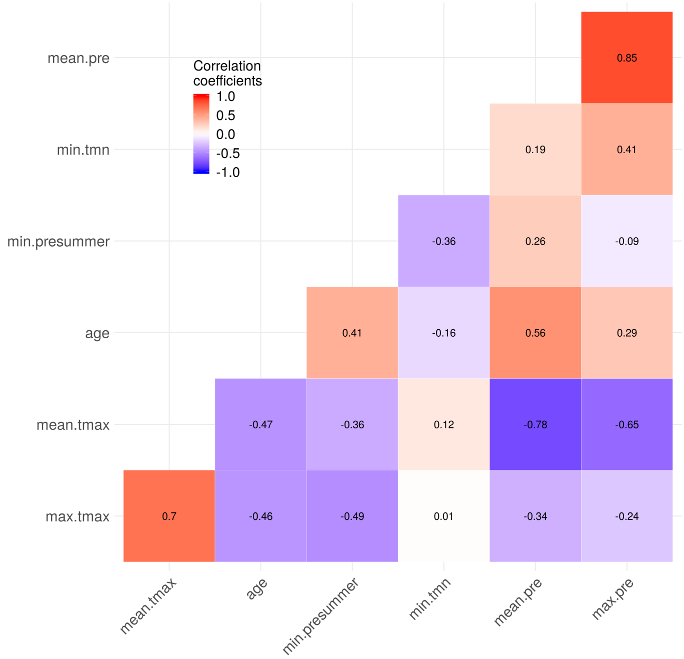

**Figure S4.** Correlation matrix between variables related to the climatic conditions in the test sites and tree age at the time of the measurements. The climatic variables, calculated over the year preceding the measurements, are: the mean of monthly precipitation (*mean.pre*, mm), minimum of monthly minimum temperatures (*min.tmn*, °C), minimum of monthly precipitation during summer -June to September- (*min.presummer*, °C), the mean of monthly maximum temperatures (*mean.tmax*, °C), maximum of monthly precipitation (*max.pre*, mm), maximum of monthly maximum temperatures (*max.tmx*, °C).

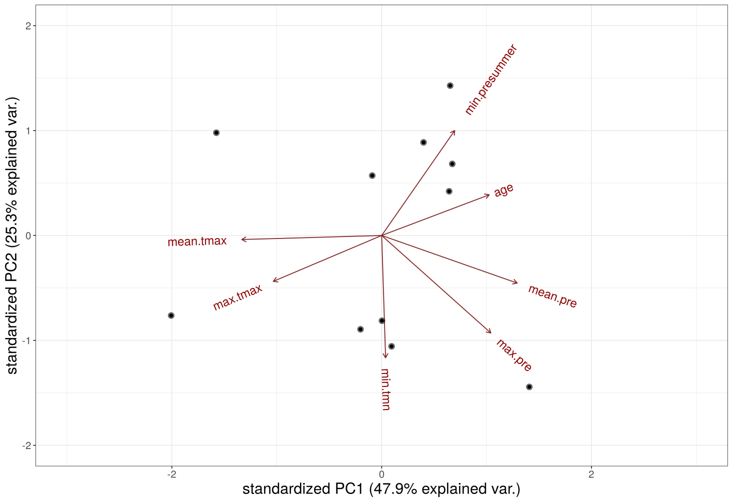

**Figure S5.** Principal component analysis of the variables related to the climatic conditions in the test sites and tree age at the time of the measurements. See fig. S4 for the meaning of variable abbreviations.

The climatic similarity among test sites during the year preceding the measurements was described by the covariance matrix $\Omega$. This covariance matrix was used to estimate the association between height-growth variation and the climatic similarity between test sites in *models M3* to *M6* (Table 1), following Jarquín et al. (2014); see also a similar approach but using Euclidean distance matrices in Thomson et al. (2018).

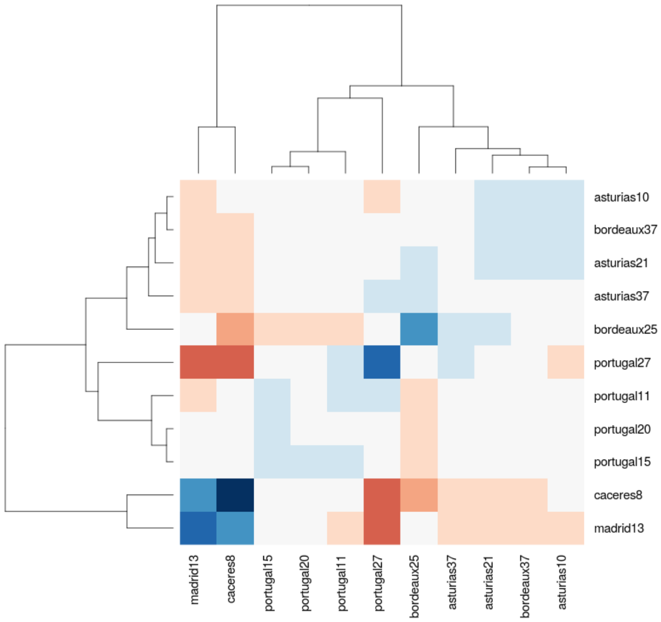

**Figure S6.** Heatmap of the covariance matrix $\Omega$ describing the climatic similarity among test sites during the year preceding the measurements. The labels correspond to the name of the test sites followed by the age of the trees at the date of the measurement (in months).

#### 3.2 In the provenances

We extracted yearly data from the EuMedClim database at 1-km resolution (Fréjaville and Benito Garzón 2018). We calculated four variables that describe the mean temperature and precipitation in the provenance locations over the period from 1901 to 2009, representing the climate under which provenances have evolved: the average of the annual daily mean temperature (*mean.temp*, °C), the average of the maximum temperature of the warmest month (*max.temp*, °C), the average of the annual precipitation (*mean.pre*, mm) and the average of the precipitation of the driest month (*min.pre*, mm). These variables had at most a correlation coefficient of 0.77 among each other (fig. S7), and three of them were correlated to the population genetic structure, i.e. the gene pool assignment ($|\rho| \geq 0.6$). Indeed, the provenance proportion belonging to the French Atlantic gene pool was positively correlated ($\rho$=0.83) to the average of the precipitation during the driest month, whereas the proportion belonging to the Central Spain gene pool was negatively correlated ($\rho$=-0.68) to the average of the annual precipitation and positively correlated ($\rho$=0.6) to the average of the maximum temperature of the warmest month (fig. S7). However, the confounding effect introduced by these correlations was mitigated by some provenances belonging to different gene pools but occurring in similar climates (i.e. French and Iberian Atlantic provenances), and some provenances occurring in different climates but belonging to the same gene pool (i.e. Corsican provenances). Soil variables from the European Soil Database (https://esdac.jrc.ec.europa.eu/) were not included in our study as they were highly correlated to some of the selected climatic variables.

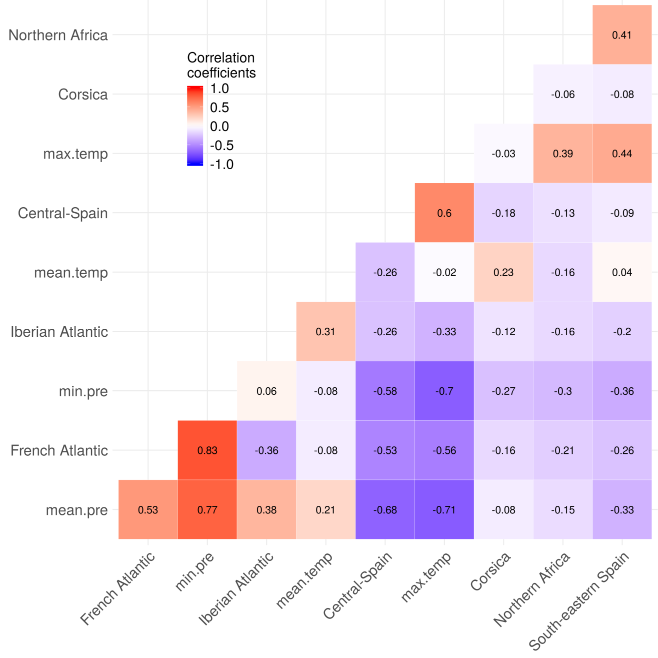

**Figure S7.** Correlation matrix of the variables related to the climatic conditions in the provenance locations and variables related to the population genetic structure (genotype proportion belonging to each gene pool). The genotypes belong to 6 distinct gene pools from: Northern Africa, Corsica, Central Spain, French Atlantic region, Iberian Atlantic region and south-eastern Spain. The climatic variables, calculated over the period from 1901 to 2009, are: the average of the annual daily mean temperature (*mean.temp*, °C), the average of the maximum temperature of the warmest month (*max.temp*, °C), the average of the annual precipitation (*mean.pre*, mm) and the average of the precipitation of the driest month (*min.pre*, mm).

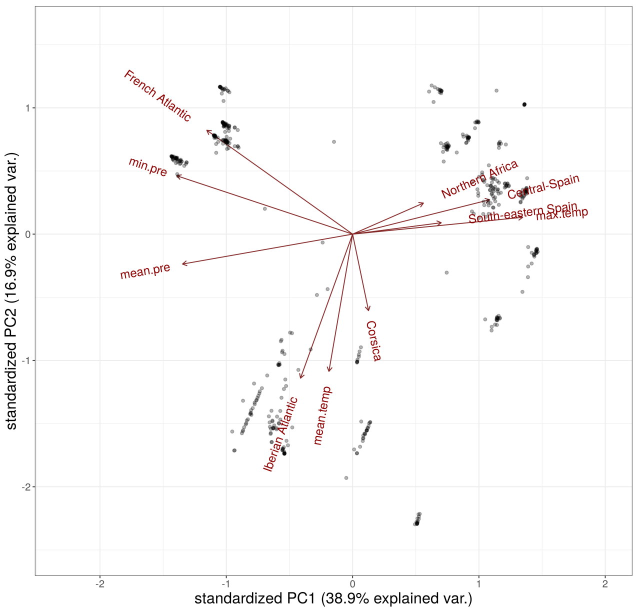

**Figure S8.** Principal component analysis of the variables related to the climatic conditions in the provenance locations and variables related to the population genetic structure (genotype proportion belonging to each gene pool). See fig. S7 for the meaning of variable abbreviations.

The climatic similarity among provenances was described by the covariance matrix $\Phi$. This covariance matrix was used to estimate the association between height-growth variation and the climatic similarity between provenances in *model M6*, following Jarquín et al. (2014); see also a similar approach but using Euclidean distance matrices in Thomson et al. (2018).

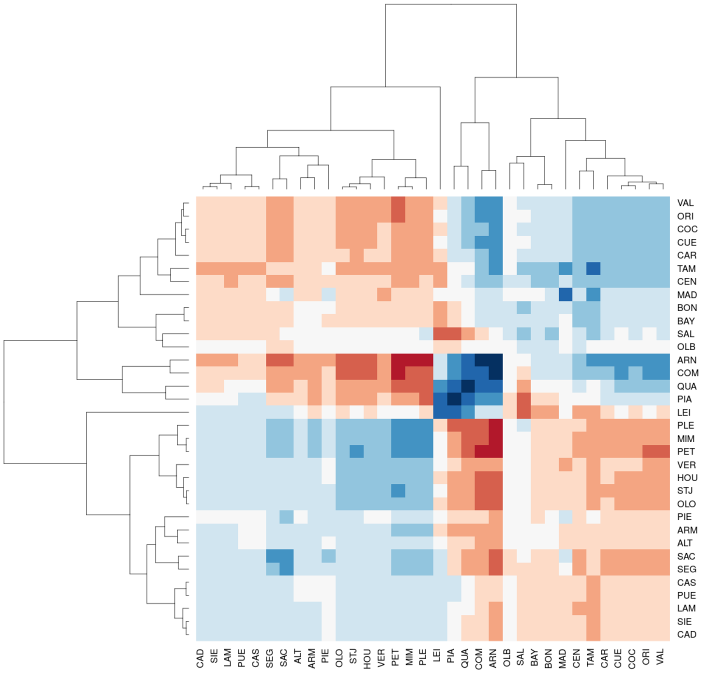

**Figure S9.** Heatmap of the covariance matrix Φ of the climatic variables in the provenances. Labels correspond to the provenance names.

### 4 Model equations and priors

#### 4.1 Model equations

In this section, the complete equations of each model are specified. The priors are indicated in the next section.

##### 4.1.1 Baseline *models M1* and *M2*: separating the genetic and plastic components

***Model M0*:**

$$\begin{aligned}\log(h_{isb}) &\sim \mathcal{N}(\mathbf{X}\beta + \mu_{sb}\,,\,\sigma^2)\\ \mathbf{X}\beta &= \beta_0 + \beta_{age}\text{age}_i + \beta_{age2}\text{age}_i^2\\ \mu_{sb} &= S_s + B_{b(s)}\end{aligned}$$

***Model M1*:**

$$\begin{aligned}\log(h_{isbpg}) &\sim \mathcal{N}(\mathbf{X}\beta + \mu_{sbpg}\,,\,\sigma^2)\\ \mathbf{X}\beta &= \beta_0 + \beta_{age}\text{age}_i + \beta_{age2}\text{age}_i^2\\ \mu_{sbpg} &= S_s + B_{b(s)} + P_p + G_{g(p)}\end{aligned}$$

***Model M2:***

$$\begin{aligned}\log(h_{isbpg}) &\sim \mathcal{N}(\mathbf{X}\beta + \mu_{sbpg}\,,\,\sigma^2)\\ \mathbf{X}\beta &= \beta_0 + \beta_{age}\text{age}_i + \beta_{age2}\text{age}_i^2\\ \mu_{sbpg} &= S_s + B_{b(s)} + P_p + G_{g(p)} + S_sP_p\end{aligned}$$

where $\mathbf{X}$ is the 3-column design matrix and $\beta$ is a vector including the intercept $\beta_0$ and the coefficients $\beta_{age}$ and $\beta_{age2}$ of the fixed effect variables ($age$ and $age^2$, respectively). $\mu_{sbpg}$ is the vector of varying intercepts with the provenance intercepts $P_p$, the genotype intercepts $G_{g(p)}$, the site intercepts $S_s$, the block intercepts $B_{b(s)}$ and the interaction between the site and provenance intercepts $S_sP_p$.

##### 4.1.2 Explanatory *models M3* to *M6*: potential drivers underlying height-growth variation

***Model M3*:**

$$\begin{aligned}\log(h_{isbpg}) &\sim \mathcal{N}(\mathbf{X}\beta + \mu_{sbpg}\,,\,\sigma^2)\\ \mathbf{X}\beta &= \beta_0 + \beta_{age}\text{age}_i + \beta_{age2}\text{age}_i^2\\ \mu_{isbpg} &= S_s + B_{b(s)} + P_p + G_{g(p)} + cs_{is}\\ cs_{is} &\sim \mathcal{N}(0, \Omega\sigma^2_{cs_{is}})\end{aligned}$$

***Models M4*, *M5* and *M6*:**

In *models M4* and *M5*, we hypothesized that the genetic component of height growth was influenced by the proportion belonging to each gene pool (proxy of the population demographic history and genetic drift). In *M4*, following Wolak and Reid (2017), gene pools $j$ were allowed to vary in their mean relative contribution $g_j$ on height growth as follows:

$$\log(h_{isbpg}) \sim \mathcal{N}(\mathbf{X}\beta + \mu_{sbpg}\ , \sigma^2)$$
$$\mathbf{X}\beta = \beta_0 + \beta_{age}\text{age}_i + \beta_{age2}\text{age}_i^2$$
$$\mu_{ijsbpg} = S_s + B_{b(s)} + P_p + G_{g(p)} + cs_{is} + \sum_{j=1}^{6} q_{gj} g_j$$
$$g_j \sim \mathcal{N}(0, \sigma_{g_j}^2)$$

where $q_{gj}$ corresponds to the proportion of each genotype $g$ belonging to the gene pool $j$ (as estimated in Jaramillo-Correa et al. 2015) and $g_j$ is the mean relative contribution of gene pool $j$ on height growth. In this model, trees from the same gene pool are considered to be unrelated, which is a reasonable assumption given the sampling scheme (see Materials & Methods).

*M5* extends *M4* by allowing gene pools $j$ to vary in their total genetic variance $\sigma_{A_j}^2$, following Muff et al. (2019). This involves replacing the genotype varying intercepts $G_{g(p)}$ in *M4* by partial genetic values $a_{gj}$ standing for the relative contribution of gene pool $j$ to the genetic value $a_g$ of genotype $g$ (approximated in *M4* by the genotype intercepts $G_{g(p)}$). Thus, *M5* can be expressed as *M4* but with $\mu_{ijsbpg}$ equal to:

$$\log(h_{isbpg}) \sim \mathcal{N}(\mathbf{X}\beta + \mu_{sbpg}\ , \sigma^2)$$
$$\mathbf{X}\beta = \beta_0 + \beta_{age}\text{age}_i + \beta_{age2}\text{age}_i^2$$
$$\mu_{ijsbpg} = S_s + B_{b(s)} + P_p + cs_{is} + \sum_{j=1}^{6} q_{gj} g_j + \sum_{j=1}^{6} a_{gj}$$
$$\mathbf{a}_j^{\intercal} = (a_{1j}, ..., a_{nj})^{\intercal} \sim \mathcal{N}(0, \sigma_{A_j}^2\ \mathbf{A_j})$$

with $\mathbf{A_j}$ the genomic relationship matrix specific to the gene pool $j$ and $\sigma_{A_j}^2$, the total genetic variance in gene pool $j$. $\mathbf{A_j}$ matrices were calculated based on SNPs that did not show any association with height at range-wide geographical scales (see Muff et al. 2019 for details on $\mathbf{A_j}$ calculation). Using the modeled residual variance $\sigma^2$ and gene-pool specific total genetic variances $\sigma_{A_j}^2$, we calculated the gene-pool specific broad-sense heritability as: $H_j^2 = \sigma_{A_j}^2/(\sigma_{A_j}^2 + \sigma^2)$.

In *model M6*, we hypothesized that populations are genetically adapted to the climatic conditions in which they evolved. Thus, we aimed to quantify the association between height growth and the climatic similarity among provenances, while accounting also for the proportion belonging to each gene pool. We kept the genotype varying intercepts (like in *M1* to *M4*) but not the gene pool-specific total genetic variances (unlike *M5*). Thus, *M6* extends *M4* as:

$$\log(h_{isbpg}) \sim \mathcal{N}(\mathbf{X}\beta + \mu_{sbpg}\ , \sigma^2)$$
$$\mathbf{X}\beta = \beta_0 + \beta_{age}\text{age}_i + \beta_{age2}\text{age}_i^2$$
$$\mu_{ijsbpg} = S_s + B_{b(s)} + P_p + G_{g(p)} + cs_{is} + cp_p + \sum_{j=1}^{6} q_{gj} g_j$$
$$cp_p \sim \mathcal{N}(0, \Phi\, \sigma_{cp_p}^2)$$

where $\Phi$ is the covariance matrix describing the climatic similarity between provenances $p$ (fig. S9) and $cp_p$ are varying intercepts associated with each provenance $p$. In *M6*, the genetic component was partitioned among the regression on the climatic covariates ($cp_p$), the gene pool covariates ($g_j$), and the deviations related to the genotype ($G_{g(p)}$) and provenance ($P_p$) effects (resulting, for instance, from adaptation to environmental variables not measured in our study).

##### 4.1.3 Predictive *models M7* to *M12*: combining climatic and genomic information to improve predictions

***Model M7***:

$$\log(h_{isb}) \sim \mathcal{N}(\mathbf{X}\beta + \mu_{jsbpg}\,,\,\sigma^2)$$
$$\mathbf{X}\beta = \beta_0 + \beta_{age}\text{age}_i + \beta_{age2}\text{age}_i^2$$
$$\mu_{jsbpg} = S_s + B_{b(s)} + \sum_{j=1}^{6} q_{gj}g_j + \beta_{min.pre,s}min.pre_p + \beta_{max.temp,s}max.temp_p + \beta_{gPEA,s}gPEA_g$$

***Model M8***:

$$\log(h_{isbr}) \sim \mathcal{N}(\mathbf{X}\beta + \mu_{jsbpgr}\,,\,\sigma^2)$$
$$\mathbf{X}\beta = \beta_0 + \beta_{age}\text{age}_i + \beta_{age2}\text{age}_i^2$$
$$\mu_{jsbpgr} = S_s + B_{b(s)} + \sum_{j=1}^{6} q_{gj}g_j + \beta_{min.pre,s}min.pre_p + \beta_{max.temp,s}max.temp_p + \beta_{rPEA,s}rPEA_{gr}$$

***Model M9***:

$$\log(h_{isb}) \sim \mathcal{N}(\mathbf{X}\beta + \mu_{jsb}\,,\,\sigma^2)$$
$$\mathbf{X}\beta = \beta_0 + \beta_{age}\text{age}_i + \beta_{age2}\text{age}_i^2$$
$$\mu_{jsb} = S_s + B_{b(s)} + \sum_{j=1}^{6} q_{gj}g_j$$

***Model M10***:

$$\log(h_{isb}) \sim \mathcal{N}(\mathbf{X}\beta + \mu_{jsbp}\,,\,\sigma^2)$$
$$\mathbf{X}\beta = \beta_0 + \beta_{age}\text{age}_i + \beta_{age2}\text{age}_i^2$$
$$\mu_{jsbp} = S_s + B_{b(s)} + \beta_{min.pre,s}min.pre_p + \beta_{max.temp,s}max.temp_p$$

***Model M11***:

$$\log(h_{isbg}) \sim \mathcal{N}(\mathbf{X}\beta + \mu_{sbg}\,,\,\sigma^2)$$
$$\mathbf{X}\beta = \beta_0 + \beta_{age}\text{age}_i + \beta_{age2}\text{age}_i^2$$
$$\mu_{sbg} = S_s + B_{b(s)} + \beta_{gPEA,s}gPEA_g$$

***Model M12***:

$$\log(h_{isbgr}) \sim \mathcal{N}(\mathbf{X}\beta + \mu_{sbgr}\,,\,\sigma^2)$$
$$\mathbf{X}\beta = \beta_0 + \beta_{age}\text{age}_i + \beta_{age2}\text{age}_i^2$$
$$\mu_{sbgr} = S_s + B_{b(s)} + \beta_{rPEA,s}rPEA_{gr}$$

where $min.pre_p$ and $max.temp_p$ are the climatic variables in the provenance locations, $\beta_{min.pre,s}$ and $\beta_{max.temp,s}$ their site-specific slopes, $gPEA_g$ and $rPEA_{gr}$ the counts of global and regional PEAs and $\beta_{gPEA,s}$ and $\beta_{rPEA,s}$ their site-specific slopes.

#### 4.2 Model priors

In **all models:**

$$\begin{bmatrix} S_s \\ B_{b(s)} \\ P_p \\ G_{g(p)} \\ S_s P_p \end{bmatrix} \sim \mathcal{N}\left(0, \begin{bmatrix} \sigma_S \\ \sigma_B \\ \sigma_P \\ \sigma_G \\ \sigma_{Inter} \end{bmatrix}\right)$$

$$(\sigma, \sigma_S, \sigma_B, \sigma_P, \sigma_G, \sigma_{Inter}, \sigma_{cs_{is}}, \sigma_{g_j}, \sigma_{A_j}, \sigma_{cp_p})^\intercal \sim \text{StudentT}(3, 0, 10)$$

$$\beta_0 \sim \mathcal{N}(0, 5)$$

$$\begin{bmatrix} \beta_{age} \\ \beta_{age2} \end{bmatrix} \sim \mathcal{N}(0, 1)$$

In ***model M7***:

$$\begin{bmatrix} S_s \\ \beta_{min.pre,s} \\ \beta_{max.temp,s} \\ \beta_{gPEA,s} \end{bmatrix} \sim \text{MVNormal}\left(\begin{bmatrix} 0 \\ 0 \end{bmatrix}, \mathbf{S}\right)$$

$$\mathbf{S} = \begin{pmatrix} \sigma_S & 0 & 0 & 0 \\ 0 & \sigma_{\beta_{min.pre,s}} & 0 & 0 \\ 0 & 0 & \sigma_{\beta_{max.temp,s}} & 0 \\ 0 & 0 & 0 & \sigma_{\beta_{gPEA,s}} \end{pmatrix} \begin{pmatrix} 1 & 1 & 1 & \rho \\ 1 & 1 & \rho & 1 \\ 1 & \rho & 1 & 1 \\ \rho & 1 & 1 & 1 \end{pmatrix} \begin{pmatrix} \sigma_S & 0 & 0 & 0 \\ 0 & \sigma_{\beta_{min.pre,s}} & 0 & 0 \\ 0 & 0 & \sigma_{\beta_{max.temp,s}} & 0 \\ 0 & 0 & 0 & \sigma_{\beta_{gPEA,s}} \end{pmatrix}$$

$$\begin{bmatrix} \sigma_S \\ \sigma_{\beta_{min.pre,s}} \\ \sigma_{\beta_{max.temp,s}} \\ \sigma_{\beta_{gPEA,s}} \end{bmatrix} \sim \text{StudentT}(3, 0, 10)$$

$$\begin{pmatrix} 1 & 1 & 1 & \rho \\ 1 & 1 & \rho & 1 \\ 1 & \rho & 1 & 1 \\ \rho & 1 & 1 & 1 \end{pmatrix} \sim \text{LKJcorr(4)}$$

where $\beta_{x,s}$ corresponds to $\beta_{min.pre,s}$ in M7 and $\beta_{max.temp,s}$ in M8.

In ***model M8***: same as M7 but replacing $\beta_{gPEA,s}$ and $\sigma_{\beta_{gPEA,s}}$ by $\beta_{rPEA,s}$ and $\sigma_{\beta_{rPEA,s}}$, respectively.

In ***models M9***, ***M10***, ***M11*** and ***M12***, same as M7 and M8.

### 5 Model comparison

#### 5.1 Description of the different indices used to compare the models

Similarly to the in-sample proportion of the variance explained by each model $m$ in each site $s$ ($\mathcal{R}^2_{ms}|age$), we calculated the out-of-sample proportion of the variance predicted by each model $m$ in each site $s$ conditional on the age effect as follows:

$$\textit{prediction}\ \mathcal{R}^2_{ms}|age = \frac{V_{\text{pred}_{ms}} - V_{\text{age}_{2s}}}{V_{\text{y}_s} - V_{\text{age}_{2s}}}$$

where $V_{\text{pred}_{ms}}$ is the variance of the modeled predictive means from model $m$ in site $s$ of the test dataset, $V_{\text{age}_{2s}}$ is the variance predicted by the age effect in the *model M2* in site $s$ and $V_{\text{y}_s}$ is the phenotypic variance in the site $s$ of the test dataset. Estimates of *prediction* $\mathcal{R}^2_{ms}|age$ in the three partitions are reported in Table 4.

We then calculated other indices that are not presented in the main manuscript as they are not necessary to support the main objectives of the paper, especially whether the models combining the climatic and genomic drivers of the genetic component can improve the prediction on new provenances. However, they are still useful to compare the goodness-of-fit and predictive ability of the models, that's why we report them here.

We calculated the total in-sample proportion of the variance explained by each model $m$ such as:

$$\mathcal{R}^2_m = \frac{V_{\text{pred}_m}}{V_\text{y}}$$

where $V_{\text{pred}_m}$ is the variance of the modeled predictive means from model $m$ in the training dataset and $V_\text{y}$ is the phenotypic variance in the training dataset. Similarly, we calculated the *prediction* $\mathcal{R}^2_m$ on the test dataset, that is the total out-of-sample proportion of variance predicted by each model $m$ in the test dataset.

We calculated the in-sample proportion of the variance explained by the fixed effects of each model $m$ such as:

$$\mathcal{R}^2_{m\,(\text{fix})} = \frac{V_{\text{pred}_m(\text{fix})}}{V_\text{y}}$$

where $V_{\text{pred}_m(\text{fix})}$ is the variance explained by the fixed effects of model $m$ in the training dataset and $V_\text{y}$ is the phenotypic variance in the training dataset. Similarly, we calculated the *prediction* $\mathcal{R}^2_m(fix)$ on the test dataset, that is the out-of-sample proportion of variance predicted by the fixed effects of each model $m$ in the test dataset.

Last, we calculated the total in-sample proportion of the variance explained by each model $m$ conditional on the age effect, such as:

$$\mathcal{R}^2_m|age = \frac{V_{\text{pred}_m} - V_{\text{age}_2}}{V_\text{y} - V_{\text{age}_2}}$$

where $V_{\text{age}_2}$ is the variance explained by the age effect in *model M2*. Similarly, we calculated the *prediction* $\mathcal{R}^2_m|age$ on the test dataset, that is the total out-of-sample proportion of variance predicted by each model $m$ conditional on the age effect in the test dataset.

To both evaluate the model goodness-of-fit and predictive ability, we also calculated the model **mean predictive error** of each model $m$ (mean of observed minus predicted responses, $\text{PE}_m$) on the training and test datasets of the three partitions.

$\mathcal{R}^2_m$, $\mathcal{R}^2_{m\,\text{(fix)}}$, $\mathcal{R}^2_m|age$, $\text{PE}_m$ and their predictive equivalents are presented in Tables S4 (P1 partition), S9 (P2 partition) and S12 (P3 partition).

To assess the model predictive ability, we also calculated the $\text{ELPD}_{loo}$, which is the Bayesian leave-one-out estimate of the expected log pointwise predictive density (equation 4 in Vehtari et al. 2017). This is a method for estimating out-of-sample prediction accuracy of Bayesian models, which is asymptotically equal to WAIC (Vehtari et al. 2017) and has the great advantage that it can be estimated without refitting the model. $\text{ELPD}_{loo}$ estimates can be found in Table S6 for the P1 partition and its pairwise comparisons between models in Tables S7 (P1 partition) and S10 (P2 partition). $\text{ELPD}_{loo}$, like WAIC, provides various advantages over AIC and DIC, especially that it is not a point estimate and, on the contrary, has an entire posterior distribution (Vehtari et al. 2017). Moreover, calculating $\text{ELPD}_{loo}$ using Pareto-smoothed importance sampling as we did in the present study using the *loo* R package, leads to more robust estimates than with WAIC (e.g. in cases with weak priors or influential observations). Models with higher $\text{ELPD}_{loo}$ are expected to have a higher predictive ability for new observations. In other words, $\text{ELPD}_{loo}$ indicates which model best captures each left-out data point. Therefore, $\text{ELPD}_{loo}$ indicates whether models have good predictive ability for new observations, but not for new groups (e.g., new provenances in our case). To estimate to predictive ability on new provenances with the $\text{ELPD}_{loo}$, we would have had to divide the dataset into $k$ partitions (e.g. 34 partitions and leaving one provenance out each time, 34 being the number of provenances; or 6 partitions and leaving $\sim$6 provenances out each time) and run the models $k$ times, which would have been very computationally heavy and was not feasible in our case given the computation time of some models (almost a week for *M5*). This is why we used the *prediction* $\mathcal{R}^2_{ms}|age$ instead of the $\text{ELPD}_{loo}$ to compare the models as it allowed us to calculate the variance explained and predicted by the models conditional on the age effect, and also to compare their predictive ability on new provenances without running the models again.

#### 5.2 In-sample proportion of variance explained conditional on age

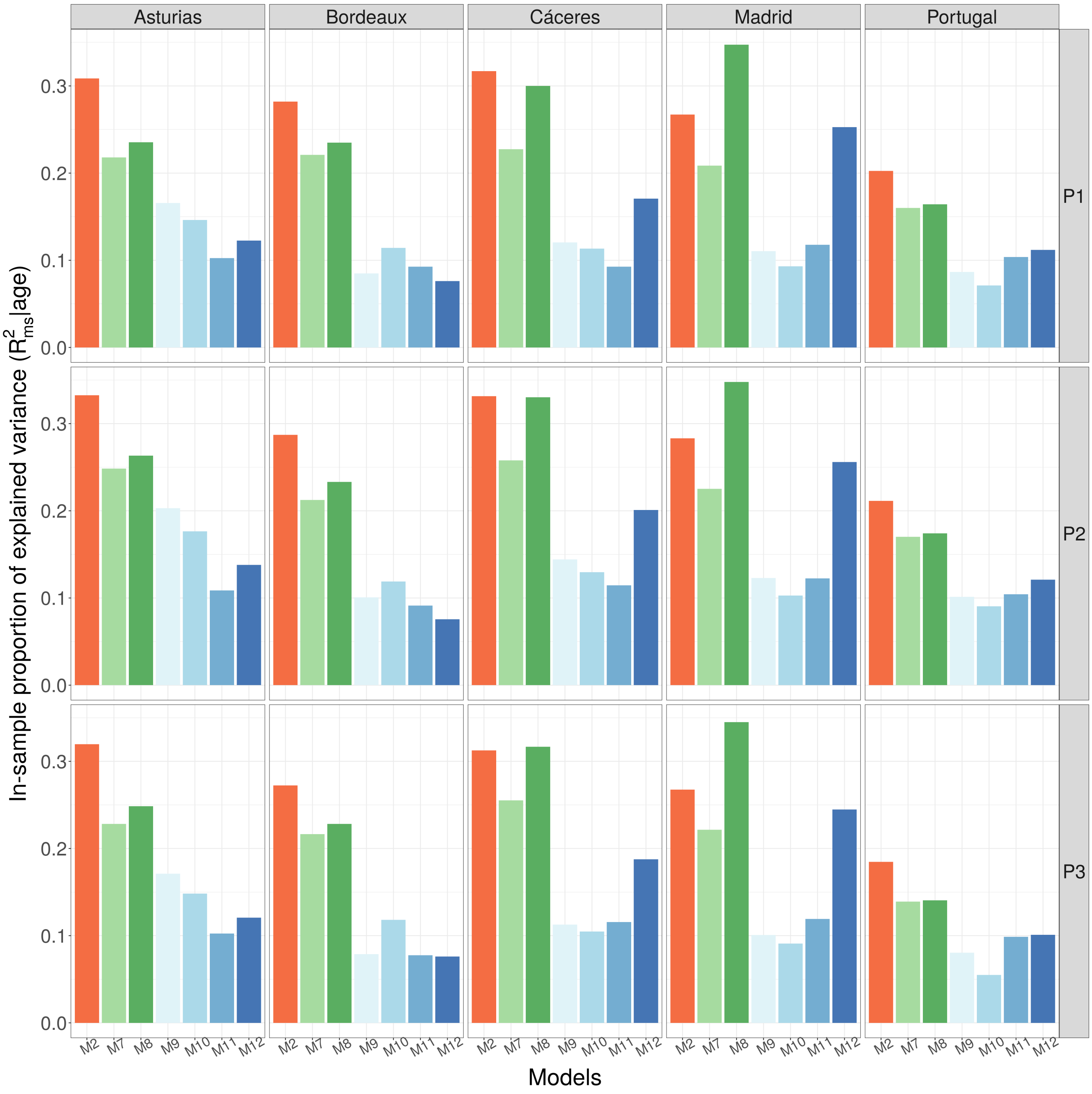

**Figure S10.** In-sample proportion of explained variance conditional on the age effect ($\mathcal{R}^2_{ms}|age$) in the training datasets (data used to fit the models) of the P1, P2 and P3 partitions. In the P1 partition, the training dataset was obtained by randomly sampling 75% of the observations and the test dataset contains the remaining 25% observations. In the P2 partition, the training dataset was obtained by randomly sampling 28 provenances and the test data set contains the remaining 6 provenances. The P3 partition corresponds to a non-random split between a training dataset of 28 provenances and a test dataset containing 6 provenances with at least one provenance from each under-represented gene pool. The exact values of the $\mathcal{R}^2_{ms}|age$ estimates and their associated credible intervals can be found in Tables S4 (P1 partition), S9 (P2 partition) and S12 (P3 partition).

#### 5.3 P1 partition (random split of the observations)

##### 5.3.1 Variance explained and predicted

| Models | **Explanatory part**: training P1 | | | | **Predictive part**: test P1 | | | |
|---|---|---|---|---|---|---|---|---|
| | $\mathcal{R}^2_m \vert age$ | $\mathcal{R}^2_m$ | $\mathcal{R}^2_m(fix)$ | PE$_m$ | prediction $\mathcal{R}^2_m \vert age$ | prediction $\mathcal{R}^2_m$ | prediction $\mathcal{R}^2_m(fix)$ | PE$_m$ |
| **M0** | 0.463 [0.439-0.487] | 0.768 [0.758-0.779] | 0.575 [0.564-0.586] | 0.267 [0.010-0.84] | 0.462 [0.437-0.487] | 0.773 [0.762-0.784] | 0.584 [0.573-0.596] | 0.269 [0.010-0.851] |
| **M1** | 0.571 [0.548-0.594] | 0.815 [0.805-0.825] | 0.569 [0.559-0.578] | 0.231 [0.008-0.754] | 0.559 [0.535-0.583] | 0.814 [0.804-0.824] | 0.578 [0.568-0.588] | 0.236 [0.008-0.769] |
| **M2** | 0.578 [0.556-0.601] | 0.818 [0.808-0.828] | 0.568 [0.559-0.578] | 0.229 [0.009-0.752] | 0.567 [0.543-0.591] | 0.817 [0.807-0.827] | 0.577 [0.568-0.587] | 0.235 [0.008-0.763] |
| **M3** | 0.574 [0.552-0.597] | 0.816 [0.807-0.826] | 0.566 [0.506-0.621] | 0.231 [0.009-0.747] | 0.563 [0.538-0.587] | 0.815 [0.805-0.825] | 0.575 [0.514-0.631] | 0.235 [0.009-0.763] |
| **M4** | 0.575 [0.553-0.597] | 0.817 [0.807-0.826] | 0.568 [0.507-0.623] | 0.230 [0.008-0.747] | 0.563 [0.54-0.587] | 0.815 [0.806-0.826] | 0.577 [0.516-0.634] | 0.235 [0.009-0.763] |
| **M5** | 0.573 [0.551-0.595] | 0.816 [0.806-0.825] | 0.568 [0.508-0.622] | 0.231 [0.008-0.751] | 0.562 [0.539-0.585] | 0.815 [0.805-0.825] | 0.577 [0.516-0.632] | 0.236 [0.009-0.767] |
| **M6** | 0.575 [0.553-0.598] | 0.817 [0.807-0.826] | 0.567 [0.504-0.624] | 0.230 [0.009-0.748] | 0.563 [0.539-0.587] | 0.816 [0.805-0.826] | 0.576 [0.512-0.635] | 0.235 [0.009-0.763] |
| **M7** | 0.546 [0.523-0.569] | 0.804 [0.794-0.814] | 0.570 [0.560-0.58] | 0.241 [0.009-0.788] | 0.529 [0.506-0.553] | 0.801 [0.792-0.811] | 0.580 [0.570-0.590] | 0.243 [0.008-0.790] |
| **M8** | 0.554 [0.532-0.577] | 0.807 [0.798-0.817] | 0.570 [0.560-0.580] | 0.238 [0.008-0.780] | 0.535 [0.512-0.558] | 0.804 [0.794-0.814] | 0.580 [0.569-0.590] | 0.239 [0.009-0.783] |
| **M9** | 0.502 [0.478-0.525] | 0.785 [0.775-0.795] | 0.574 [0.564-0.585] | 0.253 [0.010-0.822] | 0.491 [0.467-0.516] | 0.785 [0.775-0.796] | 0.584 [0.573-0.595] | 0.255 [0.009-0.826] |
| **M10** | 0.497 [0.473-0.520] | 0.783 [0.773-0.793] | 0.574 [0.564-0.585] | 0.255 [0.009-0.821] | 0.485 [0.460-0.509] | 0.783 [0.772-0.793] | 0.584 [0.573-0.594] | 0.257 [0.010-0.828] |
| **M11** | 0.500 [0.477-0.523] | 0.784 [0.774-0.794] | 0.571 [0.561-0.581] | 0.257 [0.010-0.812] | 0.495 [0.471-0.519] | 0.787 [0.777-0.797] | 0.580 [0.570-0.591] | 0.259 [0.009-0.831] |
| **M12** | 0.506 [0.482-0.530] | 0.787 [0.776-0.797] | 0.571 [0.560-0.581] | 0.255 [0.009-0.804] | 0.498 [0.473-0.523] | 0.788 [0.778-0.799] | 0.580 [0.570-0.591] | 0.257 [0.011-0.825] |

**Table S4:** Summary of model performance in the P1 partition, in which the training dataset was obtained by randomly sampling 75% of the observations and the test dataset contains the remaining 25% observations. $\mathcal{R}^2_m|age$ and *prediction* $\mathcal{R}^2_m|age$ correspond to the proportion of variance explained and predicted by the models conditional on the age effect in the training (in-sample) and test (out-of-sample) datasets, respectively. $\mathcal{R}^2_m$ and $\mathcal{R}^2_m(fix)$ correspond to the proportion of explained variance in the training dataset by the entire model and by the fixed variables only, respectively. *Prediction* $\mathcal{R}^2_m$ and $\mathcal{R}^2_m(fix)$ correspond to the proportion of variance in the test dataset predicted by the entire model or by the fixed effects, respectively. PE$_m$ corresponds to the mean predictive error (mean of observed minus predicted responses). For $\mathcal{R}^2$ and PE, the mean (over all iterations for $\mathcal{R}^2$ or over all observations for PE) and the 95% credible intervals are given. For more details on the calculation of each index, see section 5.1 of the Supplementary Information. See Table 1 and main text for description of model the components.

*M0* explained less variance (lower $\mathcal{R}^2_m$ and $\mathcal{R}^2_m|age$) than the other models as it did not account for the genetic component of height growth (Table S4). The models that accounted for the genetic component with varying intercepts for the provenances explained 81.5% of the variance (Table S4). The models combining genomic (PEAs and gene pools) and climatic drivers (*M7* and *M8*) explained more variance (higher $\mathcal{R}^2_m$ and $\mathcal{R}^2_m|age$) than models including separately each driver (*M9* to *M12*). $\mathcal{R}^2_m$ and *prediction* $\mathcal{R}^2_m$ were similar for all models, meaning that the models predicted well new observations (i.e. new observations but from the same sites and same provenances).

##### 5.3.2 Variance partitioning conditional on the age effect

We examine here the partitioning of the variance conditional on the age effect, which provides insight into the genetic and plastic components of deviations from the mean height-growth trajectory (Table S5). It was not possible to extract the variance explained by the different drivers of the genetic and plastic components (i.e. for the plastic component, the variance associated with site intercepts and the intercepts associated with climatic similarity between sites) as they were confounded, and thus their explained variance could not be disentangled.

| Models | Environment | Genetic | Genetic x Environment | Residuals |
|---|---|---|---|---|
| **M0** | 45.5% [43.7 - 47.2] | - | - | 54.6% [53.2 - 56.0] |
| **M1** | 46.5% [44.9 - 48.0] | 11.4% [10.7 - 12.1] | - | 42.9% [41.9 - 43.9] |
| **M2** | 46.8% [42.5 - 51.2] | 11.0% [9.6 - 12.5] | 1.5% [-1.0 - 4.3] | 42.2% [41.3 - 43.1] |

**Table S5:** Partitioning of the variance explained on the data used for sampling conditional on the age effect (i.e. the variance of the deviations from the mean height-growth trajectory).

##### 5.3.3 Bayesian LOO estimate of the expected log predictive density (ELPD$_{loo}$)

| Models | ELPD$_{loo}$ |
|---|---|
| M0 | -9216 [165] |
| M1 | -6652 [182] |
| M2 | -6487 [182] |
| M3 | -6551 [184] |
| M4 | -6549 [184] |
| M5 | -6591 [184] |
| M6 | -6550 [184] |
| M7 | -7135 [179] |
| M8 | -6926 [180] |
| M9 | -8276 [172] |
| M10 | -8405 [173] |
| M11 | -8334 [169] |
| M12 | -8171 [169] |

**Table S6:** ELPD$_{loo}$ of models fitted on the training dataset of the P1 partition. The mean and the standard deviation (in brackets) are given.

| | M1 | M10 | M11 | M12 | M2 | M3 | M4 | M5 | M6 | M7 | M8 | M9 |
|---|---|---|---|---|---|---|---|---|---|---|---|---|
| M0 | **-2563.62 [71.79]** | **-810.25 [39.04]** | **-881.82 [41.73]** | **-1044.94 [45.56]** | **-2728.44 [73.82]** | **-2665.02 [73.19]** | **-2666.62 [73.22]** | **-2624.87 [73.01]** | **-2665.94 [73.18]** | **-2080.78 [63.39]** | **-2289.82 [66.96]** | **-939.27 [42.76]** |
| M1 | | **1753.37 [61.78]** | **1681.8 [61.36]** | **1518.68 [62.97]** | **-164.82 [18.35]** | **-101.4 [14.94]** | **-103 [14.96]** | -61.24 [17.18] | **-102.32 [14.92]** | **482.84 [40.42]** | **273.8 [45.04]** | **1624.35 [59.03]** |
| M10 | | | -71.57 [58.87] | -234.69 [61.4] | **-1918.19 [64.12]** | **-1854.77 [63.52]** | **-1856.37 [63.39]** | **-1814.61 [63.14]** | **-1855.68 [63.31]** | **-1270.53 [51.27]** | **-1479.56 [55.64]** | **-129.01 [21.51]** |
| M11 | | | | **-163.12 [31.09]** | **-1846.62 [63.88]** | **-1783.2 [62.89]** | **-1784.8 [63.1]** | **-1743.04 [62.64]** | **-1784.12 [63.05]** | **-1198.96 [47.55]** | **-1408 [58.33]** | -57.45 [62.64] |
| M12 | | | | | **-1683.5 [63.52]** | **-1620.08 [64.65]** | **-1621.68 [64.92]** | **-1579.93 [64.43]** | **-1621 [64.89]** | **-1035.84 [56.61]** | **-1244.88 [48.18]** | 105.67 [64.94] |
| M2 | | | | | | 63.42 [24.36] | 61.82 [24.4] | **103.58 [25.8]** | 62.51 [24.39] | **647.66 [44.45]** | **438.63 [45.28]** | **1789.18 [61.98]** |
| M3 | | | | | | | -1.6 [1.24] | **40.16 [8.61]** | -0.91 [1.09] | **584.24 [43.19]** | **375.21 [47.81]** | **1725.76 [61.02]** |
| M4 | | | | | | | | **41.76 [8.52]** | 0.68 [0.78] | **585.84 [43.06]** | **376.8 [47.78]** | **1727.35 [60.75]** |
| M5 | | | | | | | | | **-41.07 [8.54]** | **544.09 [42.25]** | **335.05 [47]** | **1685.6 [60.47]** |
| M6 | | | | | | | | | | **585.16 [43.02]** | **376.12 [47.77]** | **1726.67 [60.72]** |
| M7 | | | | | | | | | | | **-209.04 [35]** | **1141.51 [48.56]** |
| M8 | | | | | | | | | | | | **1350.55 [52.98]** |

**Table S7:** ELPD$_{loo}$ differences among models fitted on the training dataset of the P1 partition. The ELPD$_{loo}$ difference among models corresponds to the ELPD$_{loo}$ of the row model minus the ELPD$_{loo}$ of the column model. Thus, a negative difference in ELPD$_{loo}$ means that the row model has a lower ELPD$_{loo}$ than the column model, and therefore a lower predictive ability on new observations. The standard error of the model differences is indicated between brackets. Two models are considered significantly different when their absolute ELPD$_{loo}$ difference is higher than four times the standard error (in bold).

##### 5.3.4 Site-specific predicted variance conditional on the age effect

| Models | Asturias | Bordeaux | Cáceres | Madrid | Portugal |
|---|---|---|---|---|---|
| **M0** | 0.064 [-0.027-0.155] | 0.002 [-0.02-0.024] | 0.017 [0.004-0.043] | 0.035 [0.015-0.063] | 0.029 [0.007-0.051] |
| **M1** | 0.341 [0.257-0.43] | 0.215 [0.192-0.239] | 0.227 [0.196-0.263] | 0.204 [0.18-0.231] | 0.161 [0.139-0.184] |
| **M2** | 0.349 [0.264-0.436] | 0.259 [0.229-0.289] | 0.266 [0.201-0.342] | 0.25 [0.208-0.297] | 0.171 [0.148-0.194] |
| **M3** | 0.242 [0.142-0.341] | 0.362 [0.324-0.401] | 0.227 [0.197-0.262] | 0.203 [0.18-0.23] | 0.168 [0.141-0.198] |
| **M4** | 0.244 [0.144-0.346] | 0.363 [0.324-0.403] | 0.228 [0.197-0.265] | 0.203 [0.179-0.23] | 0.169 [0.14-0.198] |
| **M5** | 0.246 [0.15-0.344] | 0.357 [0.32-0.397] | 0.227 [0.195-0.263] | 0.2 [0.176-0.228] | 0.166 [0.138-0.195] |
| **M6** | 0.243 [0.144-0.342] | 0.363 [0.324-0.402] | 0.228 [0.197-0.265] | 0.203 [0.179-0.231] | 0.169 [0.14-0.199] |
| **M7** | 0.251 [0.168-0.336] | 0.186 [0.157-0.215] | 0.184 [0.112-0.276] | 0.158 [0.117-0.205] | 0.122 [0.101-0.145] |
| **M8** | 0.248 [0.163-0.332] | 0.2 [0.171-0.23] | 0.235 [0.147-0.344] | 0.337 [0.269-0.414] | 0.137 [0.114-0.159] |
| **M9** | 0.167 [0.082-0.256] | 0.071 [0.05-0.092] | 0.093 [0.079-0.116] | 0.093 [0.075-0.117] | 0.063 [0.042-0.084] |
| **M10** | 0.124 [0.037-0.211] | 0.093 [0.067-0.12] | 0.08 [0.029-0.148] | 0.086 [0.052-0.126] | 0.052 [0.031-0.074] |
| **M11** | 0.129 [0.045-0.215] | 0.074 [0.049-0.098] | 0.071 [0.024-0.138] | 0.102 [0.066-0.143] | 0.087 [0.064-0.109] |
| **M12** | 0.124 [0.04-0.21] | 0.063 [0.038-0.088] | 0.128 [0.061-0.219] | 0.262 [0.196-0.335] | 0.102 [0.079-0.124] |

**Table S8:** Summary table of site-specific proportion of predicted variance conditional on the age effect (prediction $\mathcal{R}^2_{ms}|age$) in the test datasets (data not used to fit the models) of the P1 partition. The numbers shown are the mean and the 95% credible intervals.

#### 5.4 P2 partition (random split of the provenances)

##### 5.4.1 Variance explained and predicted

| Models | **Explanatory part**: training P2 | | | | **Predictive part**: test P2 | | | |
|---|---|---|---|---|---|---|---|---|
| | $\mathcal{R}^2_m|age$ | $\mathcal{R}^2_m$ | $\mathcal{R}^2_m(fix)$ | PE$_m$ | prediction $\mathcal{R}^2_m|age$ | prediction $\mathcal{R}^2_m$ | prediction $\mathcal{R}^2_m(fix)$ | PE$_m$ |
| **M0** | 0.454 [0.431-0.478] | 0.766 [0.756-0.776] | 0.577 [0.567-0.587] | 0.268 [0.011-0.839] | 0.425 [0.403-0.447] | 0.745 [0.736-0.755] | 0.563 [0.553-0.573] | 0.263 [0.010-0.881] |
| **M1** | 0.571 [0.550-0.592] | 0.816 [0.807-0.825] | 0.571 [0.562-0.581] | 0.230 [0.008-0.746] | 0.428 [0.409-0.448] | 0.747 [0.738-0.755] | 0.557 [0.548-0.566] | 0.264 [0.010-0.877] |
| **M2** | 0.578 [0.558-0.599] | 0.819 [0.810-0.828] | 0.571 [0.562-0.58] | 0.228 [0.009-0.743] | 0.426 [0.374-0.482] | 0.746 [0.723-0.770] | 0.556 [0.548-0.565] | 0.264 [0.010-0.870] |
| **M7** | 0.546 [0.524-0.567] | 0.805 [0.796-0.814] | 0.574 [0.564-0.583] | 0.239 [0.008-0.779] | 0.502 [0.481-0.522] | 0.779 [0.770-0.788] | 0.559 [0.550-0.568] | 0.247 [0.008-0.861] |
| **M8** | 0.554 [0.532-0.575] | 0.809 [0.799-0.818] | 0.574 [0.564-0.583] | 0.237 [0.009-0.771] | 0.540 [0.518-0.562] | 0.796 [0.786-0.806] | 0.559 [0.550-0.569] | 0.246 [0.008-0.844] |
| **M9** | 0.505 [0.483-0.528] | 0.788 [0.778-0.798] | 0.578 [0.568-0.587] | 0.251 [0.009-0.805] | 0.479 [0.458-0.501] | 0.769 [0.760-0.779] | 0.563 [0.554-0.573] | 0.264 [0.009-0.906] |
| **M10** | 0.499 [0.476-0.522] | 0.785 [0.775-0.795] | 0.577 [0.567-0.587] | 0.252 [0.009-0.808] | 0.492 [0.470-0.514] | 0.775 [0.765-0.785] | 0.563 [0.553-0.573] | 0.265 [0.009-0.905] |
| **M11** | 0.493 [0.470-0.516] | 0.783 [0.773-0.793] | 0.573 [0.563-0.583] | 0.257 [0.009-0.813] | 0.471 [0.449-0.493] | 0.766 [0.756-0.776] | 0.559 [0.549-0.569] | 0.258 [0.010-0.856] |
| **M12** | 0.503 [0.480-0.526] | 0.787 [0.777-0.797] | 0.573 [0.563-0.583] | 0.254 [0.010-0.802] | 0.522 [0.499-0.545] | 0.788 [0.778-0.798] | 0.559 [0.549-0.568] | 0.263 [0.010-0.833] |

**Table S9:** Summary of model performance in the P2 partition, in which the training dataset was obtained by randomly sampling 28 provenances and the test data set contains the remaining 6 provenances. $\mathcal{R}^2_m|age$ and prediction $\mathcal{R}^2_m|age$ correspond to the proportion of variance explained and predicted by the models conditional on the age effect in the training (in-sample) and test (out-of-sample) datasets, respectively. $\mathcal{R}^2_m$ and $\mathcal{R}^2_m(fix)$ correspond to the proportion of explained variance in the training dataset by the entire model and by the fixed variables only, respectively. Prediction $\mathcal{R}^2_m$ and $\mathcal{R}^2_m(fix)$ correspond to the proportion of variance in the test dataset predicted by the entire model or by the fixed effects, respectively. PE$_m$ corresponds to the mean predictive error (mean of observed minus predicted responses). For $\mathcal{R}^2$ and PE, the mean (over all iterations for $\mathcal{R}^2$ or over all observations for PE) and the 95% credible intervals are given. For more details on the calculation of each index, see section 5 of the Supplementary Information. See Table 1 and main text for description of model the components.

##### 5.4.2 Bayesian LOO estimate of the expected log predictive density (ELPD$_{loo}$)

| | M1 | M10 | M11 | M12 | M2 | M7 | M8 | M9 |
|---|---|---|---|---|---|---|---|---|
| M0 | **-3097.43 [80.4]** | **-1180.71 [47.7]** | **-1001.98 [45.13]** | **-1293 [50.8]** | **-3307.11 [82.58]** | **-2503.89 [70.49]** | **-2740.39 [74.19]** | **-1339.65 [50.55]** |
| M1 | | **1916.72 [66.85]** | **2095.45 [68.57]** | **1804.43 [69.11]** | **-209.68 [20.62]** | **593.55 [43.39]** | **357.04 [49.06]** | **1757.78 [63.01]** |
| M10 | | | 178.73 [65.95] | -112.29 [67.87] | **-2126.4 [69.47]** | **-1323.17 [54.04]** | **-1559.68 [58.5]** | **-158.94 [25.24]** |
| M11 | | | | **-291.02 [34.88]** | b | **-1501.91 [53.72]** | **-1738.41 [64.64]** | **-337.67 [69]** |
| M12 | | | | | **-2014.11 [69.7]** | **-1210.89 [60.93]** | **-1447.39 [52.32]** | -46.65 [70.45] |
| M2 | | | | | | **803.22 [48.2]** | **566.72 [49.33]** | **1967.46 [66.09]** |
| M7 | | | | | | | **-236.51 [35.87]** | **1164.23 [50.25]** |
| M8 | | | | | | | | **1400.74 [54.56]** |

**Table S10:** ELPD$_{loo}$ differences among models fitted on the training dataset of the P2 partition. The ELPD$_{loo}$ difference among models corresponds to the ELPD$_{loo}$ of the row model minus the ELPD$_{loo}$ of the column model. Thus, a negative difference in ELPD$_{loo}$ means that the row model has a lower ELPD$_{loo}$ than the column model, and therefore a lower predictive ability on new observations. The standard error of the model differences is indicated between brackets. Two models are considered significantly different when their absolute ELPD$_{loo}$ difference is higher than four times the standard error (in bold). Detail about interpretation of ELPD$_{loo}$ differences can be found in section 5.3.3.

##### 5.4.3 Site-specific predicted variance conditional on the age effect

| Models | Asturias | Bordeaux | Cáceres | Madrid | Portugal |
|---|---|---|---|---|---|
| **M0** | 0.052 [-0.024-0.128] | -0.001 [-0.022-0.021] | 0.035 [0.007-0.088] | 0.042 [0.017-0.076] | 0.033 [0.013-0.053] |
| **M2** | 0.005 [-0.061-0.074] | 0.004 [-0.016-0.024] | 0.035 [0.008-0.086] | 0.034 [0.015-0.062] | 0.017 [0.000-0.035] |
| **M1** | 0.011 [-0.056-0.079] | 0.004 [-0.015-0.023] | 0.035 [0.008-0.082] | 0.036 [0.016-0.063] | 0.019 [0.002-0.036] |
| **M7** | 0.149 [0.079-0.219] | 0.140 [0.114-0.167] | 0.247 [0.150-0.371] | 0.229 [0.170-0.299] | 0.111 [0.092-0.131] |
| **M8** | 0.124 [0.054-0.196] | 0.166 [0.137-0.195] | 0.290 [0.182-0.422] | 0.356 [0.281-0.439] | 0.101 [0.082-0.121] |
| **M9** | 0.166 [0.093-0.240] | 0.101 [0.078-0.123] | 0.162 [0.132-0.212] | 0.139 [0.115-0.169] | 0.076 [0.056-0.095] |
| **M10** | 0.178 [0.100-0.252] | 0.152 [0.120-0.185] | 0.204 [0.090-0.358] | 0.131 [0.085-0.187] | 0.081 [0.061-0.101] |
| **M11** | 0.100 [0.025-0.177] | 0.100 [0.072-0.129] | 0.189 [0.081-0.330] | 0.209 [0.138-0.292] | 0.103 [0.082-0.125] |
| **M12** | 0.141 [0.065-0.217] | 0.070 [0.044-0.095] | 0.274 [0.150-0.434] | 0.329 [0.253-0.413] | 0.131 [0.109-0.152] |

**Table S11:** Summary table of site-specific proportion of predicted variance conditional on the age effect (prediction $\mathcal{R}^2_{ms}|age$) in the test datasets (data not used to fit the models) of the P2 partition. The numbers shown are the mean and the 95% credible intervals.

#### 5.5 P3 partition (non-random split of the provenances)

##### 5.5.1 Variance explained and predicted

| Models | **Explanatory part**: training P3 | | | | **Predictive part**: test P3 | | | |
|---|---|---|---|---|---|---|---|---|
| | $\mathcal{R}^2_m\|age$ | $\mathcal{R}^2_m$ | $\mathcal{R}^2_m(fix)$ | PE$_m$ | prediction $\mathcal{R}^2_m\|age$ | prediction $\mathcal{R}^2_m$ | prediction $\mathcal{R}^2_m(fix)$ | PE$_m$ |
| **M0** | 0.448 [0.425-0.472] | 0.765 [0.755-0.775] | 0.58 [0.569-0.591] | 0.266 [0.01-0.842] | 0.382 [0.362-0.402] | 0.716 [0.706-0.725] | 0.545 [0.535-0.555] | 0.27 [0.009-0.853] |
| **M1** | 0.557 [0.535-0.579] | 0.811 [0.802-0.821] | 0.575 [0.565-0.584] | 0.232 [0.008-0.759] | 0.383 [0.365-0.402] | 0.716 [0.708-0.725] | 0.54 [0.531-0.549] | 0.271 [0.011-0.841] |
| **M2** | 0.563 [0.54-0.584] | 0.814 [0.804-0.823] | 0.574 [0.564-0.584] | 0.23 [0.008-0.753] | 0.388 [0.343-0.438] | 0.719 [0.698-0.741] | 0.539 [0.531-0.549] | 0.271 [0.011-0.842] |
| **M7** | 0.53 [0.507-0.553] | 0.8 [0.79-0.81] | 0.577 [0.566-0.587] | 0.241 [0.009-0.792] | 0.439 [0.415-0.464] | 0.742 [0.731-0.753] | 0.542 [0.532-0.551] | 0.242 [0.008-0.81] |
| **M8** | 0.537 [0.514-0.56] | 0.803 [0.793-0.813] | 0.577 [0.567-0.587] | 0.24 [0.009-0.784] | 0.504 [0.478-0.532] | 0.772 [0.76-0.785] | 0.542 [0.533-0.551] | 0.237 [0.009-0.78] |
| **M9** | 0.487 [0.464-0.511] | 0.782 [0.772-0.792] | 0.58 [0.57-0.591] | 0.253 [0.01-0.825] | 0.406 [0.385-0.428] | 0.727 [0.717-0.737] | 0.546 [0.536-0.555] | 0.286 [0.011-0.905] |
| **M10** | 0.48 [0.457-0.503] | 0.779 [0.769-0.789] | 0.581 [0.571-0.591] | 0.256 [0.01-0.827] | 0.394 [0.373-0.414] | 0.721 [0.712-0.731] | 0.546 [0.536-0.555] | 0.257 [0.01-0.807] |
| **M11** | 0.484 [0.46-0.508] | 0.781 [0.77-0.791] | 0.576 [0.566-0.587] | 0.257 [0.009-0.818] | 0.404 [0.384-0.425] | 0.726 [0.717-0.735] | 0.542 [0.532-0.551] | 0.258 [0.01-0.825] |
| **M12** | 0.49 [0.466-0.513] | 0.783 [0.773-0.793] | 0.576 [0.566-0.587] | 0.255 [0.01-0.809] | 0.442 [0.421-0.462] | 0.743 [0.734-0.753] | 0.542 [0.532-0.552] | 0.255 [0.009-0.819] |

**Table S12:** Summary of model performance in the P3 partition, in which the training dataset contains 28 provenances and the test dataset contains the remaining 6 provenances. $\mathcal{R}^2_m|age$ and prediction $\mathcal{R}^2_m|age$ correspond to the proportion of variance explained and predicted by the models conditional on the age effect in the training (in-sample) and test (out-of-sample) datasets, respectively. $\mathcal{R}^2_m$ and $\mathcal{R}^2_m(fix)$ correspond to the proportion of explained variance in the training dataset by the entire model and by the fixed variables only, respectively. Prediction $\mathcal{R}^2_m$ and $\mathcal{R}^2_m(fix)$ correspond to the proportion of variance in the test dataset predicted by the entire model or by the fixed effects, respectively. PE$_m$ corresponds to the mean predictive error (mean of observed minus predicted responses). For $\mathcal{R}^2$ and PE, the mean (over all iterations for $\mathcal{R}^2$ or over all observations for PE) and the 95% credible intervals are given. For more details on the calculation of each index, see section 5 of the Supplementary Information. See Table 1 and main text for description of model the components.

##### 5.5.2 Site-specific predicted variance conditional on the age effect

| Models | Asturias | Bordeaux | Cáceres | Madrid | Portugal |
|---|---|---|---|---|---|
| **M0** | 0.045 [-0.028-0.117] | 0.000 [-0.024-0.025] | 0.021 [0.005-0.048] | 0.033 [0.014-0.059] | 0.029 [0.010-0.048] |
| **M1** | 0.010 [-0.054-0.074] | 0.004 [-0.017-0.025] | 0.018 [0.004-0.044] | 0.024 [0.010-0.044] | 0.016 [-0.001-0.034] |
| **M2** | 0.006 [-0.057-0.071] | 0.004 [-0.017-0.026] | 0.019 [0.004-0.046] | 0.023 [0.009-0.043] | 0.015 [-0.002-0.032] |
| **M7** | 0.188 [0.110-0.266] | 0.246 [0.193-0.305] | 0.271 [0.160-0.409] | 0.205 [0.145-0.271] | 0.124 [0.097-0.152] |
| **M8** | 0.277 [0.197-0.360] | 0.310 [0.254-0.371] | 0.250 [0.152-0.369] | 0.223 [0.176-0.276] | 0.164 [0.134-0.197] |
| **M9** | 0.107 [0.035-0.180] | 0.061 [0.032-0.092] | 0.105 [0.074-0.148] | 0.096 [0.068-0.129] | 0.062 [0.041-0.084] |
| **M10** | 0.125 [0.056-0.195] | 0.146 [0.113-0.18] | 0.111 [0.040-0.209] | 0.075 [0.043-0.114] | 0.053 [0.034-0.073] |
| **M11** | 0.061 [-0.010-0.130] | 0.066 [0.04-0.092] | 0.080 [0.033-0.146] | 0.096 [0.063-0.134] | 0.062 [0.042-0.081] |
| **M12** | 0.061 [-0.007-0.132] | 0.095 [0.066-0.126] | 0.061 [0.032-0.102] | 0.121 [0.091-0.154] | 0.060 [0.041-0.080] |

**Table S13:** Summary table of site-specific proportion of predicted variance conditional on the age effect (prediction $\mathcal{R}^2_{ms}|age$) in the test datasets (data not used to fit the models) of the P3 partition. The numbers shown are the mean and the 95% credible intervals.

### 6 Posterior distributions and parameter interpretation

#### 6.1 P1 partition (random split of the observations)

##### 6.1.1 Baseline *models M0, M1* and *M2*: separating the genetic and plastic components

In the baseline model *M0*, only the plastic component is included (via the site and block intercepts). The genetic component is not considered (no intercepts for the provenances and the genotypes). *M0* was performed to compare the gain in explanatory and predictive power of models that account for the genetic component, compared to the model *M0* that does not.

***MODEL M0***

| Parameter | Median | SD | InfCI | SupCI |
|---|---|---|---|---|
| $\sigma^2_S$ | 0.105 | 0.370 | 0.027 | 0.979 |
| $\sigma^2_B$ | 0.002 | 0.001 | 0.001 | 0.004 |
| $\sigma^2$ | 0.123 | 0.001 | 0.121 | 0.125 |
| $\beta_0$ | 6.273 | 0.176 | 5.942 | 6.664 |
| $\beta_{age}$ | 0.601 | 0.003 | 0.596 | 0.607 |
| $\beta_{age2}$ | -0.153 | 0.003 | -0.159 | -0.147 |

**Table S14:** Parameter estimates of the varying-intercept variances ($\sigma^2_S$, $\sigma^2_B$), the global variance $\sigma^2$, the global intercept $\beta_0$ and the slopes associated with the age effect ($\beta_{age}$ and $\beta_{age2}$) in *M0*. SD corresponds to the standard deviation and InfCI and SupCI correspond to the lower and upper bounds of the 0.95 credible interval.

***MODEL M1***

In *M1*, the plastic component was mainly attributed to the variance $\sigma^2_S$ between sites (median of 0.108), while the variance $\sigma^2_B$ between blocks was almost null (median of 0.002) (Table S15). Some sites showed heights deviating strongly from the global mean: Madrid, where trees grew the least (median of -0.376), and Asturias where they grew particularly well (median of 0.272) (fig. 3 & Table S16). The genetic component was equally attributed to the variance between provenances $\sigma^2_P$ and genotypes $\sigma^2_G$, with a median of 0.013 and 0.012, respectively (Table S15).

| Parameter | Median | SD | InfCI | SupCI |
|---|---|---|---|---|
| $\sigma^2_P$ | 0.013 | 0.004 | 0.008 | 0.023 |
| $\sigma^2_G$ | 0.012 | 0.001 | 0.010 | 0.013 |
| $\sigma^2_S$ | 0.108 | 0.444 | 0.028 | 1.046 |
| $\sigma^2_B$ | 0.002 | 0.001 | 0.001 | 0.003 |
| $\sigma^2$ | 0.098 | 0.001 | 0.096 | 0.100 |
| $\beta_0$ | 6.247 | 0.193 | 5.842 | 6.643 |
| $\beta_{age}$ | 0.598 | 0.003 | 0.592 | 0.603 |
| $\beta_{age2}$ | -0.150 | 0.003 | -0.155 | -0.145 |

**Table S15:** Parameter estimates of the varying-intercept variances ($\sigma^2_S$, $\sigma^2_B$, $\sigma^2_P$ and $\sigma^2_G$), the global variance $\sigma^2$, the global intercept $\beta_0$ and the slopes associated with the age effect ($\beta_{age}$ and $\beta_{age2}$) in *M1*.

| Parameter | Median | SD | InfCI | SupCI |
|---|---|---|---|---|
| $S_{Asturias}$ | 0.292 | 0.193 | -0.103 | 0.690 |
| $S_{Bordeaux}$ | 0.151 | 0.193 | -0.254 | 0.555 |
| $S_{Caceres}$ | 0.041 | 0.194 | -0.366 | 0.443 |
| $S_{Madrid}$ | -0.376 | 0.193 | -0.780 | 0.019 |
| $S_{Portugal}$ | -0.110 | 0.193 | -0.513 | 0.288 |

**Table S16:** Parameter estimates of the site intercepts $S_s$ in *M1*. SD corresponds to the standard deviation and InfCI and SupCI correspond to the lower and upper bounds of the 0.95 credible interval.

| Parameter | Median | SD | InfCI | SupCI |
|---|---|---|---|---|
| $P_{ALT}$ | 0.089 | 0.041 | 0.009 | 0.168 |
| $P_{ARM}$ | 0.121 | 0.043 | 0.039 | 0.205 |
| $P_{ARN}$ | -0.093 | 0.032 | -0.158 | -0.031 |
| $P_{BAY}$ | -0.135 | 0.033 | -0.196 | -0.072 |
| $P_{BON}$ | -0.040 | 0.042 | -0.121 | 0.041 |
| $P_{CAD}$ | 0.061 | 0.040 | -0.016 | 0.142 |
| $P_{CAR}$ | -0.162 | 0.047 | -0.254 | -0.071 |
| $P_{CAS}$ | 0.054 | 0.041 | -0.026 | 0.132 |
| $P_{CEN}$ | 0.011 | 0.042 | -0.073 | 0.093 |
| $P_{COC}$ | -0.154 | 0.034 | -0.222 | -0.088 |
| $P_{COM}$ | 0.076 | 0.054 | -0.033 | 0.181 |
| $P_{CUE}$ | -0.171 | 0.030 | -0.233 | -0.113 |
| $P_{HOU}$ | 0.162 | 0.030 | 0.102 | 0.220 |
| $P_{LAM}$ | 0.014 | 0.040 | -0.064 | 0.093 |
| $P_{LEI}$ | 0.038 | 0.032 | -0.023 | 0.101 |
| $P_{MAD}$ | 0.021 | 0.083 | -0.146 | 0.181 |
| $P_{MIM}$ | 0.005 | 0.034 | -0.061 | 0.071 |
| $P_{OLB}$ | 0.047 | 0.031 | -0.013 | 0.108 |
| $P_{OLO}$ | 0.141 | 0.032 | 0.078 | 0.204 |
| $P_{ORI}$ | -0.121 | 0.030 | -0.180 | -0.062 |
| $P_{PET}$ | 0.123 | 0.030 | 0.065 | 0.182 |
| $P_{PIA}$ | 0.089 | 0.034 | 0.023 | 0.154 |
| $P_{PIE}$ | -0.078 | 0.042 | -0.163 | 0.002 |
| $P_{PLE}$ | 0.000 | 0.032 | -0.062 | 0.063 |
| $P_{PUE}$ | 0.073 | 0.044 | -0.011 | 0.161 |
| $P_{QUA}$ | 0.020 | 0.034 | -0.046 | 0.089 |
| $P_{SAC}$ | -0.034 | 0.042 | -0.115 | 0.047 |
| $P_{SAL}$ | -0.171 | 0.036 | -0.244 | -0.100 |
| $P_{SEG}$ | 0.007 | 0.031 | -0.052 | 0.070 |
| $P_{SIE}$ | 0.002 | 0.042 | -0.078 | 0.086 |
| $P_{STJ}$ | 0.172 | 0.029 | 0.117 | 0.231 |
| $P_{TAM}$ | -0.226 | 0.036 | -0.297 | -0.153 |
| $P_{VAL}$ | -0.047 | 0.038 | -0.120 | 0.027 |
| $P_{VER}$ | 0.106 | 0.030 | 0.047 | 0.165 |

**Table S17:** Parameter estimates of the provenance intercepts $P_p$ in *M1*. SD corresponds to the standard deviation and InfCI and SupCI correspond to the lower and upper bounds of the 0.95 credible interval.

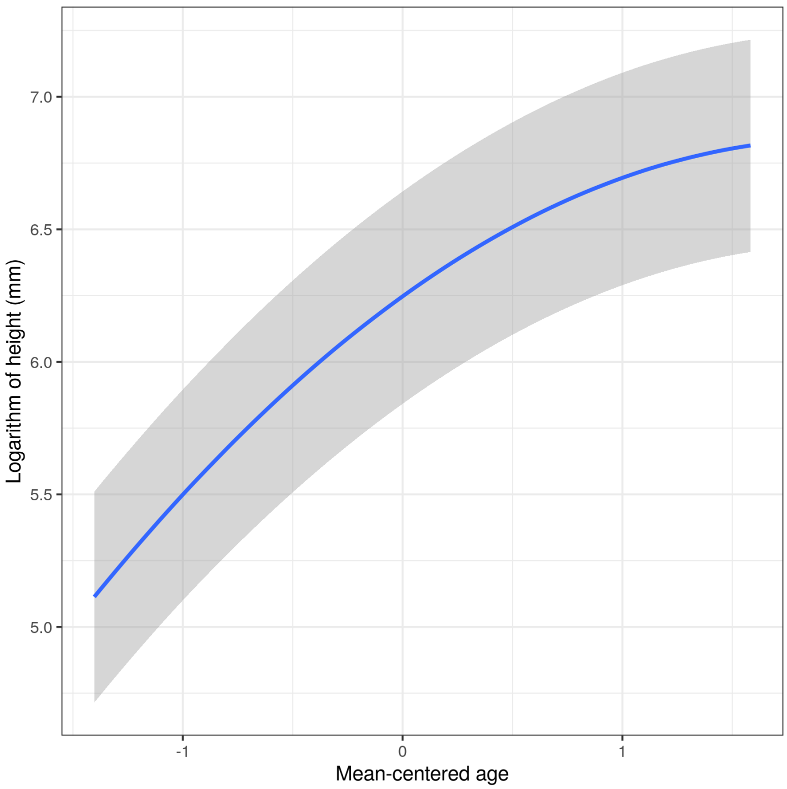

**Figure S11.** Relationship between mean-centered age and height on the log scale in *M1*.

##### *MODEL M2*

In *M2*, parameter estimates were similar to *M1*. The variance $\sigma^2_{Inter}$ of the provenance-by-site interaction was much smaller than the variances among provenances or genotypes (median of 0.004 and 95% CIs: 0.003-0.006; Table S18).

| Parameter | Median | SD | InfCI | SupCI |
|---|---|---|---|---|
| $\sigma^2_P$ | 0.011 | 0.004 | 0.007 | 0.022 |
| $\sigma^2_G$ | 0.012 | 0.001 | 0.010 | 0.014 |
| $\sigma^2_{Inter}$ | 0.004 | 0.001 | 0.003 | 0.006 |
| $\sigma^2_S$ | 0.113 | 1.525 | 0.030 | 1.423 |
| $\sigma^2_B$ | 0.002 | 0.001 | 0.001 | 0.003 |
| $\sigma^2$ | 0.096 | 0.001 | 0.095 | 0.098 |
| $\beta_0$ | 6.231 | 0.285 | 5.753 | 6.670 |
| $\beta_{age}$ | 0.597 | 0.003 | 0.592 | 0.602 |
| $\beta_{age2}$ | -0.150 | 0.002 | -0.155 | -0.145 |

**Table S18:** Parameter estimates of the varying-intercept variances ($\sigma^2_S$, $\sigma^2_B$, $\sigma^2_P$, $\sigma^2_G$ and $\sigma^2_{Inter}$), the global variance $\sigma$, the global intercept $\beta_0$ and the slopes associated with the age effect ($\beta_{age}$ and $\beta_{age2}$). SD corresponds to the standard deviation and InfCI and SupCI correspond to the lower and upper bounds of the 0.95 credible interval.

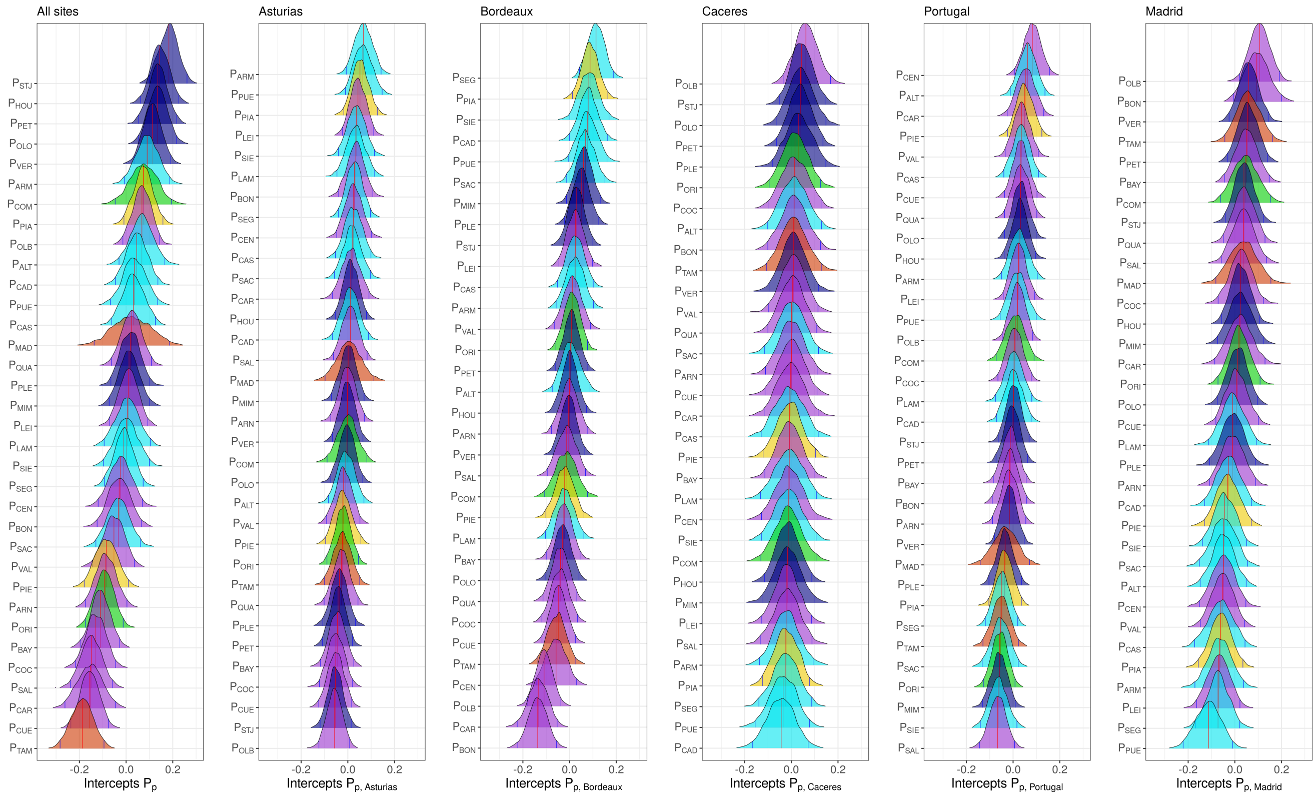

**Figure S12.** Posterior distributions of the provenance intercepts across all sites ($P_p$) and the site-specific intercepts of each provenance ($P_{p,s}$) from *model M2*. The colors correspond to the gene pool from which each population mainly belongs: gene pool in Northern Africa in orange, gene pool in Corsica in yellow, gene pool in Central Spain in purple, gene pool in the French Atlantic region in marine blue, gene pool in the Iberian Atlantic region in sky blue and gene pool in South-Eastern Spain in green.

Provenance varying intercepts are shown in fig. 3 (global intercepts across all sites from *M1*) and fig. S12 (site-specific intercepts from *M2*). Most provenances from the French Atlantic gene pool (e.g. STJ, HOU, PET, OLO and VER) were taller on average than other provenances in all sites. Most provenances from the Iberian Atlantic gene pool (e.g. ARM or PUE) were among the tallest in Asturias and Bordeaux but the shortest in Madrid and Cáceres. In contrast, most provenances from the Central Spain gene pool were among the shortest, especially in Asturias and Bordeaux. The two provenances from the Corsican gene pool showed highly contrasted intercepts. The provenance Pinia (PIA) was taller than other provenances on average, especially in Asturias and Bordeaux. In contrast, the second Corsican provenance (Pineta, PIE), was shorter than other provenances on average but grew particularly well in Portugal. Provenances belonging to the south-eastern Spain gene pool (i.e. ORI and COM) also showed contrasted intercepts (trees from ORI were shorter than those from COM on average), but did not show strong differences between sites. Lastly, regarding provenances belonging to the northern African gene pool, the TAM provenance had the lowest growth in our dataset and was more likely to be taller in Cáceres and Madrid (the harsh Mediterranean sites) than in the other sites.

##### 6.1.2 *Models M3 and M3bis*: potential drivers underlying the plastic component of height-growth variation

The plastic component of height-growth in *M3* (but also in all subsequent models until *M6*; Table 1) was only marginally associated with the climatic similarity among sites ($\sigma^2_{cs_{is}}$ with a median of 0.023 in *M3*), compared to the variance associated with site intercepts ($\sigma^2_S$ with a median of 0.126 in *M3*; Table S19). However, estimates of the intercepts ($S_s$ and $cs_{is}$) and variances ($\sigma^2_S$ and $\sigma^2_{cs_{is}}$) were uncertain (Tables S19 to S21), and the median and the credible interval of $\sigma^2_S$ increased from *M1* to *M3*, suggesting that *M3* may hardly separate between $\sigma^2_S$ and $\sigma^2_{cs_{is}}$ (Tables S15 & S19). To check this, we ran a supplementary model identical to *M3* but without the site intercepts $S_s$ (see *model M3bis* below). In this model, the variance related to the climatic similarity among sites was nearly equal and as uncertain as in *M3*, suggesting that our variance estimation of the plastic component in *M3* was robust (Table S23). However, the posterior distributions of the intercepts in *M3bis* were different from *M3*: height growth was positively associated with the climatic conditions in Bordeaux and Asturias, and negatively with the climatic conditions in Madrid and Cáceres, the two Mediterranean sites, and to a lesser extent in Portugal (Table S24).

***MODEL M3***

| Parameter | Median | SD | InfCI | SupCI |
|---|---|---|---|---|
| $\sigma^2_G$ | 0.012 | 0.001 | 0.010 | 0.013 |
| $\sigma^2_P$ | 0.013 | 0.004 | 0.008 | 0.023 |
| $\sigma^2_S$ | 0.126 | 1.694 | 0.027 | 1.895 |
| $\sigma^2_{cs_{is}}$ | 0.023 | 0.314 | 0.003 | 0.274 |
| $\sigma^2_B$ | 0.002 | 0.001 | 0.001 | 0.003 |
| $\sigma^2$ | 0.097 | 0.001 | 0.096 | 0.099 |
| $\beta_0$ | 6.295 | 0.315 | 5.825 | 6.808 |
| $\beta_{age}$ | 0.604 | 0.013 | 0.577 | 0.625 |
| $\beta_{age2}$ | -0.186 | 0.016 | -0.223 | -0.157 |

**Table S19:** Parameter estimates of the varying-intercept variances ($\sigma^2_S$, $\sigma^2_B$, $\sigma^2_P$, $\sigma^2_G$ and $\sigma^2_{cs_{is}}$), the global variance $\sigma^2$, the global intercept $\beta_0$ and the slopes associated with the age effect ($\beta_{age}$ and $\beta_{age2}$). SD corresponds to the standard deviation and InfCI and SupCI correspond to the lower and upper bounds of the 0.95 credible interval.

| Parameter | Median | SD | InfCI | SupCI |
|---|---|---|---|---|
| $S_{Asturias}$ | 0.283 | 0.326 | -0.248 | 0.809 |
| $S_{Bordeaux}$ | 0.164 | 0.341 | -0.382 | 0.726 |
| $S_{Caceres}$ | 0.086 | 0.341 | -0.520 | 0.596 |
| $S_{Madrid}$ | -0.349 | 0.321 | -0.902 | 0.092 |
| $S_{Portugal}$ | -0.162 | 0.320 | -0.717 | 0.291 |

**Table S20:** Parameter estimates of the site intercepts $S_s$ in *model M3*. SD corresponds to the standard deviation and InfCI and SupCI correspond to the lower and upper bounds of the 0.95 credible interval.

| Parameter | Median | SD | InfCI | SupCI |
|---|---|---|---|---|
| $cs_{1,Asturias}$ | 0.012 | 0.076 | -0.139 | 0.168 |
| $cs_{2,Asturias}$ | -0.027 | 0.085 | -0.198 | 0.141 |
| $cs_{3,Asturias}$ | 0.010 | 0.040 | -0.047 | 0.114 |
| $cs_{1,Bordeaux}$ | -0.109 | 0.126 | -0.353 | 0.160 |
| $cs_{2,Bordeaux}$ | 0.043 | 0.084 | -0.096 | 0.246 |
| $cs_{1,Caceres}$ | -0.001 | 0.145 | -0.347 | 0.242 |
| $cs_{1,Madrid}$ | -0.035 | 0.064 | -0.166 | 0.095 |
| $cs_{1,Portugal}$ | 0.028 | 0.055 | -0.089 | 0.132 |
| $cs_{2,Portugal}$ | 0.065 | 0.051 | -0.040 | 0.165 |
| $cs_{3,Portugal}$ | 0.033 | 0.048 | -0.072 | 0.123 |
| $cs_{4,Portugal}$ | -0.019 | 0.061 | -0.142 | 0.103 |

**Table S21:** Parameter estimates of the $cs_{is}$ intercepts related to climatic similarity between test sites during the year preceding the measurements. As the saplings were measured 2 to 4 times in Asturias, Bordeaux and Portugal, the numbers 1 to 4 correspond to each measurement, in the temporal order in which they were done. For example, the intercept $cs_{1,Asturias}$ corresponds to the first measurement taken in Asturias, when the saplings were 10 month old. See the table S1 for the sapling age at each measurement. SD corresponds to the standard deviation and InfCI and SupCI correspond to the lower and upper bounds of the 0.95 credible interval.

#### MODEL *M3bis*

| Models | **Explanatory part**: training P1 | | | | **Predictive part**: test P1 | | | |
|---|---|---|---|---|---|---|---|---|
| | $\mathcal{R}^2_m\|age$ | $\mathcal{R}^2_m$ | $\mathcal{R}^2_m(fix)$ | **PE**$_m$ | prediction $\mathcal{R}^2_m\|age$ | prediction $\mathcal{R}^2_m$ | prediction $\mathcal{R}^2_m(fix)$ | **PE**$_m$ |
| **M3** | 0.574 [0.552-0.597] | 0.816 [0.807-0.826] | 0.566 [0.506-0.621] | 0.231 [0.009-0.747] | 0.563 [0.538-0.587] | 0.815 [0.805-0.825] | 0.575 [0.514-0.631] | 0.235 [0.009-0.763] |
| **M13** | 0.575 [0.553-0.597] | 0.816 [0.807-0.826] | 0.634 [0.594-0.673] | 0.23 [0.009-0.746] | 0.563 [0.54-0.587] | 0.816 [0.806-0.825] | 0.644 [0.604-0.685] | 0.235 [0.009-0.767] |

**Table S22:** Comparing the performance of *M3* and *M3bis* in the P1 partition, in which the training dataset was obtained by randomly sampling 75% of the observations and the test dataset contains the remaining 25% observations. $\mathcal{R}^2_m|age$ and *prediction* $\mathcal{R}^2_m|age$ correspond to the proportion of variance explained and predicted by the models conditional on the age effect in the training (in-sample) and test (out-of-sample) datasets, respectively. $\mathcal{R}^2_m$ and $\mathcal{R}^2_m(fix)$ correspond to the proportion of explained variance in the training dataset by the entire model and by the fixed variables only, respectively. *Prediction* $\mathcal{R}^2_m$ and $\mathcal{R}^2_m(fix)$ correspond to the proportion of variance in the test dataset predicted by the entire model or by the fixed effects, respectively. PE$_m$ corresponds to the mean predictive error (mean of observed minus predicted responses). For $\mathcal{R}^2$ and PE, the mean (over all iterations for $\mathcal{R}^2$ or over all observations for PE) and the 95% credible intervals are given. For more details on the calculation of each index, see section 5 of the Supplementary Information.

| Parameter | Median | SD | InfCI | SupCI |
|---|---|---|---|---|
| $\sigma^2_B$ | 0.029 | 0.009 | 0.017 | 0.053 |
| $\sigma^2_P$ | 0.013 | 0.004 | 0.008 | 0.022 |
| $\sigma^2_G$ | 0.012 | 0.001 | 0.010 | 0.013 |
| $\sigma^2_{cs_{is}}$ | 0.025 | 0.083 | 0.006 | 0.229 |
| $\sigma^2$ | 0.097 | 0.001 | 0.096 | 0.099 |
| $\beta_0$ | 6.284 | 0.036 | 6.213 | 6.356 |
| $\beta_{age}$ | 0.630 | 0.008 | 0.614 | 0.645 |
| $\beta_{age2}$ | -0.153 | 0.010 | -0.172 | -0.134 |

**Table S23:** Parameter estimates of the varying-intercept variances ($\sigma^2_B$, $\sigma^2_P$, $\sigma^2_G$ and $\sigma^2_{cs_{is}}$), the global variance $\sigma^2$, the global intercept $\beta_0$ and the slopes associated with the age effect ($\beta_{age}$ and $\beta_{age2}$). SD corresponds to the standard deviation and InfCI and SupCI correspond to the lower and upper bounds of the 0.95 credible interval.

| Parameter | Median | SD | InfCI | SupCI |
|---|---|---|---|---|
| $cs_{1,Asturias}$ | 0.133 | 0.026 | 0.080 | 0.180 |
| $cs_{2,Asturias}$ | 0.117 | 0.030 | 0.054 | 0.172 |
| $cs_{3,Asturias}$ | 0.026 | 0.020 | -0.009 | 0.069 |
| $cs_{1,Bordeaux}$ | 0.085 | 0.042 | -0.003 | 0.162 |
| $cs_{2,Bordeaux}$ | 0.134 | 0.031 | 0.076 | 0.195 |
| $cs_{1,Caceres}$ | -0.158 | 0.053 | -0.264 | -0.057 |
| $cs_{1,Madrid}$ | -0.140 | 0.030 | -0.198 | -0.082 |
| $cs_{1,Portugal}$ | -0.047 | 0.022 | -0.088 | -0.004 |
| $cs_{2,Portugal}$ | -0.007 | 0.023 | -0.051 | 0.041 |
| $cs_{3,Portugal}$ | -0.028 | 0.020 | -0.066 | 0.011 |
| $cs_{4,Portugal}$ | -0.114 | 0.028 | -0.168 | -0.056 |

**Table S24:** Parameter estimates of the $cs_{is}$ intercepts related to climatic similarity between test sites during the year preceding the measurements. SD corresponds to the standard deviation and InfCI and SupCI correspond to the lower and upper bounds of the 0.95 credible interval.

##### 6.1.3 *Models M4 to M6*: potential drivers underlying the genetic component of height-growth variation

In *M4*, the variance $\sigma^2_{g_j}$ between gene pools was as important as the variance $\sigma^2_G$ between genotypes, but had a very wide posterior distribution (Table S25). From *M3* to *M4*, adding the gene pool intercepts $g_j$ resulted in a decreased variance between provenances (from a median of 0.013 to a median of 0.006, Tables S19 & S25). This indicates redundant information between gene pools and provenances. Genotypes belonging to the French Atlantic gene pool, and to a lesser extent to the Iberian Atlantic gene pool, were on average taller than genotypes belonging to the northern Africa gene pool, and to a lesser extent to the Central Spain gene pool (fig. S14).

***MODEL M4***

| Parameter | Median | SD | InfCI | SupCI |
|---|---|---|---|---|
| $\sigma^2_{g_j}$ | 0.013 | 0.044 | 0.002 | 0.093 |
| $\sigma^2_P$ | 0.006 | 0.003 | 0.003 | 0.013 |
| $\sigma^2_G$ | 0.012 | 0.001 | 0.010 | 0.013 |
| $\sigma^2_S$ | 0.127 | 1.365 | 0.026 | 2.125 |
| $\sigma^2_{cs_{is}}$ | 0.022 | 0.187 | 0.003 | 0.267 |
| $\sigma^2_B$ | 0.002 | 0.001 | 0.001 | 0.003 |
| $\sigma^2$ | 0.097 | 0.001 | 0.095 | 0.099 |
| $\beta_0$ | 6.280 | 0.269 | 5.704 | 6.811 |
| $\beta_{age}$ | 0.605 | 0.012 | 0.577 | 0.626 |
| $\beta_{age2}$ | -0.185 | 0.016 | -0.221 | -0.156 |

**Table S25:** Parameter estimates of the varying-intercept variances ($\sigma^2_S$, $\sigma^2_B$, $\sigma^2_P$, $\sigma^2_G$, $\sigma^2_{cs_{is}}$ and $\sigma^2_{g_j}$), the global variance $\sigma^2$, the global intercept $\beta_0$ and the slopes associated with the age effect ($\beta_{age}$ and $\beta_{age2}$). SD corresponds to the standard deviation and InfCI and SupCI correspond to the lower and upper bounds of the 0.95 credible interval.

***MODEL M5*: gene pool-specific heritabilities, the telltale of distinct adaptive histories**

Heritability variation across populations or gene pools can inform on differences in the drivers underlying their adaptive histories, such as evolutionary mechanisms (e.g. capacity of dispersion, selection strength, migration) and local environmental constraints (e.g. environmental heterogeneity). Using CLONAPIN data from all sites except Bordeaux, Rodríguez-Quilón et al. (2016) showed that populations from the Mediterranean gene pools had a higher heritability than those from the Atlantic gene pools. Nonetheless, their heritability estimates did not consider population admixture, a notable feature in maritime pine (fig. 1). In our study, we applied the recent methodology proposed by Muff et al. (2019) to calculate gene pool-specific total genetic variance and broad-sense heritabilities in a single model that accounts for population admixture. We showed that the total genetic variance of the Iberian Atlantic gene pool had a probability higher than 0.95 of being lower than that of the Corsican and south-eastern Spain gene pools, and a probability higher than 0.90 of being lower than that of the Central Spain gene pool (Table S27; see also fig. S13), despite their geographical proximity. The total genetic variance of the French Atlantic gene pool had a probability higher than 0.90 of being lower than that of the Corsican and south-eastern Spain gene pools (Table S27; see also fig. S13). However, this should be taken with caution as the total variance of the gene pools from south-eastern Spain, Corsica and northern Africa had wide posterior distributions, which is probably due to the small number of genotypes from these gene pools (see Table S3). In line with the genetic variance estimates, the medians of the gene-pool specific estimates of heritability $H^2_j$ varied between 0.104 in the Iberian Atlantic gene-pool (95% CIs: 0.065-0.146) and 0.223 in the south-eastern Spain gene pool (95% CIs: 0.093-0.363) (Table S28 and fig. S13A). Interestingly, provenances that showed the highest broad-sense heritabilities (i.e. provenances from the Corsican, south-eastern Spain, and central Spain gene pools) are also the ones facing more contrasted climates. Indeed, Corsica and south-eastern Spain are mountainous areas, with strong environmental heterogeneity at small spatial scales. Central Spain is less contrasted spatially but experiences a high continentality, and thus strong daily and annually climatic variation. Noticeably, genotypes displayed genetic values mostly determined by the dominant gene pool from which they belong to, but also, to a lesser extent, by other gene pools (fig. S13B). Taken together, these results may suggest that gene pools from regions with high environmental heterogeneity (in space and/or time) have also higher heritability. Indeed, theoretical and empirical works have proposed that high levels of adaptive genetic variance can be maintained in regions of high environmental heterogeneity (mainly spatial, but to a lesser extent also temporal) and some degree of gene flow between populations (Yeaman and Jarvis 2006, Yeaman and Otto 2011, McDonald and Yeaman 2018). Further research involving multiple adaptive traits and more detailed environmental data would be needed to confirm this hypothesis.

| Parameter | Median | SD | InfCI | SupCI |
|---|---|---|---|---|
| $\sigma^2_{A_{NA}}$ | 0.015 | 0.010 | 0.005 | 0.041 |
| $\sigma^2_{A_C}$ | 0.025 | 0.010 | 0.013 | 0.050 |
| $\sigma^2_{A_{CS}}$ | 0.017 | 0.003 | 0.013 | 0.024 |
| $\sigma^2_{A_{FA}}$ | 0.015 | 0.002 | 0.012 | 0.020 |
| $\sigma^2_{A_{IA}}$ | 0.011 | 0.003 | 0.007 | 0.017 |
| $\sigma^2_{A_{SES}}$ | 0.027 | 0.012 | 0.011 | 0.059 |
| $\sigma^2_{g_j}$ | 0.004 | 0.031 | 0.000 | 0.064 |
| $\sigma^2_P$ | 0.011 | 0.004 | 0.006 | 0.021 |
| $\sigma^2_S$ | 0.121 | 0.565 | 0.026 | 1.259 |
| $\sigma^2_{cs_{is}}$ | 0.024 | 0.157 | 0.003 | 0.262 |
| $\sigma^2_B$ | 0.002 | 0.001 | 0.001 | 0.003 |
| $\sigma^2$ | 0.098 | 0.001 | 0.096 | 0.099 |
| $\beta_0$ | 6.289 | 0.220 | 5.819 | 6.708 |
| $\beta_{age}$ | 0.605 | 0.012 | 0.578 | 0.626 |
| $\beta_{age2}$ | -0.185 | 0.016 | -0.221 | -0.158 |

**Table S26:** Parameter estimates of the varying-intercept variances ($\sigma^2_S$, $\sigma^2_B$, $\sigma^2_P$, $\sigma^2_G$, $\sigma^2_{cs_{is}}$ and $\sigma_{A_j}$), the gene-pool specific total genetic variances ($\sigma_{A_j}$), the global variance $\sigma^2$, the global intercept $\beta_0$ and the slopes associated with the age effect ($\beta_{age}$ and $\beta_{age2}$). SD corresponds to the standard deviation and InfCI and SupCI correspond to the lower and upper bounds of the 0.95 credible interval.

| Hypothesis | Estimate | Est.Error | CI.Lower | CI.Upper | Evid.Ratio | Post.Prob | Star |
|---|---|---|---|---|---|---|---|
| $\sigma^2_{A_{NA}} - \sigma^2_{A_C} < 0$ | -0.035 | 0.043 | -0.104 | 0.035 | 4.291 | 0.811 | |
| $\sigma^2_{A_{NA}} - \sigma^2_{A_{CS}} < 0$ | -0.007 | 0.035 | -0.057 | 0.056 | 1.656 | 0.623 | |
| $\sigma^2_{A_{NA}} - \sigma^2_{A_{FA}} < 0$ | 0.001 | 0.035 | -0.049 | 0.065 | 1.127 | 0.530 | |
| $\sigma^2_{A_{NA}} - \sigma^2_{A_{IA}} < 0$ | 0.019 | 0.036 | -0.033 | 0.084 | 0.450 | 0.310 | |
| $\sigma^2_{A_{NA}} - \sigma^2_{A_{SES}} < 0$ | -0.042 | 0.052 | -0.121 | 0.046 | 4.396 | 0.815 | |
| $\sigma^2_{A_C} - \sigma^2_{A_{NA}} < 0$ | 0.035 | 0.043 | -0.035 | 0.104 | 0.233 | 0.189 | |
| $\sigma^2_{A_C} - \sigma^2_{A_{CS}} < 0$ | 0.029 | 0.030 | -0.015 | 0.081 | 0.183 | 0.155 | |
| $\sigma^2_{A_C} - \sigma^2_{A_{FA}} < 0$ | 0.037 | 0.029 | -0.006 | 0.089 | 0.091 | 0.084 | |
| $\sigma^2_{A_C} - \sigma^2_{A_{IA}} < 0$ | 0.055 | 0.030 | 0.009 | 0.109 | 0.022 | 0.022 | |
| $\sigma^2_{A_C} - \sigma^2_{A_{SES}} < 0$ | -0.006 | 0.044 | -0.080 | 0.065 | 1.239 | 0.553 | |
| $\sigma^2_{A_{CS}} - \sigma^2_{A_{NA}} < 0$ | 0.007 | 0.035 | -0.056 | 0.057 | 0.604 | 0.377 | |
| $\sigma^2_{A_{CS}} - \sigma^2_{A_C} < 0$ | -0.029 | 0.030 | -0.081 | 0.015 | 5.459 | 0.845 | |
| $\sigma^2_{A_{CS}} - \sigma^2_{A_{FA}} < 0$ | 0.008 | 0.013 | -0.013 | 0.030 | 0.384 | 0.277 | |
| $\sigma^2_{A_{CS}} - \sigma^2_{A_{IA}} < 0$ | 0.026 | 0.017 | -0.001 | 0.053 | 0.064 | 0.060 | |
| $\sigma^2_{A_{CS}} - \sigma^2_{A_{SES}} < 0$ | -0.035 | 0.038 | -0.103 | 0.022 | 4.859 | 0.829 | |
| $\sigma^2_{A_{FA}} - \sigma^2_{A_{NA}} < 0$ | -0.001 | 0.035 | -0.065 | 0.049 | 0.887 | 0.470 | |
| $\sigma^2_{A_{FA}} - \sigma^2_{A_C} < 0$ | -0.037 | 0.029 | -0.089 | 0.006 | 10.952 | 0.916 | * |
| $\sigma^2_{A_{FA}} - \sigma^2_{A_{CS}} < 0$ | -0.008 | 0.013 | -0.030 | 0.013 | 2.606 | 0.723 | |
| $\sigma^2_{A_{FA}} - \sigma^2_{A_{IA}} < 0$ | 0.018 | 0.014 | -0.006 | 0.041 | 0.117 | 0.105 | |
| $\sigma^2_{A_{FA}} - \sigma^2_{A_{SES}} < 0$ | -0.043 | 0.035 | -0.107 | 0.010 | 9.435 | 0.904 | * |
| $\sigma^2_{A_{IA}} - \sigma^2_{A_{NA}} < 0$ | -0.019 | 0.036 | -0.084 | 0.033 | 2.221 | 0.690 | |
| $\sigma^2_{A_{IA}} - \sigma^2_{A_C} < 0$ | -0.055 | 0.030 | -0.109 | -0.009 | 44.455 | 0.978 | ** |
| $\sigma^2_{A_{IA}} - \sigma^2_{A_{CS}} < 0$ | -0.026 | 0.017 | -0.053 | 0.001 | 15.667 | 0.940 | * |
| $\sigma^2_{A_{IA}} - \sigma^2_{A_{FA}} < 0$ | -0.018 | 0.014 | -0.041 | 0.006 | 8.524 | 0.895 | |
| $\sigma^2_{A_{IA}} - \sigma^2_{A_{SES}} < 0$ | -0.061 | 0.036 | -0.125 | -0.006 | 29.928 | 0.968 | ** |
| $\sigma^2_{A_{SES}} - \sigma^2_{A_{NA}} < 0$ | 0.042 | 0.052 | -0.046 | 0.121 | 0.227 | 0.185 | |
| $\sigma^2_{A_{SES}} - \sigma^2_{A_C} < 0$ | 0.006 | 0.044 | -0.065 | 0.080 | 0.807 | 0.447 | |
| $\sigma^2_{A_{SES}} - \sigma^2_{A_{CS}} < 0$ | 0.035 | 0.038 | -0.022 | 0.103 | 0.206 | 0.171 | |
| $\sigma^2_{A_{SES}} - \sigma^2_{A_{FA}} < 0$ | 0.043 | 0.035 | -0.010 | 0.107 | 0.106 | 0.096 | |
| $\sigma^2_{A_{SES}} - \sigma^2_{A_{IA}} < 0$ | 0.061 | 0.036 | 0.006 | 0.125 | 0.033 | 0.032 | |

**Table S27:** One-sided hypothesis testing on the probability that the gene pool-specific total genetic variances in *M5* are different, using the function ‘hypothesis‘ from the ‘brms‘ package (Bürkner 2017). ‘Est.Error‘ is the standard deviation of the estimated difference between two genetic variances (‘Estimate‘). The ‘CI.Lower‘ and ‘CI.Upper‘ are the lower and upper bounds of the 95% credible interval, respectively. ‘Evid.Ratio‘ is the evidence ratio of each hypothesis, i.e. the posterior probability (‘Post.Prob‘) under the hypothesis against its alternative. For instance, the evidence ratio of the hypothesis $\sigma^2_{A_{NA}} - \sigma^2_{A_C} < 0$ is the ratio of the posterior probability of $\sigma^2_{A_{NA}} - \sigma^2_{A_C} < 0$ and the posterior probability of $\sigma^2_{A_{NA}} - \sigma^2_{A_C} > 0$. The * and ** in the ’Star’ column indicate hypotheses with a posterior probability higher than 0.90 and 0.95, respectively.

Using the estimates of the gene-pool specific total genetic variances from *M5*, the gene-pool specific heritabilities were calculated such as:

$$H_j^2 = \frac{\sigma_{A_j}^2}{\sigma_{A_j}^2 + \sigma^2}$$

where $\sigma^2$ is the modeled residual variance and $\sigma_{A_j}^2$ the gene-pool specific total genetic variances.

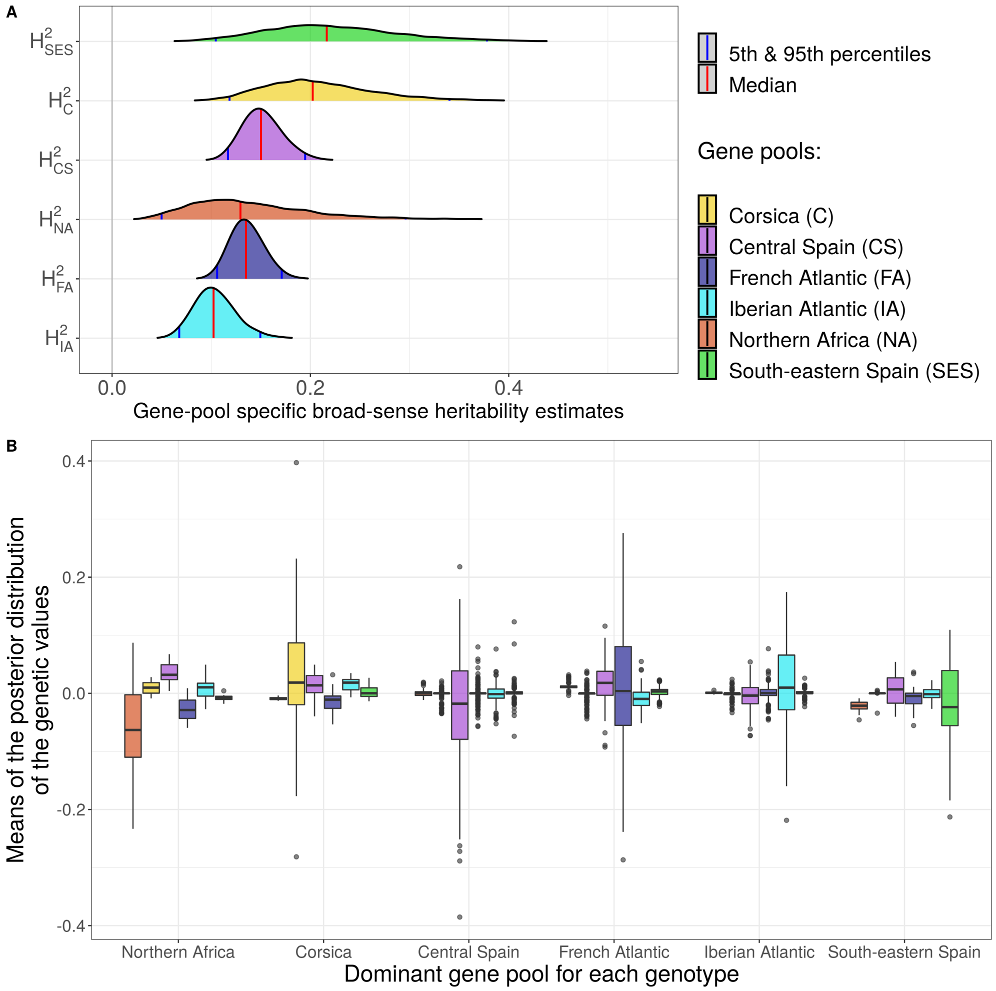

**Figure S13.** Posterior distribution of $H_j^2$, the gene pool-specific heritabilities obtained from *model M5* (A). Boxplot of he gene pool partial genetic values for genotypes grouped according to the main gene pool to which they belong (B). The colors represent the gene pool from which the partial genetic values were estimated. The lower, middle and upper parts of the hinges correspond to the 25%, 50% and 75% quantiles, respectively. The lower/upper whiskers extend from the hinge to the smallest/largest value no further than 1.5 time the IQR from the hinge (where IQR is the distance between the first and third quartiles).

| Parameter | Median | SD | InfCI | SupCI |
|---|---|---|---|---|
| $H^2_{NA}$ | 0.141 | 0.064 | 0.034 | 0.266 |
| $H^2_{C}$ | 0.210 | 0.057 | 0.107 | 0.322 |
| $H^2_{CS}$ | 0.152 | 0.020 | 0.114 | 0.190 |
| $H^2_{FA}$ | 0.136 | 0.017 | 0.105 | 0.169 |
| $H^2_{IA}$ | 0.104 | 0.021 | 0.065 | 0.146 |
| $H^2_{SES}$ | 0.223 | 0.070 | 0.093 | 0.363 |

**Table S28:** Gene pool-specific heritability estimates $H^2_j$. The gene pools come from: Northern Africa (NA), Corsica (C), Central Spain (CS), French Atlantic region (FA), Iberian Atlantic region (IA) and south-eastern Spain (SES). SD corresponds to the standard deviation and InfCI and SupCI correspond to the lower and upper bounds of the 0.95 credible interval.

***MODEL M6***

In *M6*, the variance $\sigma^2_{cp_p}$ associated with the provenance climate-of-origin had a higher median than the variances attributed to either gene pools or provenances ($\sigma^2_{g_j}$ and $\sigma^2_P$; Table S29). Including the provenance climate-of-origin in *M6* resulted in a decreased variance of the provenances and gene pools, suggesting confounded effects (Tables S25 & S29). Once these confounded effects are taking into account, trees from climatic regions neighboring the Atlantic Ocean were generally the tallest (e.g. CAD, SIE, PUE, LAM and CAS in northwestern Spain; all provenances along the French Atlantic coast; fig. S14). Interestingly, the Leiria (LEI) provenance, which has a strong Iberian Atlantic component (Table S3) and had the highest climate intercept estimate (similar to that of the French Atlantic provenances), was not among the tallest provenances (fig. S14). Also, the Corsican provenances showed contrasted climate intercepts, with a positive influence on height growth for Pinia (PIA) but not for Pineta (PIE), which could explain their striking differences in height-growth patterns (fig. 3). Finally, the four provenances from south-eastern Spain and northern Africa gene pools showed all negative climate intercepts (fig. S14).

| Parameter | Median | SD | InfCI | SupCI |
|---|---|---|---|---|
| $\sigma^2_{g_j}$ | 0.003 | 0.018 | 0.000 | 0.050 |
| $\sigma^2_P$ | 0.005 | 0.002 | 0.003 | 0.011 |
| $\sigma^2_{cp_p}$ | 0.009 | 0.291 | 0.000 | 0.244 |
| $\sigma^2_G$ | 0.012 | 0.001 | 0.010 | 0.014 |
| $\sigma^2_S$ | 0.128 | 0.761 | 0.026 | 1.436 |
| $\sigma^2_{cs_{is}}$ | 0.023 | 0.111 | 0.003 | 0.297 |
| $\sigma^2_B$ | 0.002 | 0.001 | 0.001 | 0.003 |
| $\sigma^2$ | 0.097 | 0.001 | 0.096 | 0.099 |
| $\beta_0$ | 6.280 | 0.235 | 5.777 | 6.723 |
| $\beta_{age}$ | 0.605 | 0.013 | 0.576 | 0.627 |
| $\beta_{age2}$ | -0.185 | 0.018 | -0.224 | -0.156 |

**Table S29:** Parameter estimates of the varying-intercept variances ($\sigma^2_S$, $\sigma^2_B$, $\sigma^2_P$, $\sigma^2_G$, $\sigma^2_{cs_{is}}$ and $\sigma^2_{cp_p}$), the gene-pool specific total genetic variances ($\sigma_{A_j}$), the global variance $\sigma^2$, the global intercept $\beta_0$ and the slopes associated with the age effect ($\beta_{age}$ and $\beta_{age2}$). SD corresponds to the standard deviation and InfCI and SupCI correspond to the lower and upper bounds of the 0.95 credible interval.

#### Summary figure of models *models M3, M5* and *M6*

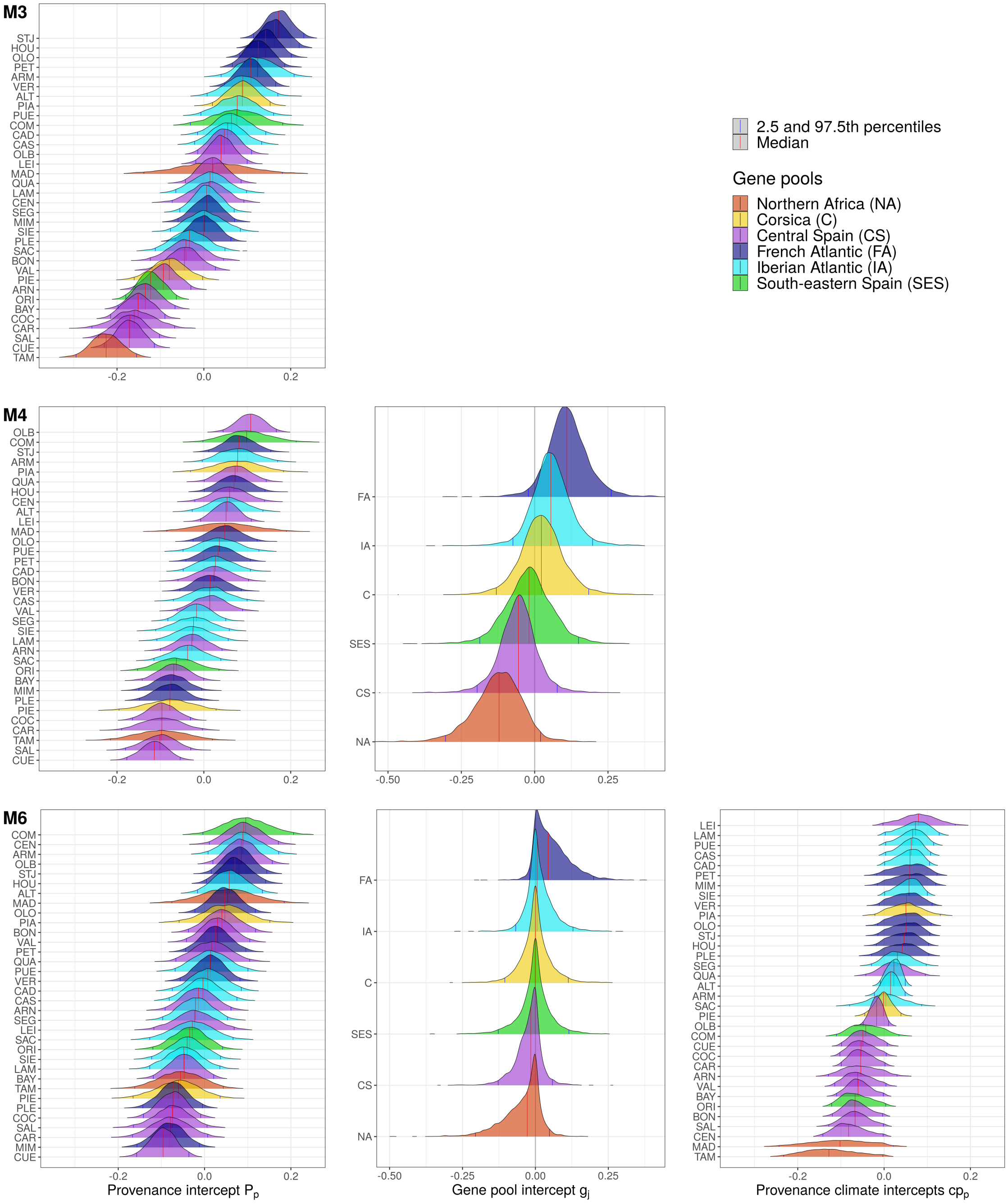

**Figure S14.** Posterior distributions of the provenance intercepts $P_p$, gene pool intercepts $g_j$ and the $cp_p$ intercepts related to the provenance climate. The first row corresponds to *M3* including only $P_p$. The second row corresponds to *M4* including both $P_p$ and $g_j$. The third row corresponds to *M6* including $P_p$, $g_j$ and $cp_p$. In *M3*, *M4* and *M6*, the plastic component is included in the same way via the intercepts $S_S$ and $cs_{is}$. Only the genetic component changes between these models. Provenances were colored based on the main gene pool they belong to.

##### 6.1.4 Predictive *models M7* and *M8*: combining climatic and genomic drivers

***MODEL M7***

| Term | Median | SD | InfCI | SupCI |
|---|---|---|---|---|
| $\sigma^2_B$ | 0.002 | 0.001 | 0.001 | 0.003 |
| $\sigma^2_{g_j}$ | 0.027 | 0.100 | 0.008 | 0.192 |
| $\sigma^2_S$ | 0.172 | 0.992 | 0.036 | 2.252 |
| $\sigma^2_{\beta_{max.temp,s}}$ | 0.001 | 0.005 | 0.000 | 0.009 |
| $\sigma^2_{\beta_{min.pre,s}}$ | 0.001 | 0.003 | 0.000 | 0.007 |
| $\sigma^2_{\beta_{gPEA,s}}$ | 0.018 | 0.048 | 0.005 | 0.135 |
| $\sigma^2$ | 0.104 | 0.001 | 0.102 | 0.106 |
| $\beta_0$ | 6.303 | 0.403 | 5.518 | 7.147 |
| $\beta_{age}$ | 0.599 | 0.003 | 0.593 | 0.604 |
| $\beta_{age2}$ | -0.151 | 0.003 | -0.156 | -0.146 |

**Table S30:** Parameter estimates from model *M7* fitted on the P1 partition. SD corresponds to the standard deviation and InfCI and SupCI correspond to the lower and upper bounds of the 0.95 credible interval.

***MODEL M8***

| Term | Median | SD | InfCI | SupCI |
|---|---|---|---|---|
| $\sigma^2_B$ | 0.002 | 0.001 | 0.001 | 0.003 |
| $\sigma^2_{g_j}$ | 0.033 | 0.078 | 0.009 | 0.228 |
| $\sigma^2_S$ | 0.185 | 1.548 | 0.041 | 2.349 |
| $\sigma^2_{\beta_{max.temp,s}}$ | 0.003 | 0.009 | 0.001 | 0.020 |
| $\sigma^2_{\beta_{min.pre,s}}$ | 0.002 | 0.005 | 0.000 | 0.014 |
| $\sigma^2_{\beta_{rPEA,s}}$ | 0.032 | 0.133 | 0.010 | 0.202 |
| $\sigma^2$ | 0.102 | 0.001 | 0.100 | 0.104 |
| $\beta_0$ | 6.333 | 0.353 | 5.631 | 7.034 |
| $\beta_{age}$ | 0.599 | 0.003 | 0.593 | 0.604 |
| $\beta_{age2}$ | -0.151 | 0.003 | -0.156 | -0.146 |

**Table S31:** Parameter estimates from model *M8* fitted on the P1 partition. SD corresponds to the standard deviation and InfCI and SupCI correspond to the lower and upper bounds of the 0.95 credible interval.

#### Summary figure of *models M7* and *M8*

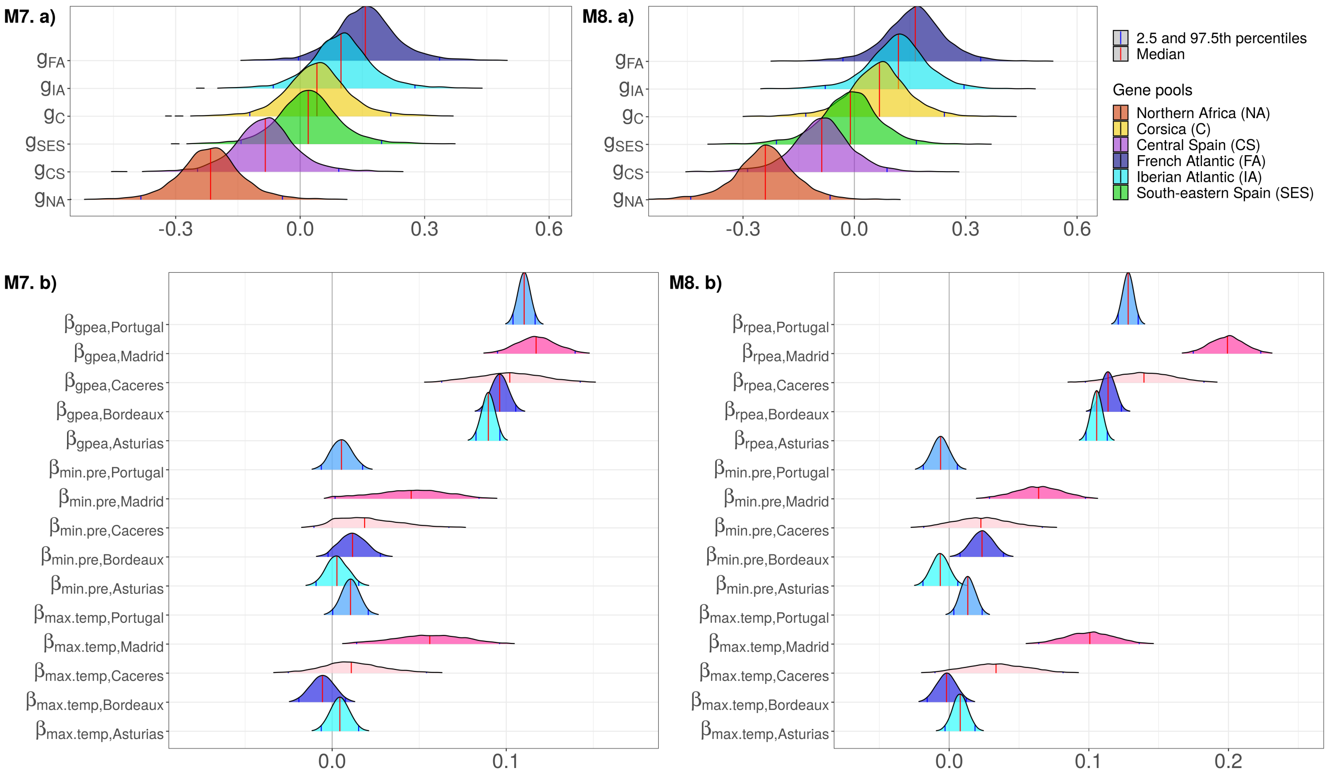

**Figure S15.** Posterior distributions of parameters from models *M7* and *M8* fitted on the P1 partition. In panels a), the parameter estimates $g_j$ correspond to the effect of the gene pools $j$. In panels b), the parameter estimates correspond to the site-specific effects of gPEAs in *M7* ($\beta_{gPEA,s}$), rPEAs in *M8* ($\beta_{rPEA,s}$), the minimum precipitation during the driest month ($\beta_{min.pre,s}$) and the maximum temperature of the warmest month ($\beta_{max.temp,s}$). Parameters specific to the same site are colored the same to aid visualization.

##### 6.1.5 Predictive *models M9* to *M12*: including separately climatic and genomic drivers

***MODEL M9***

| Parameter | Median | SD | InfCI | SupCI |
|---|---|---|---|---|
| $\sigma_{g_j}$ | 0.024 | 0.077 | 0.007 | 0.170 |
| $\sigma_S^2$ | 0.107 | 0.921 | 0.027 | 1.346 |
| $\sigma_B^2$ | 0.002 | 0.001 | 0.001 | 0.003 |
| $\sigma^2$ | 0.114 | 0.001 | 0.112 | 0.116 |
| $\beta_0$ | 6.233 | 0.237 | 5.708 | 6.637 |
| $\beta_{age}$ | 0.601 | 0.003 | 0.596 | 0.607 |
| $\beta_{age2}$ | -0.153 | 0.003 | -0.158 | -0.148 |

**Table S32:** Parameter estimates from model *M9* fitted on the P1 partition. SD corresponds to the standard deviation and InfCI and SupCI correspond to the lower and upper bounds of the 0.95 credible interval.

***MODEL M10***

| Term | Median | SD | InfCI | SupCI |
|---|---|---|---|---|
| $\sigma_B^2$ | 0.002 | 0.001 | 0.001 | 0.003 |
| $\sigma_S^2$ | 0.173 | 0.909 | 0.034 | 2.404 |
| $\sigma^2_{\beta_{max.temp,s}}$ | 0.005 | 0.032 | 0.001 | 0.033 |
| $\sigma^2_{\beta_{min.pre,s}}$ | 0.006 | 0.063 | 0.001 | 0.046 |
| $\sigma^2$ | 0.115 | 0.001 | 0.113 | 0.117 |
| $\beta_0$ | 6.202 | 0.399 | 5.237 | 6.869 |
| $\beta_{age}$ | 0.601 | 0.003 | 0.596 | 0.607 |
| $\beta_{age^2}$ | -0.153 | 0.003 | -0.159 | -0.148 |

**Table S33:** Parameter estimates from model *M10* fitted on the P1 partition. SD corresponds to the standard deviation and InfCI and SupCI correspond to the lower and upper bounds of the 0.95 credible interval.

***MODEL M11***

| Term | Median | SD | InfCI | SupCI |
|---|---|---|---|---|
| $\sigma_B^2$ | 0.002 | 0.001 | 0.001 | 0.003 |
| $\sigma_S^2$ | 0.249 | 4.243 | 0.040 | 7.779 |
| $\sigma^2_{\beta_{gPEA,s}}$ | 0.013 | 0.056 | 0.004 | 0.092 |
| $\sigma^2$ | 0.114 | 0.001 | 0.112 | 0.117 |
| $\beta_0$ | 6.322 | 0.763 | 4.768 | 7.742 |
| $\beta_{age}$ | 0.599 | 0.003 | 0.593 | 0.604 |
| $\beta_{age2}$ | -0.151 | 0.003 | -0.156 | -0.146 |

**Table S34:** Parameter estimates from model *M11* fitted on the P1 partition. SD corresponds to the standard deviation and InfCI and SupCI correspond to the lower and upper bounds of the 0.95 credible interval.

***MODEL M12***

| Term | Median | SD | InfCI | SupCI |
|---|---|---|---|---|
| $\sigma^2_B$ | 0.002 | 0.001 | 0.001 | 0.003 |
| $\sigma^2_S$ | 0.332 | 1.572 | 0.044 | 4.542 |
| $\sigma^2_{\beta_{rPEA,s}}$ | 0.023 | 0.047 | 0.007 | 0.152 |
| $\sigma^2$ | 0.113 | 0.001 | 0.111 | 0.115 |
| $\beta_0$ | 6.597 | 0.499 | 5.735 | 7.663 |
| $\beta_{age}$ | 0.599 | 0.003 | 0.593 | 0.604 |
| $\beta_{age2}$ | -0.151 | 0.003 | -0.156 | -0.146 |

**Table S35:** Parameter estimates from model *M12* fitted on the P1 partition. SD corresponds to the standard deviation and InfCI and SupCI correspond to the lower and upper bounds of the 0.95 credible interval.

##### Summary figure of *models M9* to *M12*

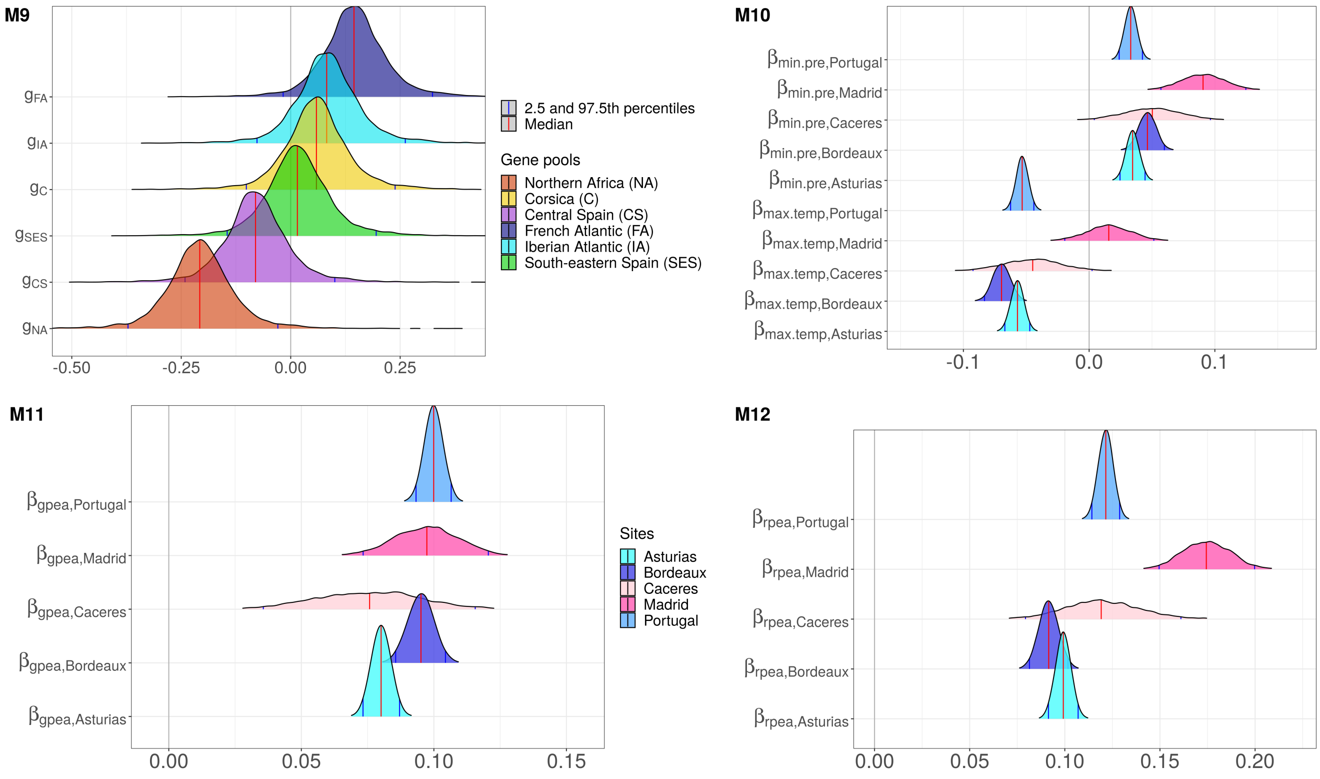

**Figure S16.** Posterior distributions of parameters from models *M9* and *M12* fitted on P1. For *M9*, the parameter estimates $g_j$ correspond to the effect of the gene pools $j$. For *M10*, the parameter estimates correspond to the site-specific effects of the minimum precipitation during the driest month ($\beta_{min.pre,s}$) and the maximum temperature of the warmest month ($\beta_{max.temp,s}$). For *M11*, the parameter estimates correspond to the site-specific effect of the gPEAs ($\beta_{gPEA,s}$). For *M12*, parameter estimates correspond to the site-specific effects of the rPEAs ($\beta_{rPEA,s}$).

#### 6.2 P2 partition (random split of the provenances)

The unknown provenances of the P2 test dataset were chosen randomly among the 34 provenances of our study.

##### 6.2.1 *M0*, *M1* and *M2*

| Parameter | Median | SD | InfCI | SupCI |
|---|---|---|---|---|
| $\sigma_S^2$ | 0.101 | 0.527 | 0.025 | 1.165 |
| $\sigma_B^2$ | 0.002 | 0.001 | 0.001 | 0.004 |
| $\sigma^2$ | 0.124 | 0.001 | 0.121 | 0.126 |
| $\beta_0$ | 6.278 | 0.210 | 5.840 | 6.631 |
| $\beta_{age}$ | 0.602 | 0.003 | 0.597 | 0.607 |
| $\beta_{age2}$ | -0.153 | 0.003 | -0.158 | -0.148 |

**Table S36:** Parameter estimates from model M0 fitted on the P2 partition. SD corresponds to the standard deviation and InfCI and SupCI correspond to the lower and upper bounds of the 0.95 credible interval.

| Parameter | Median | SD | InfCI | SupCI |
|---|---|---|---|---|
| $\sigma_P^2$ | 0.016 | 0.005 | 0.009 | 0.029 |
| $\sigma_G^2$ | 0.011 | 0.001 | 0.010 | 0.013 |
| $\sigma_S^2$ | 0.110 | 0.540 | 0.028 | 1.032 |
| $\sigma_B^2$ | 0.002 | 0.001 | 0.001 | 0.003 |
| $\sigma^2$ | 0.097 | 0.001 | 0.095 | 0.099 |
| $\beta_0$ | 6.240 | 0.190 | 5.866 | 6.627 |
| $\beta_{age}$ | 0.599 | 0.002 | 0.594 | 0.603 |
| $\beta_{age2}$ | -0.150 | 0.002 | -0.155 | -0.146 |

**Table S37:** Parameter estimates from model *M1* fitted on the P2 partition. SD corresponds to the standard deviation and InfCI and SupCI correspond to the lower and upper bounds of the 0.95 credible interval.

| Parameter | Median | SD | InfCI | SupCI |
|---|---|---|---|---|
| $\sigma_P^2$ | 0.014 | 0.005 | 0.008 | 0.027 |
| $\sigma_G^2$ | 0.011 | 0.001 | 0.010 | 0.013 |
| $\sigma_{Inter}^2$ | 0.005 | 0.001 | 0.003 | 0.007 |
| $\sigma_S^2$ | 0.123 | 0.584 | 0.030 | 1.532 |
| $\sigma_B^2$ | 0.002 | 0.001 | 0.001 | 0.003 |
| $\sigma^2$ | 0.095 | 0.001 | 0.094 | 0.097 |
| $\beta_0$ | 6.229 | 0.226 | 5.777 | 6.686 |
| $\beta_{age}$ | 0.598 | 0.002 | 0.593 | 0.603 |
| $\beta_{age2}$ | -0.150 | 0.002 | -0.155 | -0.145 |

**Table S38:** Parameter estimates from model *M2* fitted on the P2 partition. SD corresponds to the standard deviation and InfCI and SupCI correspond to the lower and upper bounds of the 0.95 credible interval.

##### 6.2.2 Predictive *models M7* and *M8*: combining climatic and genomic drivers

| Term | Median | SD | InfCI | SupCI |
|---|---|---|---|---|
| $\sigma^2_B$ | 0.002 | 0.001 | 0.001 | 0.004 |
| $\sigma^2_{g_j}$ | 0.027 | 0.076 | 0.008 | 0.195 |
| $\sigma^2_S$ | 0.157 | 0.837 | 0.036 | 1.926 |
| $\sigma^2_{\beta_{max.temp,s}}$ | 0.001 | 0.004 | 0.000 | 0.007 |
| $\sigma^2_{\beta_{min.pre,s}}$ | 0.001 | 0.004 | 0.000 | 0.007 |
| $\sigma^2_{\beta_{gPEA,s}}$ | 0.019 | 0.055 | 0.005 | 0.136 |
| $\sigma^2$ | 0.103 | 0.001 | 0.101 | 0.104 |
| $\beta_0$ | 6.310 | 0.345 | 5.624 | 7.068 |
| $\beta_{age}$ | 0.600 | 0.003 | 0.595 | 0.605 |
| $\beta_{age2}$ | -0.151 | 0.002 | -0.156 | -0.147 |

**Table S39:** Parameter estimates from model *M7* fitted on the P2 partition. SD corresponds to the standard deviation and InfCI and SupCI correspond to the lower and upper bounds of the 0.95 credible interval.

| Term | Median | SD | InfCI | SupCI |
|---|---|---|---|---|
| $\sigma^2_B$ | 0.002 | 0.001 | 0.001 | 0.003 |
| $\sigma^2_{g_j}$ | 0.032 | 0.075 | 0.010 | 0.224 |
| $\sigma^2_S$ | 0.190 | 1.195 | 0.041 | 2.366 |
| $\sigma^2_{\beta_{max.temp,s}}$ | 0.003 | 0.006 | 0.001 | 0.019 |
| $\sigma^2_{\beta_{min.pre,s}}$ | 0.002 | 0.004 | 0.000 | 0.014 |
| $\sigma^2_{\beta_{rPEA,s}}$ | 0.032 | 0.085 | 0.010 | 0.241 |
| $\sigma^2$ | 0.101 | 0.001 | 0.099 | 0.103 |
| $\beta_0$ | 6.337 | 0.366 | 5.602 | 7.106 |
| $\beta_{age}$ | 0.600 | 0.003 | 0.595 | 0.605 |
| $\beta_{age2}$ | -0.151 | 0.002 | -0.156 | -0.147 |

**Table S40:** Parameter estimates from model *M8* fitted on the P2 partition. SD corresponds to the standard deviation and InfCI and SupCI correspond to the lower and upper bounds of the 0.95 credible interval.

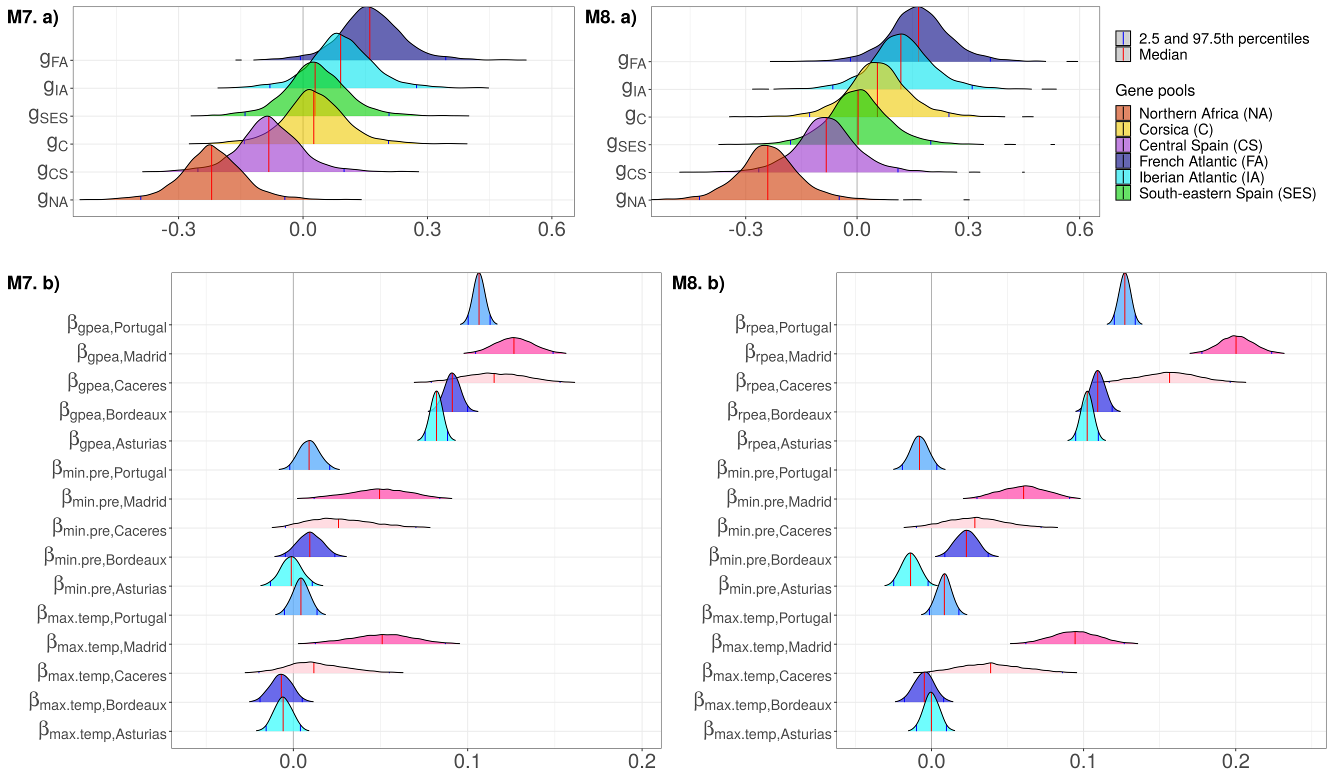

**Figure S17.** Posterior distributions of parameters from models *M7* and *M8* fit on P2. In panels a), the parameter estimates $g_j$ correspond to the effect of the gene pools $j$. In panels b), the parameter estimates correspond to the site-specific effects of the PEAs ($\beta_{gPEA,s}$ for the global PEAs in *M7* and $\beta_{rPEA,s}$ for the regional PEAs in *M8*), the minimum precipitation during the driest month ($\beta_{min.pre,s}$) and the maximum temperature of the warmest month ($\beta_{max.temp,s}$). Parameters specific to the same test site are colored the same to aid visualization.

##### 6.2.3 Predictive *models M9* to *M12*: including separately climatic and genomic drivers

| Parameter | Median | SD | InfCI | SupCI |
|---|---|---|---|---|
| $\sigma^2_{g_j}$ | 0.027 | 0.083 | 0.008 | 0.199 |
| $\sigma^2_S$ | 0.108 | 0.622 | 0.027 | 1.019 |
| $\sigma^2_B$ | 0.002 | 0.001 | 0.001 | 0.004 |
| $\sigma^2$ | 0.112 | 0.001 | 0.110 | 0.114 |
| $\beta_0$ | 6.230 | 0.217 | 5.802 | 6.647 |
| $\beta_{age}$ | 0.602 | 0.003 | 0.597 | 0.608 |
| $\beta_{age2}$ | -0.153 | 0.003 | -0.158 | -0.148 |

**Table S41:** Parameter estimates from model *M9* fitted on the P2 partition. SD corresponds to the standard deviation and InfCI and SupCI correspond to the lower and upper bounds of the 0.95 credible interval.

| Parameter | Median | SD | InfCI | SupCI |
|---|---|---|---|---|
| $\sigma^2_B$ | 0.002 | 0.001 | 0.001 | 0.004 |
| $\sigma^2_S$ | 0.183 | 1.789 | 0.036 | 3.006 |
| $\sigma^2_{\beta_{max.temp,s}}$ | 0.005 | 0.011 | 0.002 | 0.035 |
| $\sigma^2_{\beta_{min.pre,s}}$ | 0.006 | 0.018 | 0.002 | 0.041 |
| $\sigma^2$ | 0.113 | 0.001 | 0.111 | 0.115 |
| $\beta_0$ | 6.239 | 0.456 | 5.194 | 7.075 |
| $\beta_{age}$ | 0.602 | 0.003 | 0.597 | 0.607 |
| $\beta_{age2}$ | -0.153 | 0.003 | -0.158 | -0.148 |

**Table S42:** Parameter estimates from model *M10* fitted on the P2 partition. SD corresponds to the standard deviation and InfCI and SupCI correspond to the lower and upper bounds of the 0.95 credible interval.

| Term | Median | SD | InfCI | SupCI |
|---|---|---|---|---|
| $\sigma^2_B$ | 0.002 | 0.001 | 0.001 | 0.004 |
| $\sigma^2_S$ | 0.265 | 2.482 | 0.038 | 5.597 |
| $\sigma^2_{\beta_{gPEA,s}}$ | 0.013 | 0.031 | 0.004 | 0.085 |
| $\sigma^2$ | 0.115 | 0.001 | 0.113 | 0.117 |
| $\beta_0$ | 6.374 | 0.677 | 5.164 | 7.956 |
| $\beta_{age}$ | 0.600 | 0.003 | 0.594 | 0.605 |
| $\beta_{age2}$ | -0.151 | 0.003 | -0.156 | -0.146 |

**Table S43:** Parameter estimates from model *M11* fitted on the P2 partition. SD corresponds to the standard deviation and InfCI and SupCI correspond to the lower and upper bounds of the 0.95 credible interval.

| Term | Median | SD | InfCI | SupCI |
|---|---|---|---|---|
| $\sigma^2_B$ | 0.002 | 0.001 | 0.001 | 0.003 |
| $\sigma^2_S$ | 0.332 | 2.743 | 0.042 | 4.656 |
| $\sigma^2_{\beta_{rPEA,s}}$ | 0.026 | 0.083 | 0.008 | 0.177 |
| $\sigma^2$ | 0.112 | 0.001 | 0.110 | 0.114 |
| $\beta_0$ | 6.556 | 0.532 | 5.620 | 7.714 |
| $\beta_{age}$ | 0.599 | 0.003 | 0.594 | 0.605 |
| $\beta_{age2}$ | -0.151 | 0.003 | -0.156 | -0.146 |

**Table S44:** Parameter estimates from model *M12* fitted on the P1 partition. SD corresponds to the standard deviation and InfCI and SupCI correspond to the lower and upper bounds of the 0.95 credible interval.

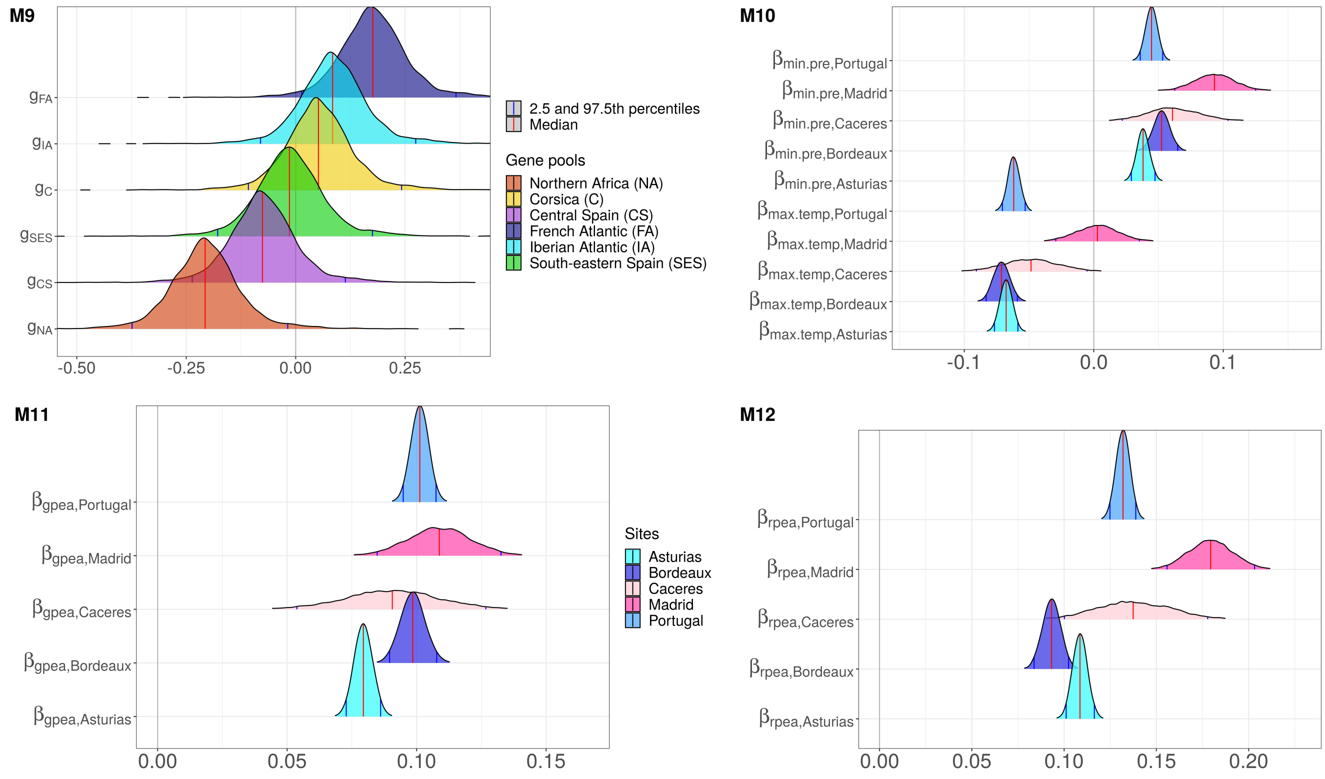

**Figure S18.** Posterior distributions of parameters from models *M9* and *M12* fitted on P2. For *M9*, the parameter estimates $g_j$ correspond to the effect of the gene pools $j$. For *M10*, the parameter estimates correspond to the site-specific effects of the minimum precipitation during the driest month ($\beta_{min.pre,s}$) and the maximum temperature of the warmest month ($\beta_{max.temp,s}$). For *M11*, the parameter estimates correspond to the site-specific effect of the gPEAs ($\beta_{gPEA,s}$). For *M12*, parameter estimates correspond to the site-specific effects of the rPEAs ($\beta_{rPEA,s}$).

#### 6.3 P3 partition (non-random split of the provenances)

Evaluation of model performance on new provenances was replicated on six other provenances to assess the robustness of the results. In the P3 partition, the new provenances were not totally randomly selected to ensure that each under-represented gene pool in our study was represented by at least one provenance. Thus, one provenance was randomly selected from the two provenances belonging mainly to the northern Africa gene pool. The same was done for the gene pools from south-eastern Spain and Corsica. The last three provenances were randomly selected from the three remaining gene pools.

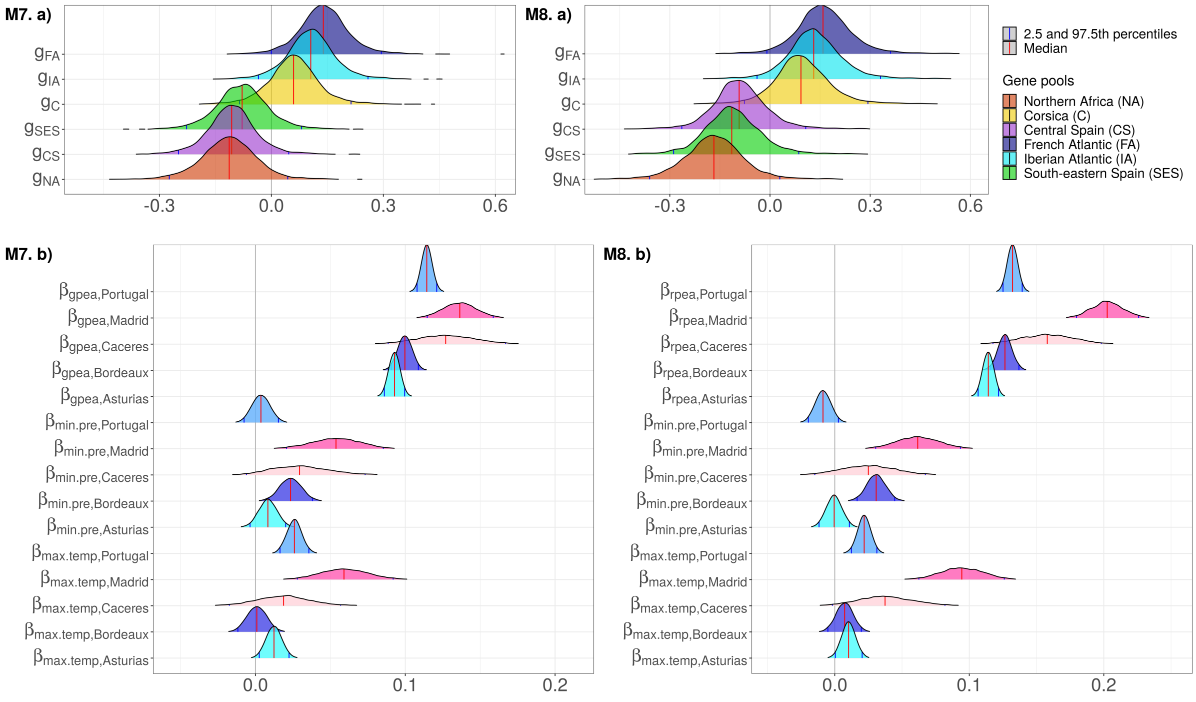

**Figure S19.** Posterior distributions of parameters from models *M7* and *M8* fit on P3. In panels a), the parameter estimates $g_j$ correspond to the effect of the gene pools $j$. In panels b), the parameter estimates correspond to the site-specific effects of gPEAs in *M7* ($\beta_{gPEA,s}$, rPEAs in *M8* $\beta_{rPEA,s}$, the minimum precipitation during the driest month ($\beta_{min.pre,s}$) and the maximum temperature of the warmest month ($\beta_{max.temp,s}$). Parameters specific to the same site are colored the same to aid visualization.

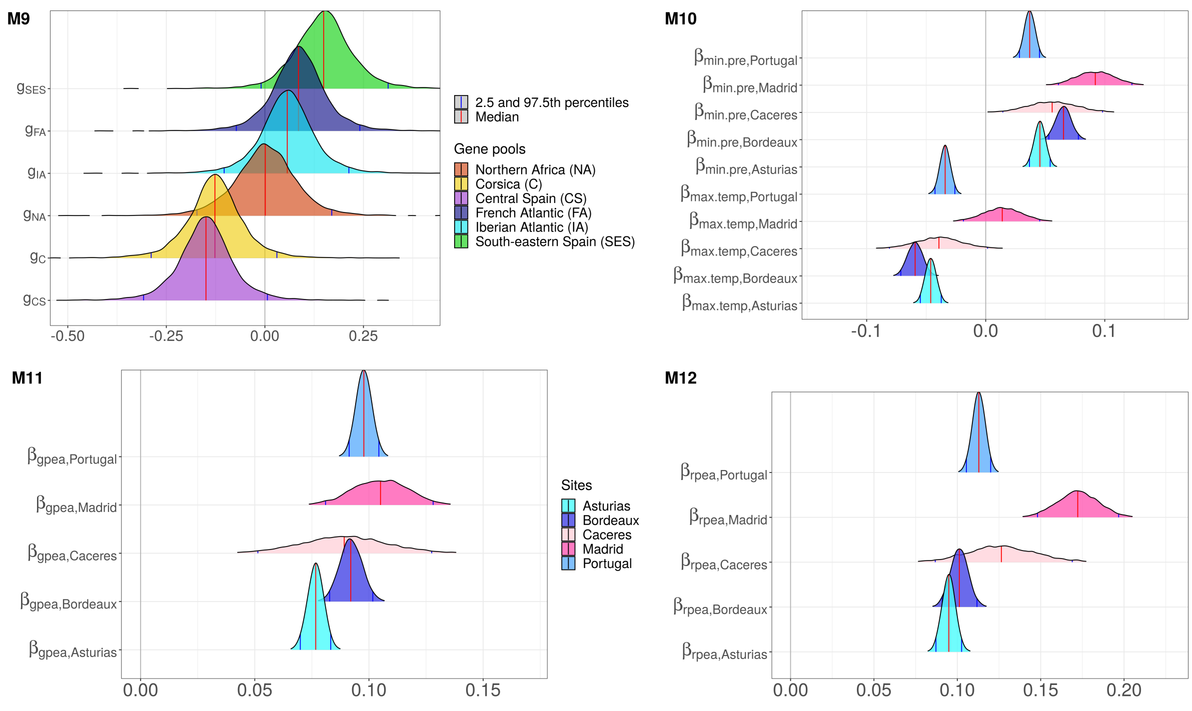

**Figure S20.** Posterior distributions of parameters from models *M9* and *M12* fit on P3. For *M9*, the parameter estimates $g_j$ correspond to the effect of the gene pools $j$. For *M10*, the parameter estimates correspond to the site-specific effects of the minimum precipitation during the driest month ($\beta_{min.pre,s}$) and the maximum temperature of the warmest month ($\beta_{max.temp,s}$). For *M11*, the parameter estimates correspond to the site-specific effect of the gPEAs ($\beta_{gPEA,s}$). For *M12*, parameter estimates correspond to the site-specific effects of the rPEAs ($\beta_{rPEA,s}$).

#### 6.4 Interpretation of the PEAs coefficients

Let's take model *M12* as an example. Here is the equation of *M12* (see equation 4.1.3 in the Supplementary Information):

$$\log(h_{isbr}) \sim \mathcal{N}(\mathbf{X}\beta + S_s + B_{b(s)} + \beta_{rPEA,s} rPEA_{gr} \, , \, \sigma^2)$$

with $rPEA_{gr}$ is the scaled explanatory variable (i.e. the counts of regionally-selected positive-effect alleles). Let's call $r\tilde{PEA}_{gr}$ the explanatory variable before being scaled, that is $rPEA_{gr} = (r\tilde{PEA}_{gr} - \mu_{r\tilde{PEA}_{gr}})/\sigma_{r\tilde{PEA}_{gr}}$, where $\mu_{r\tilde{PEA}_{gr}}$ is the mean of $r\tilde{PEA}_{gr}$ and $\sigma_{r\tilde{PEA}_{gr}}$ is its standard deviation.

We want to calculate the percent of change in height associated with a one-unit increase in $rPEA_{gr}$, that is a one-standard deviation increase in $r\tilde{PEA}_{gr}$. For that, we call $h_{new}$ the value of $h_{isbr}$ after increasing $rPEA_{gr}$ by one unit, and we have:

$$\begin{aligned} \log(h_{new}) &= \log(\mathbf{X}\beta + S_s + B_{b(s)} + \beta_{rPEA,s}(rPEA_{gr} + 1) \\ &= \log(h_{isbr}) + \beta_{rPEA,s} \end{aligned}$$

Therefore:

$$\log(h_{new}) - \log(h_{isbr}) = \beta_{rPEA,s}$$

$$\frac{h_{new}}{h_{isbr}} = \exp(\beta_{rPEA,s})$$

$$100 \times \left(\frac{h_{new}}{h_{isbr}} - 1\right) = 100 \times (\exp(\beta_{rPEA,s}) - 1)$$

$$100 \times \left(\frac{h_{new} - h_{isbr}}{h_{isbr}}\right) = 100 \times (\exp(\beta_{rPEA,s}) - 1)$$

is the percent change in $h_{isbr}$ associated with a one-unit increase in $rPEA_{gr}$ (that is, a one-standard deviation increase in $r\tilde{PEA}_{gr}$). For instance, a one-standard deviation increase in the counts of rPEAs is associated, on average, with $100 \times (\exp(0.174) - 1) = 19\%$ change in height in Madrid, with $100 \times (\exp(0.120) - 1) = 12.7\%$ change in height in Cáceres, with $100 \times (\exp(0.092) - 1) = 9.6\%$ change in height in Bordeaux, with $100 \times (\exp(0.099) - 1) = 10.4\%$ change in height in Asturias and with $100 \times (\exp(0.122) - 1) = 13.0\%$ change in height in Portugal.

### 7 $Q_{ST} - F_{ST}$ analysis

To determine whether height growth shows footprints of adaptive differentiation, we performed a $Q_{ST} - F_{ST}$ analysis. We used the global $F_{ST}$ estimate calculated in de Miguel et al. (2020) on the same data as our study (i.e. the 5,165 SNPs from the Illumina Infinium SNP array). de Miguel et al. (2020) used the *diveRsity* R package and 1,000 bootstrap iterations across loci to estimate the 95% confidence interval of the global $F_{ST}$. They obtained a $F_{ST}$ of 0.112 (95% confidence interval: 0.090 - 0.141).

To calculate the $Q_{ST}$, we used the following formula from Spitze (1993):

$$Q_{ST} = \frac{\sigma_P^2}{\sigma_P^2 + 2\sigma_G^2}$$

where $\sigma_P^2$ is the variance among provenances, and $\sigma_G^2$ is the variance among clones (.i.e. genotypes) within provenances.

The median estimate of the $Q_{ST}$ was 0.358 (95% confidence interval: 0.251-0.506). Quantitative ($Q_{ST}$) and molecular ($F_{ST}$) genetic differentiation among provenances were considered significantly different as their posterior distributions had non-overlapping 95% confidence intervals, which therefore suggests that there is adaptive differentiation in height growth in our study.

### 8 Distribution of heights in Cáceres and Madrid.

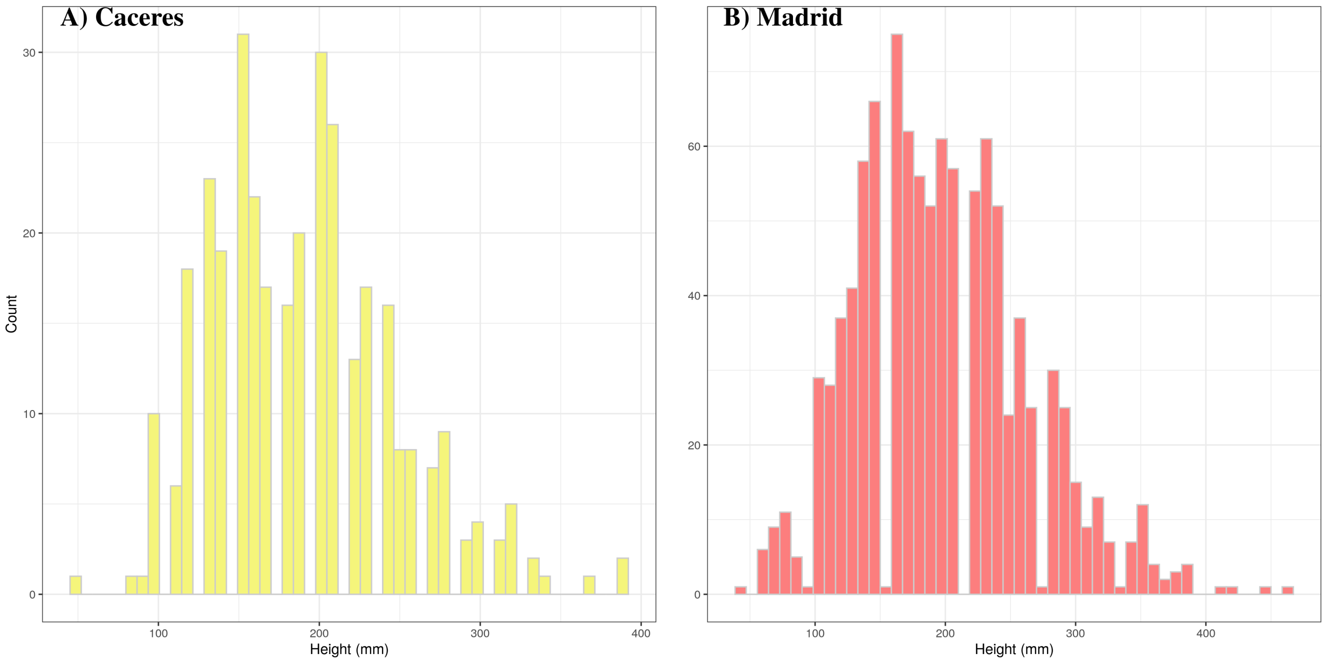

**Figure S21.** Distribution of height measurements (in mm) in Cáceres and Madrid.
